# Integrating pharmacogenomics data-driven prediction with bulk and single-cell RNAseq to demonstrate the efficacy of an NAMPT inhibitor against aggressive, taxane-resistant, and stem-like cells in lethal prostate cancer

**DOI:** 10.1101/2022.07.20.500378

**Authors:** Suman Mazumder, Taraswi Mitra Ghosh, Ujjal K. Mukherjee, Sayak Chakravarti, Farshad Amiri, Farnaz Hemmati, Panagiotis Mistriotis, Isra Elhussin, Ahmad-bin Salam, Windy Dean-Colomb, Feng Li, Clayton Yates, Robert D. Arnold, Amit K. Mitra

**Affiliations:** Department of Drug Discovery and Development, Harrison College of Pharmacy, Auburn University, AL; Center for Pharmacogenomics and Single-Cell Omics (AUPharmGx), Harrison College of Pharmacy, Auburn University, Auburn, AL; Department of Business Administration, University of Illinois at Urbana-Champaign, Champaign, IL; Biomedical and Translational Sciences, Carle Illinois College of Medicine, University of Illinois Urbana- Champaign; Department of Chemical Engineering, Samuel Ginn College of Engineering, Auburn University, AL; Department of Biology and Canter for Cancer Research, Tuskegee University, Tuskegee, AL; Piedmont Hospital, Newnan, GA; Department of Pathology, University of Alabama at Birmingham School of Medicine, Birmingham, AL; UAB O’Neal Comprehensive Cancer, University of Alabama at Birmingham School of Medicine, Birmingham, AL

**Keywords:** secDrug, FK866, taxane, prostate cancer, synergism, RNAseq, scRNAseq, DEGs, IPA

## Abstract

Metastatic prostate cancer is the second leading cause of cancer deaths in US men. Resistance to standard medical castration and secondary taxane-based chemotherapy is nearly universal. Further, presence of cancer stem-like cells (EMT/epithelial to mesenchymal transdifferentiation) and neuroendocrine PCa (NEPC) subtypes significantly contribute to *aggressive/advanced/lethal variants of PCa (AVPC)*.

In this study, first we used single-cell RNA sequencing (scRNAseq) analysis to demonstrate that AR^low^ PCa cells in metastatic prostate cancer, including castration-sensitive tumors, harbored signatures of EMT, and ‘cancer stemness’. Next, we introduced a novel pharmacogenomics data-driven computational approach and identified several potential agents that can be re-purposed as novel secondary drugs (“secDrugs”) to treat advance variants of Prostate cancer. Using scRNAseq as a biomarker-based drug screen, we demonstrated that a majority of the single-cell subclones in mCRPC and mCSPC cell lines also showed significantly high expression of the NAMPT pathway genes, indicating that the secDrug FK866, which targets NAMPT, is potentially effective against drug-resistant and stem-cell-like subpopulation cluster. Next, we showed significant *in vitro* cytotoxicity of FK866 as single-agent and in combination with the taxanes or Enzalutamide against models of clinically-advanced PCa. We performed bulk- and single-cell RNAseq to identify several pathways underlining FK866 mechanism of action and found that in addition to NAMPT inhibition, FK866 regulates tumor metastasis, cell migration, invasion, DNA repair machinery, redox homeostasis, autophagy, as well as cancer stemness–related genes HES1 and CD44. Further, we performed a microfluidic chip-based cell migration assay that demonstrated that FK866 reduces cancer cell invasion and motility, indicating abrogation of metastasis. Finally, using multiple PCa patient datasets, we showed that FK866 is potentially capable of reversing expression of several genes associated with biochemical recurrence and inter-ethnic differences, including IFITM3 and LTB4R.

Thus, using FK866 as a proof-of-concept drug, we introduced a novel, universally applicable preclinical drug development pipeline to circumvent subclonal aggressiveness, drug resistance, and stemness in lethal PCa.

## 1. INTRODUCTION

Prostate cancer (PCa) is the second leading cause of non-cutaneous cancer-related deaths in the US (www.cancer.org). The androgen signaling pathway plays a crucial role in PCa development(Shafi *et al*., 2013). Therefore, the standard treatment options for PCa are radical prostatectomy (RP) or radiation therapy with androgen-deprivation therapy (ADT)(Hamdy *et al*., 2016). Most early-stage PCa patients (castration-sensitive or CSPC) treated with ADT show good initial response with a high 5-year survival rate (www.cancer.org, 2019). However, a vast majority of these men eventually become unresponsive towards hormone therapy, and despite low levels of androgen, the disease progresses with continuously rising Prostate Serum Antigen (PSA), eventually developing more aggressive forms called Castration-resistant prostate cancer (CRPC)(Kapoor *et al*., 2016; Scher *et al*., 2014; Theodoros Karantanos, Paul G. Corn, 2013; Wadosky and Koochekpour, 2016). Metastatic castration-resistant prostate cancer (mCRPC) is the clinically most advanced and lethal disease state with signs of metastasis to distant organs like brain, bone, lung, lymph node and median survival of less than 3 years (5-year median survival rate of 31%)(Scher *et al*., 2015). Although next-generation AR-targeting chemotherapeutic treatments like abiraterone plus prednisone (AA/P) or enzalutamide (ENZ), and combination with taxanes (Docetaxel/DTX or cabazitaxel/CBZ), increase survival rate slightly, eventual development of resistance (acquired resistance) is nearly universal where progression-free survival approaches ∼0% in 3 years, often with severe side effects(Cornford *et al*., 2017; De Bono *et al*., 2010; Galletti *et al*., 2017; Petrylak *et al*., 2004; Saad and Miller, 2014; Sartor, 2011; Scher *et al*., 2012, 2015; Tannock *et al*., 2004; Y. Wang *et al*., 2021). Chemotherapy options become limited once patients fail DTX therapy. Further, neuroendocrine PCa or NEPC (also known as small cell carcinoma) is an intrinsically resistant, poorly differentiated aggressive variant of PCa that lacks AR expression(Conteduca *et al*., 2019; Vashchenko and Abrahamsson, 2005).

In addition, several groups, including ours, have shown that the presence of cancer stem-like cells (CSCs) like side populations (SPs) and CD133^+^ cells with self-renewal and differentiation (acquisition of mesenchymal phenotype or epithelial to mesenchymal transdifferentiation/EMT) capacities significantly contribute to tumor aggressiveness and the development of drug resistance(Contreras *et al*., 2020; P. Li *et al*., 2014; Zhou *et al*., 2011).

Drug development for these clinically most-aggressive and lethal variants of PCa (AVPC) thus poses a significant challenge with very few therapeutic successes.

Our laboratory has designed a pharmacogenomics data-driven computational pipeline (secDrug) that identifies novel secondary drugs (“secDrugs”) for the treatment of drug-resistant advanced-state cancers(Kumar *et al*., 2022).

In this study, we applied the secDrug algorithm to PCa models and identified several novel secondary drug candidates for the treatment of AVPC. Next, using single-cell RNA sequencing (scRNAseq), we demonstrated the presence of PCa subclones representing aggressive, TX-resistant, and cancer stem-like cells. Further, our scRNAseq data predicted that the secDrug, FK866 (a NAMPT inhibitor), is potentially effective against PCa subclones with enrichment of treatment-resistant and stem-like genes. We hypothesize that our predicted and pre-screened secDrugs would be helpful in curbing oncogenic progressions as single-agent or in combination with taxanes in AVPC through simultaneous inhibition of multiple oncogenic factors/pathways. Using *in vitro* model systems of treatment-refractory and treatment-emergent AVPC (representing mCRPC, NEPC, and EMT), we demonstrated that the FK866 not only showed efficacy as a single-agent but also enhanced the efficacy of the taxane drugs DTX and CBZ. Further, we performed a sophisticated microfluidic chip-based confined cell migration assay that recapitulates diverse micro-environmental cues encountered by cancer cells during locomotion (e.g., the dimensionality of pores and 3D longitudinal, channel-like tracks) to investigate the effect of FK866-based regimens on cancer cell invasion, motility, and metastasis. Finally, we demonstrated the impact of FK866 in eroding ‘stem-like’ subpopulations (including SPs, quiescent/dormant cells, and ALDH1+cells).

Earlier studies have shown FK866 inhibits the growth of PCa engineered tumors by interference with the energy metabolism(Bowlby *et al*., 2012; Sauer *et al*., 2021). However, the clinical benefit and comprehensive mechanism of action (MOA) for FK866 have not been uncovered fully. Therefore, we performed pre- vs. post-treatment bulk and single-cell tumor RNAseq to identify differentially expressed genes (DEGs) and potential molecular pathways associated with the FK866 mechanism of action in AVPC at the tumor and subclonal levels. Finally, using comparative analysis of whole-genome transcriptomics data between clinically sensitive and resistant PCa patients, we demonstrated that FK866 has the potential to be clinically effective based on the reverse matching of GEP signatures and top dysregulated pathways.

Hence, using an innovative approach that integrates single-cell -omics technologies, microfluidics, and tumor mRNA sequencing with *In vitro* studies and patient data based validation, we conclude that FK866 has the potential to improve the clinical outcome in AVPC chemotherapy by enhancing the therapeutic efficacy and abrogating the possibilities of development of bulk and subclonal drug resistance. Such an evidence-based approach promises to minimize the chances of trial failures and improve the probability of clinical success.

## 2. RESULTS

### scRNAseq showed AR^low^ PCa cells with signatures of Epithelial-mesenchymal transition (EMT) and cancer ‘stemness’

**Figure 1A** displays t-SNE clusters generated from baseline (untreated) scRNAseq data in mCSPC and mCRPC cell lines. Each dot represents a single cell. Further, the AR status of each cell is represented in **Figure 1B**. Epithelial-mesenchymal transitions have been mechanistically linked with the generation and maintenance of stem-like cell populations during tumorigenesis. PCa cells that have undergone EMT are phenotypically and genomically similar to stem cells. For example, Vimentin is a well-characterized filament protein that is highly expressed in mesenchymal cells. Thus, enhanced levels of Vimentin and downregulation of E-cadherin served as markers for identifying cells that have undergone EMT. **Figures 1C-E** demonstrates that the AR^low^ cells (primarily belonging to the mCRPC subtype) show higher expression of several mesenchymal gene signatures involved in Epithelial-mesenchymal transition with NEPC phenotype, including Vimentin (**Figure 1C**); N-cadherin (CDH2), Fibronectin (FN1), S100A4, Snail (SNAI1), Slug (SNAI2) (**Figure 1D**); and other major EMT markers CDH11, TWIST1, ZEB1 (**Figure 1E**). Further, **Figures 1F-G** show upregulation of cancer stemness-related markers Urokinase-type plasminogen activator (PLAU), Urokinase-type plasminogen activator receptor (PLAUR), and CD44, primarily in mCRPC cells.

**Figure 1.**
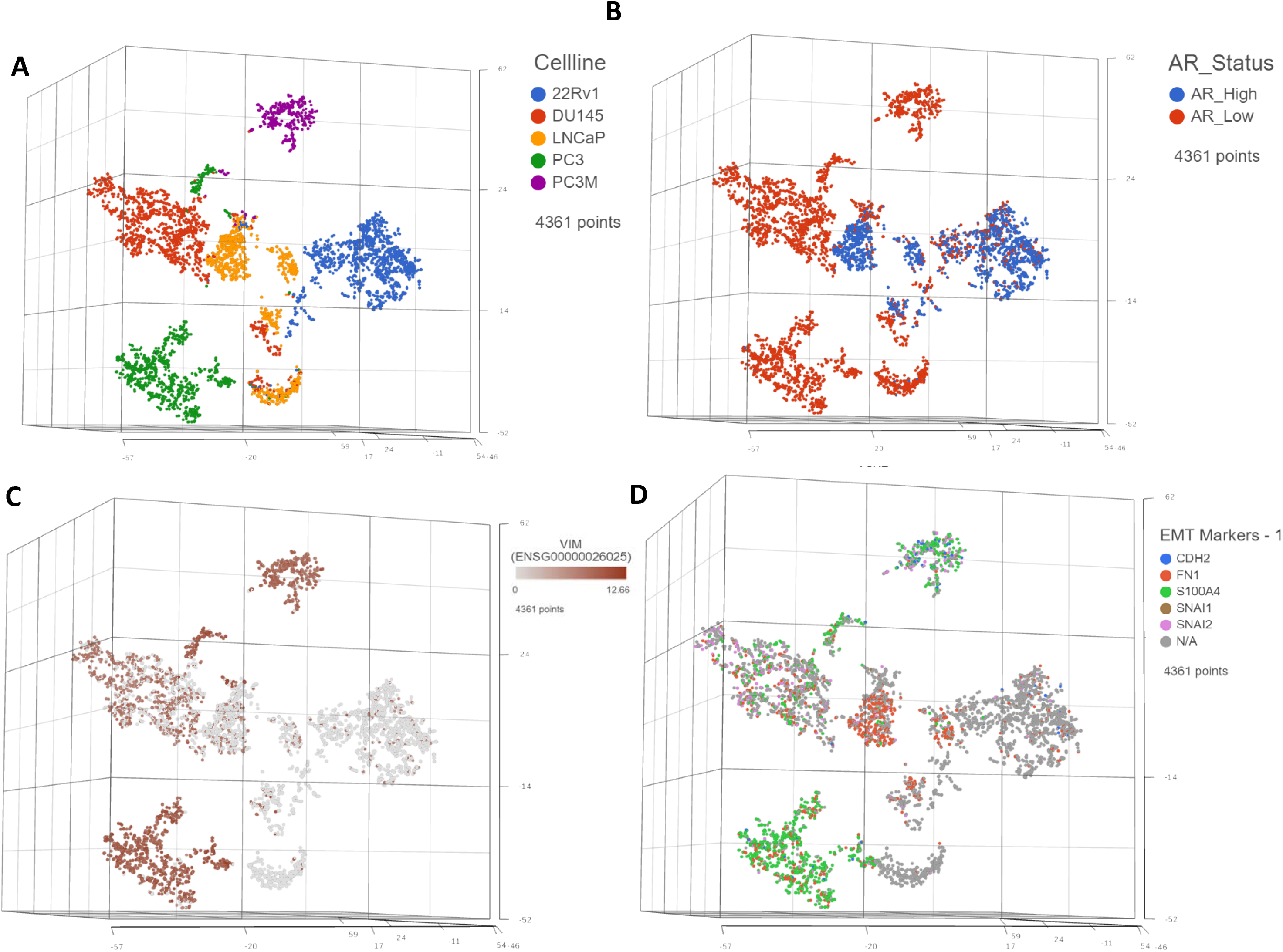
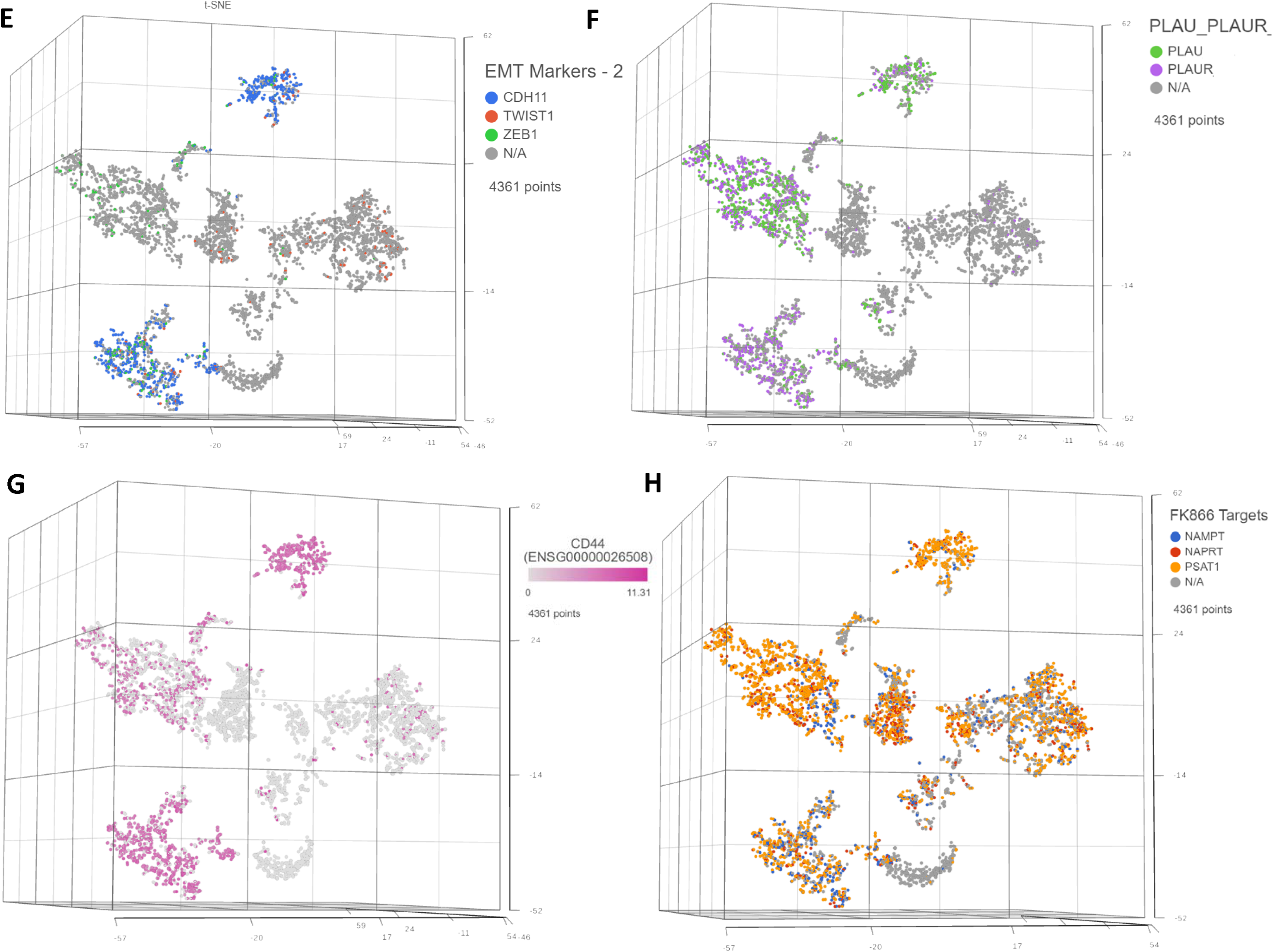
Single-cell transcriptomics identifies signatures of epithelial to mesenchymal transition (EMT) and stemness in metastatic prostate cancer cells. Single-cell RNA sequencing using the Droplet sequencing method (10X Genomics) was performed on the PCa cell lines 22Rv1, LnCAP, DU145, PC3, and PC3M. t-distributed stochastic neighbor embedding (tSNE) plots showing the comparison between the single-cell clusters representing **A)** All cell lines; **B)** AR Status; Expression of mesenchymal markers involved in EMT transdifferentiation, including the **C)** Vimentin (VIM); **D)** N-Cadherin (CDH1), Fibronectin (FN1), S100A4, Snail (SNAI1), Slug (SNAI2); other major EMT markers **E)** CDH11, TWIST1, ZEB1. Expression of genes potentially involved in Cancer stemness **F)** Urokinase-type plasminogen activator (PLAU), and Urokinase-type plasminogen activator receptor (PLAUR); **G)** CD44; FK866 target pathway genes **H)**Nicotinamide phosphoribosyltransferase (NAMPT), Nicotinate Phosphoribosyltransferase (NAPRT), and Phosphoserine Aminotransferase 1 (PSAT1). Each dot represents a single cell. Contaminated (doublet) cells were not included.

### Interestingly, signatures of cancer stemness and EMT transdifferentiation were also observed in a subgroup of AR^low^ single cells within the mCSPC cell lines, 22Rv1, LnCAP

Next, we compared the single-cell gene expression markers between taxane-sensitive (DU145) and the clonally derived acquired taxane-resistant mCRPC cell line DUTXR (**Figure 2**). We observed upregulation of gene signatures association with mesenchymal transition (VIM and TGFB1) and downregulation of the epithelial marker epithelial cadherin/E-cadherin (CDH1) in the DUTXR cell line compared to DU145 (**Figure 2B-D**). Further, the taxane-resistant DUTXR also showed enrichment of biomarkers that play significant roles in cancer progression, development, and maintenance of cancer stemness (CD44; **Figure 2E-F**) and drug resistance (CDK1, CXCL8; **Figure 2G**), indicating probable involvement in mCRPC development and progression(L. Deng *et al*., 2019).

**Figure 2.**
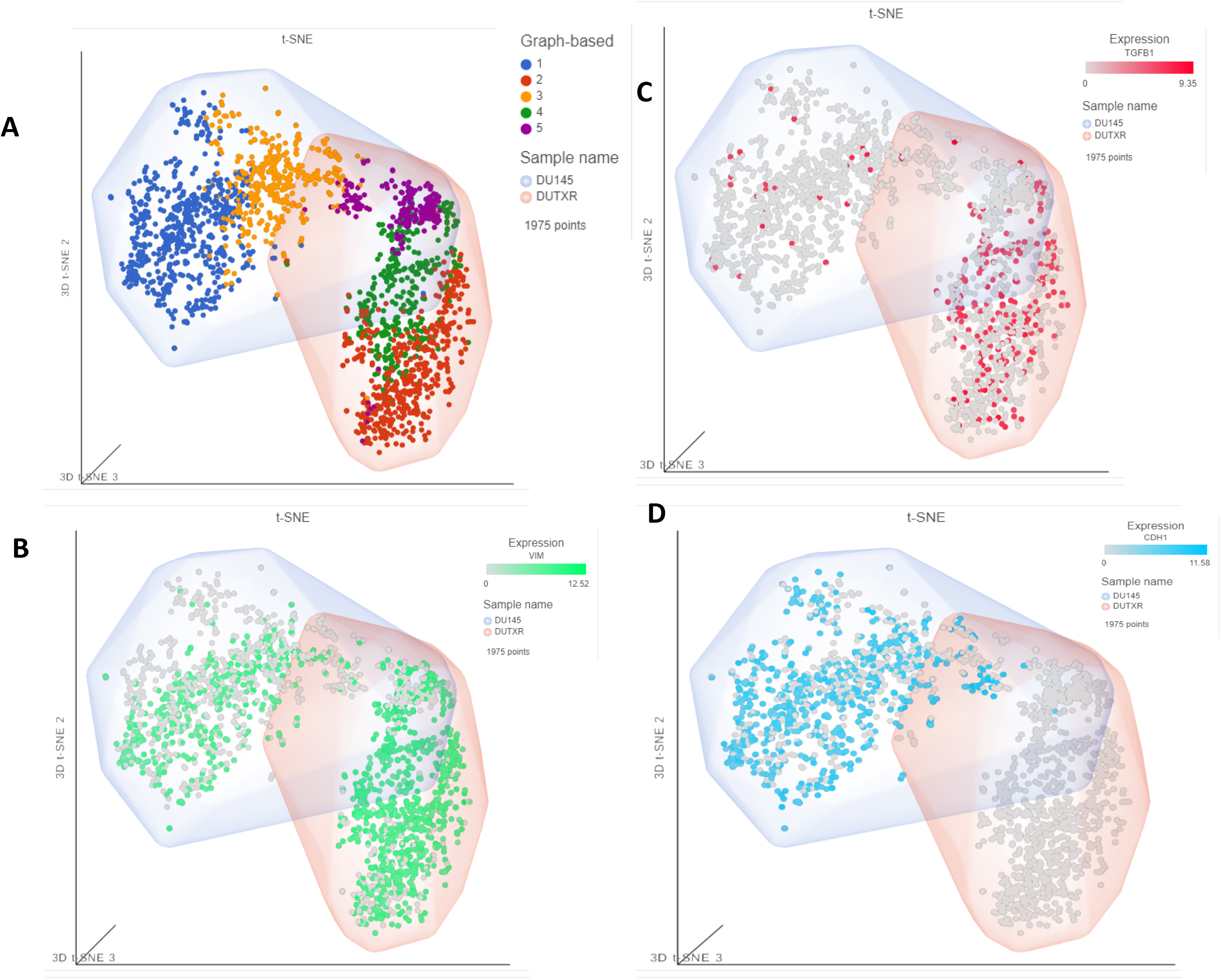
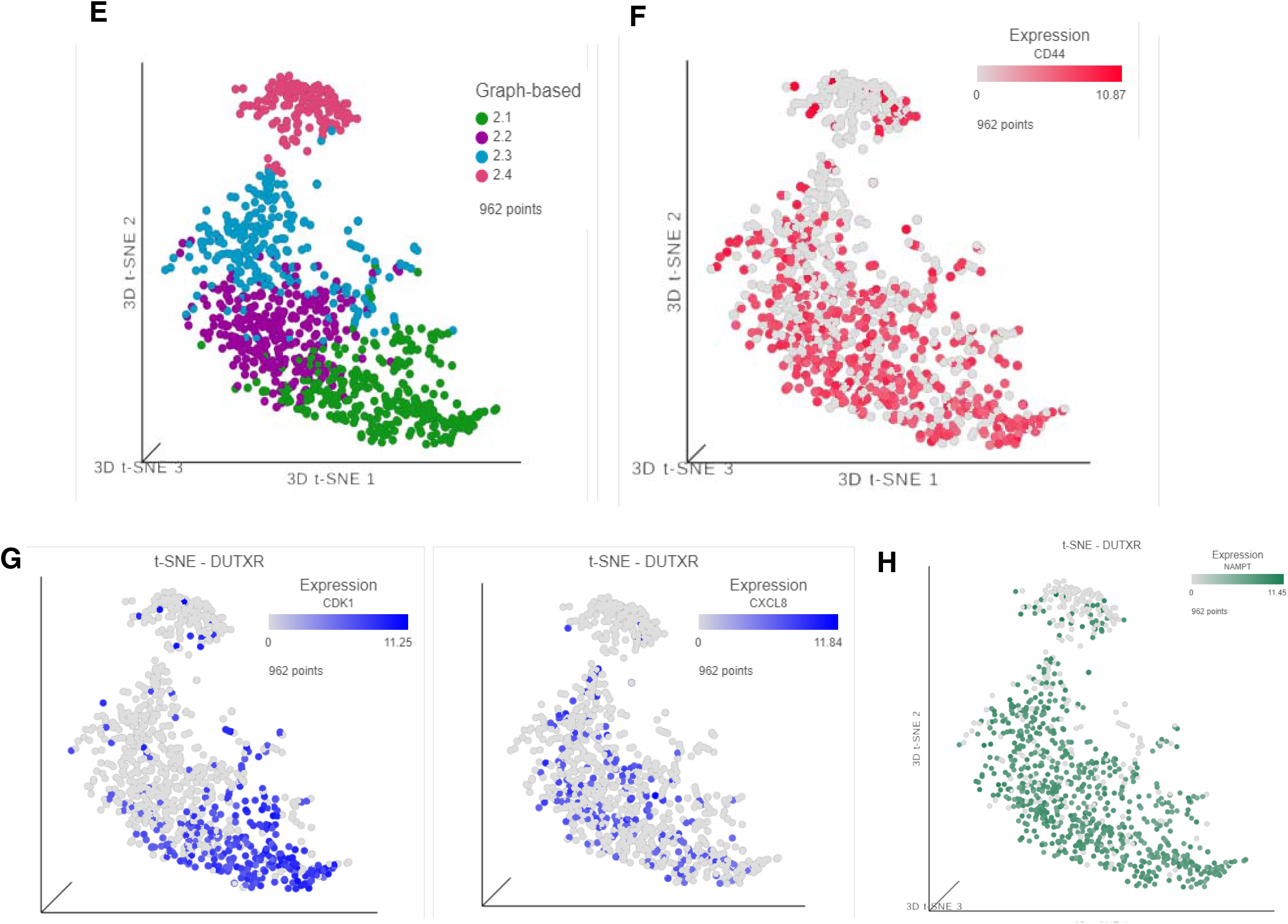
Comparison of single-cell RNAseq data between DU145 and DUTXR. **A)** All 4 t-SNE clusters, **B-D)** showing the EMT markers. **E)** scRNAseq data shows DUTXR cells representing enrichment of gene signatures of **F)** cancer stemness CD44^high^; **G)** Drug resistance (CDK1^high^; CXCL8 ^high^), and the **H)** FK866 target NAMPT ^high^. Each dot represents a single cell. Contaminated (doublet) cells were not included. Results show expression of the following B); TGF-B1; and E-Cadherin (CDH1).

### Pharmacogenomics data-driven algorithm and scRNAseq-based screening predicted FK866 as a top secDrug potentially effective against lethal PCa

Our pharmacogenomics data-driven *in silico* prediction algorithm (described in the *Methods* section) identified several potential agents that can be re-purposed as novel secondary drugs (“secDrugs”; **Table S1**) to treat DTX-resistant Prostate cancer when used as single-agent or in combination with the primary drug (Taxanes). These include FK866 (NAMPT inhibitor), TAK715 (<u>p38 MAPK inhibitor)</u>, YM155 (survivin inhibitor), MK-2206 (Akt1/Akt2/Akt3 inhibitor), LY317615 (PKCβ inhibitor), XAV939 (Wnt/β-catenin pathway inhibitor), RDEA119 (MEK1/2 inhibitor), and WZ3146 (mutant-selective irreversible inhibitor of EGFR (L858R)/EGFR (E746_A750)). Interestingly, using scRNAseq as a biomarker-based drug screen, we demonstrated that a majority of the single-cell subclones in mCRPC and mCSPC cell lines also showed significantly high expression of the NAMPT pathway genes, indicating that FK866, which targets NAMPT, is potentially effective against these taxane-resistant and stem-cell-like subpopulation clusters (**Figures 1H and 2H**).

### FK866 induced loss of viability in PCa cell lines as single-agent treatment and showed synergy with Taxanes and AR inhibitors

The effect of DTX, CBZ, and FK866 as single-agent administration on mCSPC, mCRPC cells lines, as well as the acquired taxane resistant mCRPC line DUTXR cells, were assessed by MTT assay at increasing drug concentrations treatment of each drug. *In vitro* single-agent cytotoxicity assay results are displayed in **Figure S1** as dose vs. %survival. FK866 shows time-dependent decreases in cell survival after 48hr of drug treatment in all the PCa cell lines. Furthermore, IC_50_ values of FK866 were inversely correlated with the IC_50_ values of taxanes, indicating possible synergy (Spearman r<-0.9; p=0.0167). Next, we evaluated the effect of increasing concentrations of DTX + FK866 combination treatment on the AR^lo^ mCRPC cell lines PC-3, DU145, the acquired taxane-resistant mCRPC lines DUTXR and PC3-TXR, the clonally-derived metastatic (PC3M), and the AR-independent (C42B) PCa lines, as well as the AR^high^ mCSPC lines LnCAP and 22Rv1 (**Figure 3A-D**). The dose-response curves for the drug combinations and CI values indicated high synergy, which was even more profound (CI between 0.2-0.37) in the TX-resistant lines (**Figure 3E**). Similar highly synergistic results were observed for the FK866+CBZ combination treatment as well were in the acquired taxane resistant mCRPC (DU145 resistant-DUTXR) cells (**Figure S2**).

**Figure 3.**
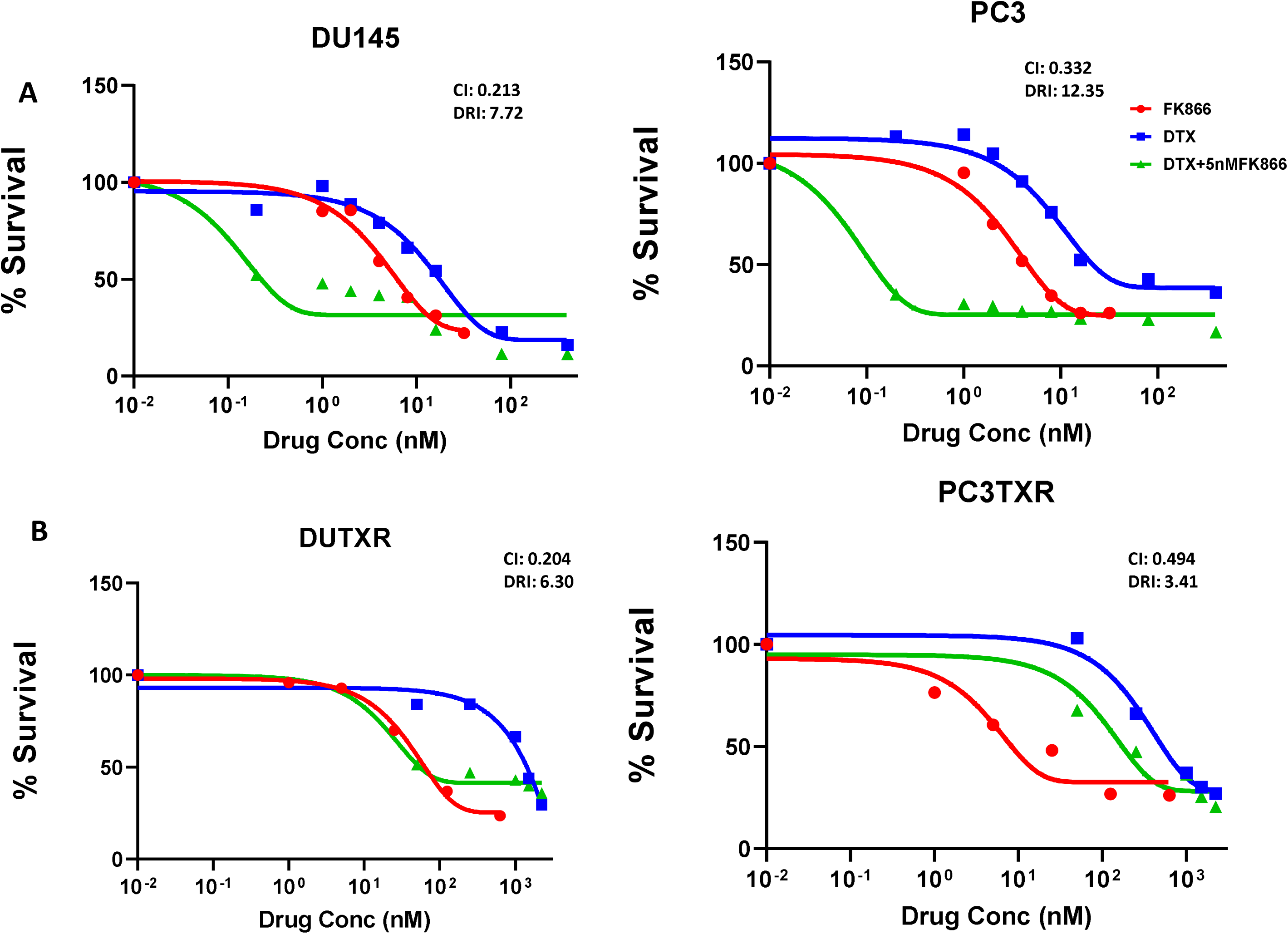
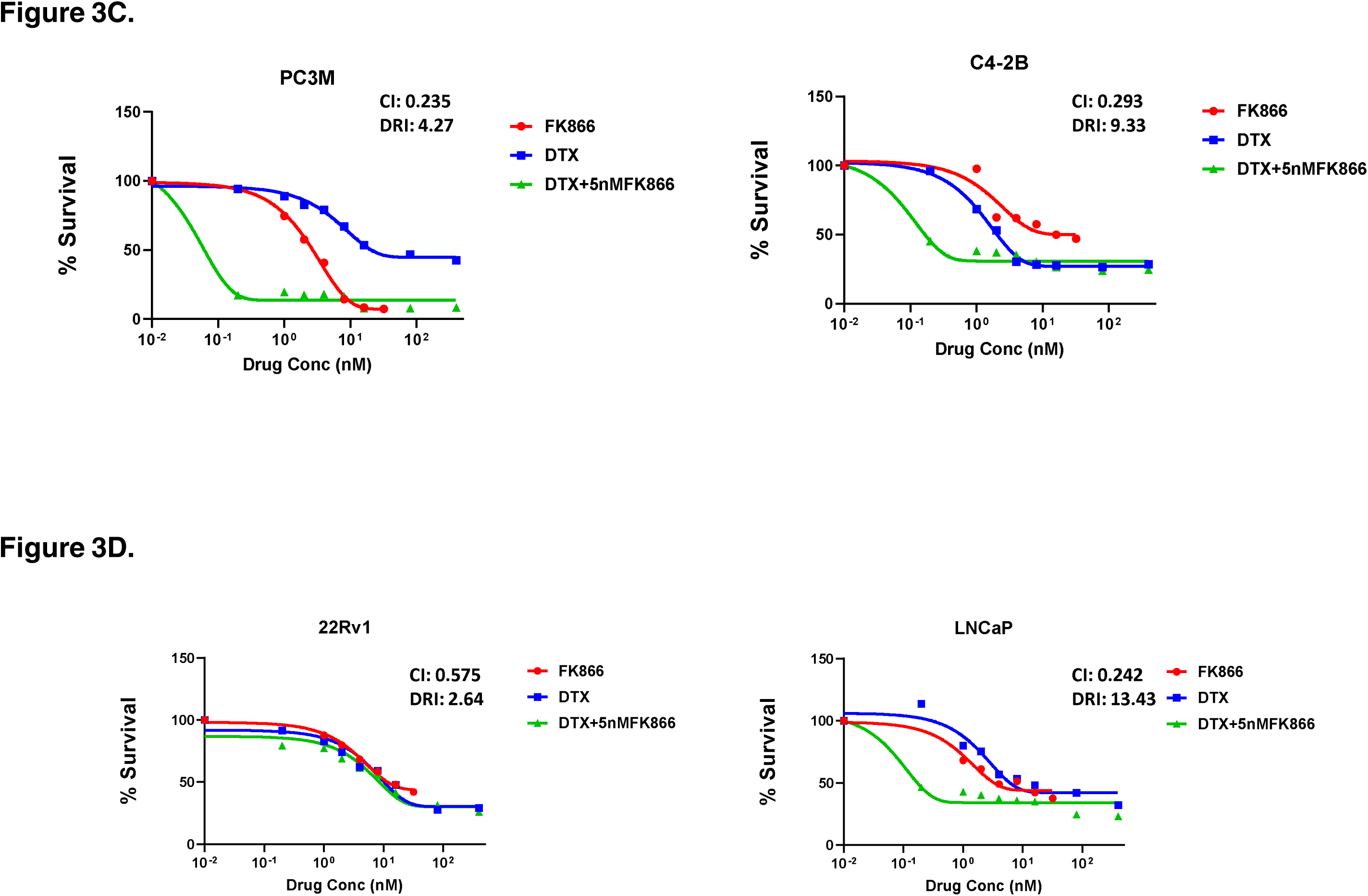
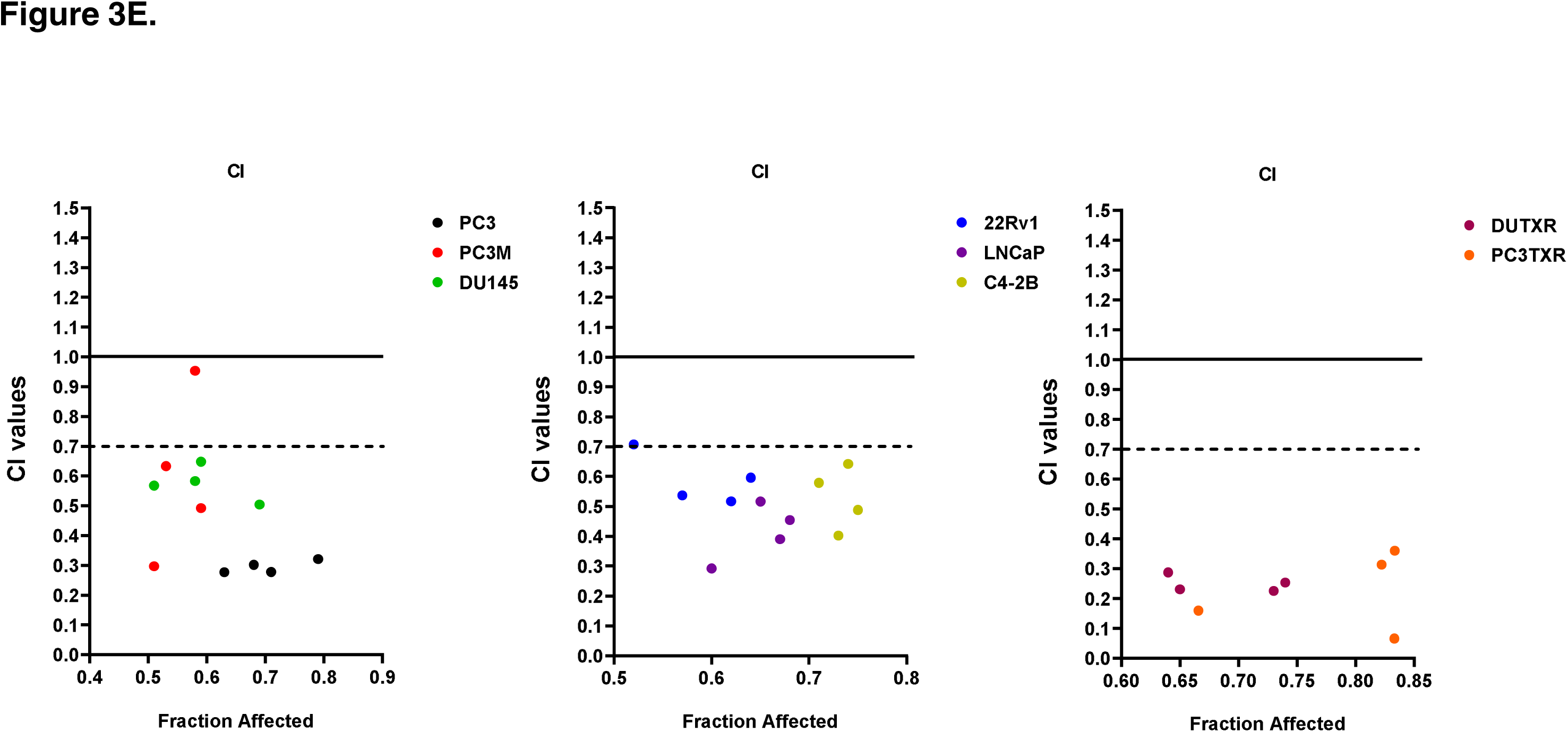
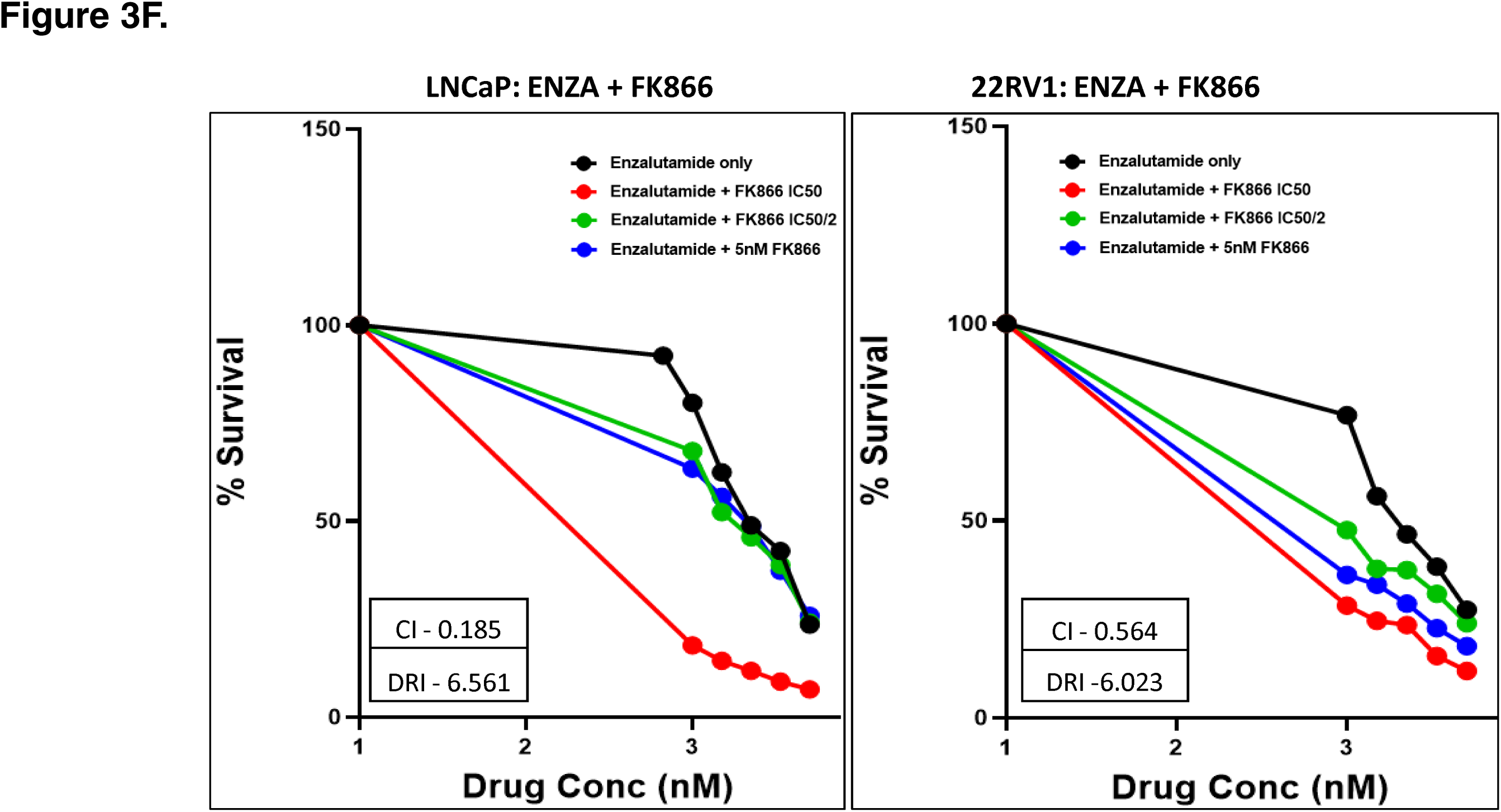
Dose-response curves representing *in vitro* cytotoxicity of the FK866 drug combination in metastatic PCa cell lines. **A-D)** *In vitro* cytotoxicity was assessed using MTT or 4, 5-Dimethylthiazol-2-yl)-2,5-diphenyltetrazolium bromide assay following treatment with single-agent DTX or single-agent FK866 at increasing concentrations or increasing concentrations of DTX in combination with fixed (5nM) concentration of FK866. **A)** AR^lo^ mCRPC cell lines PC-3, DU145, **B)** Acquired taxane-resistant mCRPC lines DUTXR and PC3-TXR, **C)** clonally-derived metastatic (PC3M), and the AR-independent (C42B) PCa lines; **D)** AR^high^ mCSPC lines LnCAP and 22Rv1. **E)** CI values representing synergy between FK866 and TX in metastatic PCa lines (mCRPC lines, mCSPC lines, and Clonally-derived mCRPC lines). Combination Index (CI) and Dose reduction (DRI) values were calculated according to Chou-Talalay’s method. CI values between 0.9-0.3 and 0.3-0.1 signify synergism and strong synergism, respectively, between the drugs treated in combination. **F)** Dose-response plots showing FK866 in combination with the androgen receptor inhibitor Enzalutamide combination in metastatic PCa cell lines.

Interestingly, we also demonstrated synergism between FK866 with the AR antagonist Enzalutamide as combination treatment in *in vitro* models of mCSPC. **Figure 3F** shows the cell survival curves representing the combination treatment with FK866 and the AR inhibitor Enzalutamide in mCSPC cell lines indicating that FK866 not only synergizes with Taxanes, the combination of FK866 and AR antagonists also showed significant synergy.

To further determine the synergy of these compounds, we performed the assessment of annexin V (a marker of apoptosis) and PI (a marker of necrosis) using flow cytometry in DU145 and DUTXR cells following exposure to FK866 as a single agent as well as in combination with DTX. Our results in **Figure 4A** showed significant increases in cells staining positive for annexin V and PI following FK866 and FK866+DTX treatment compared to control and DTX treatment, confirming significantly higher treatment-induced apoptosis post FK866-treatment. To confirm whether the loss of cell viability following FK866 treatment was indeed due to apoptosis, schedule-dependent effects of FK866 single-agent and combination treatments were determined by measuring the caspase-3/7 activity at estimated IC_50_ (**Figure 4B**). Our results showed that treatment with FK866 and DTX+FK866 induced apoptosis in every cell line compared to control (no drug treatment) cells. The relative increase (fold change) in Caspase 3/7 activity following were 3.09, 2.96, 2.36, and 2.66 for FK866 single-agent treatment and 3.96, 5.94, 6.23, and 4.66 for FK866+Taxane combination treatment in PC3, PC3M, DU145, and DUTXR, respectively.

**Figure 4.**
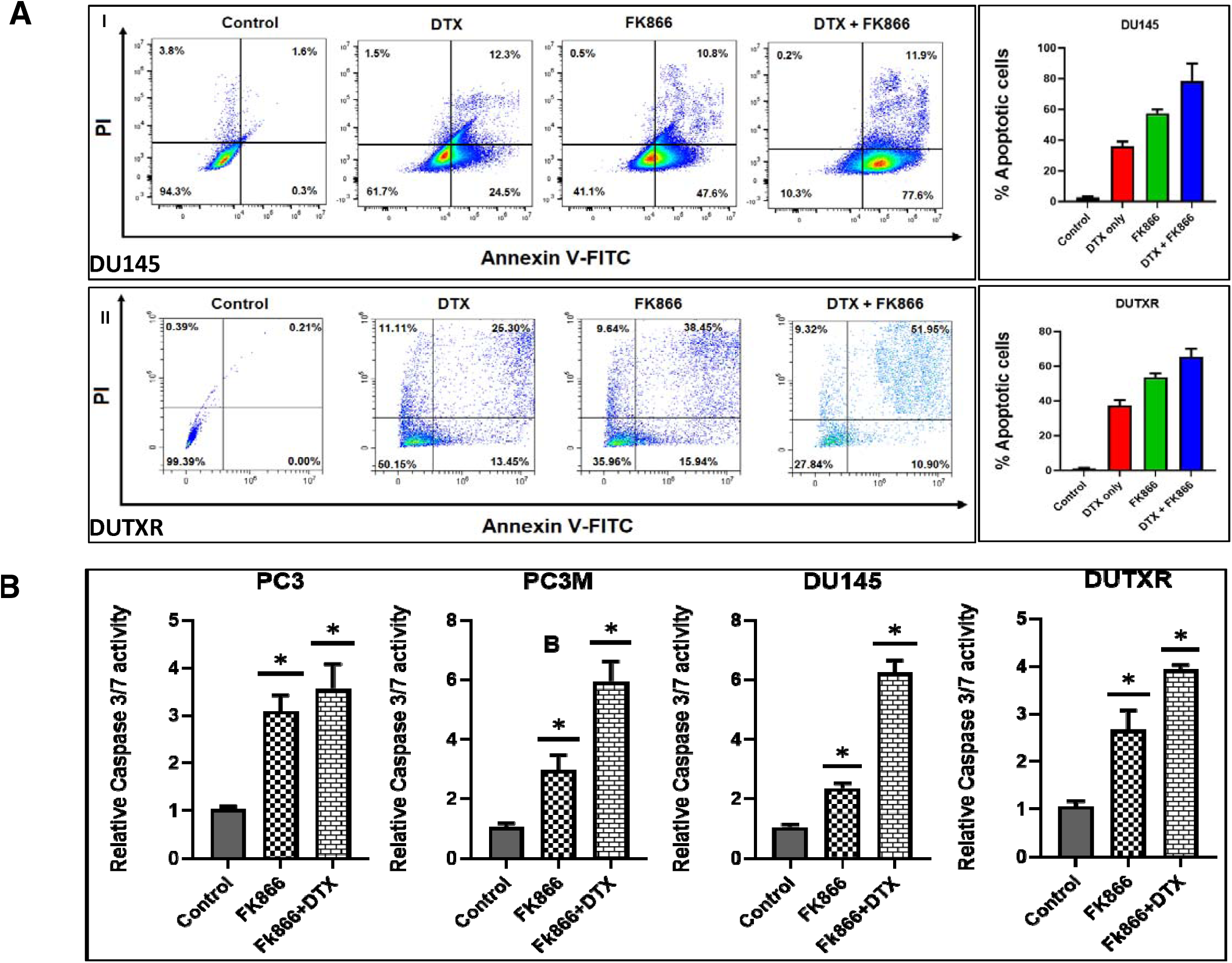
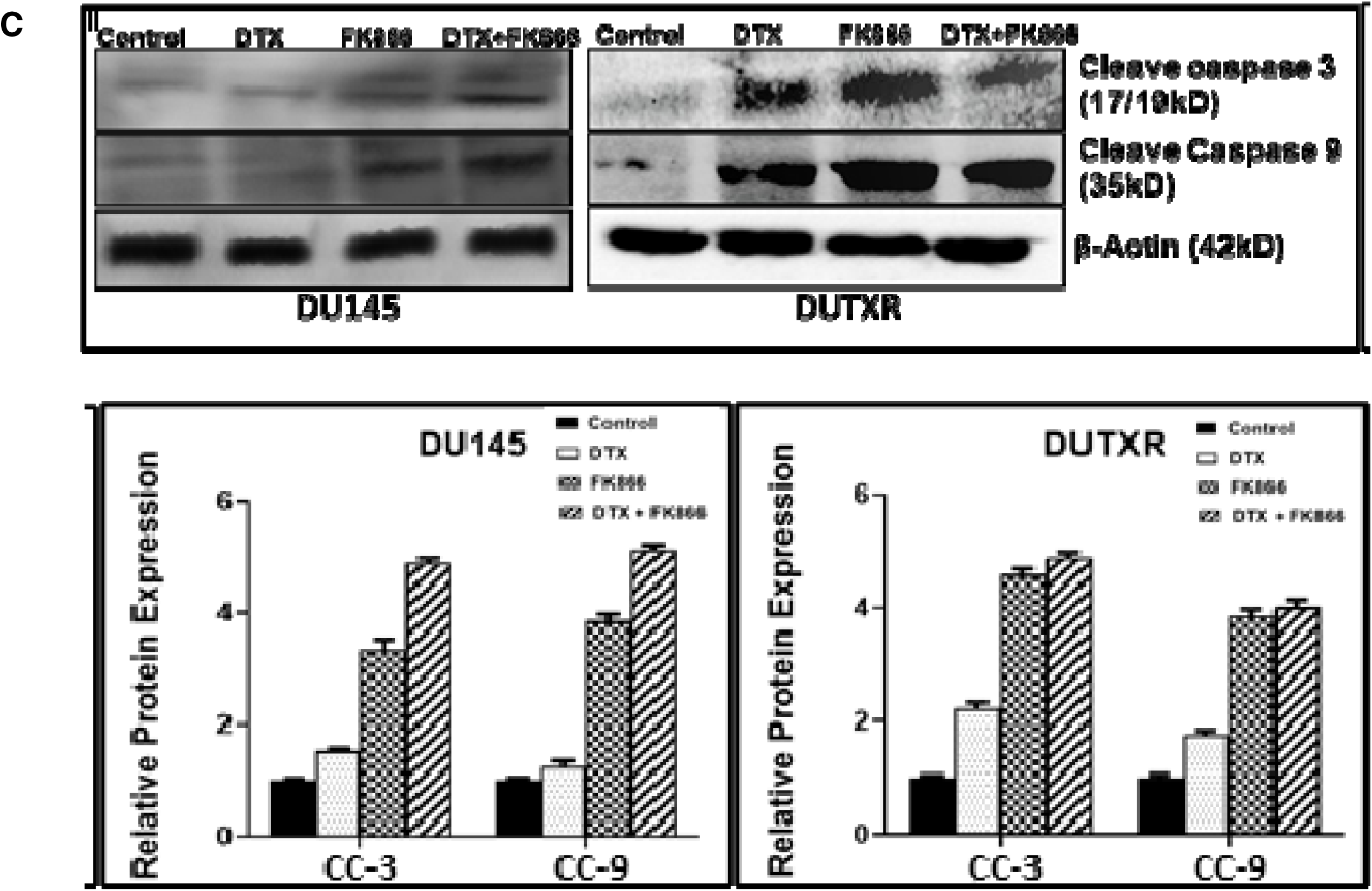
FK866 -induces apoptotic cell death in lethal Pca. **A)** Representative figures showing Annexin-V positive cells measured using flow cytometry following FK866 single-agent and DTX+FK866 combination treatment; Bar plot shows higher apoptosis in combination treatment compared to single-agent treatment (Significance P-value * = p ≤ 0.05). Apoptosis was confirmed using **B)** Caspase3/7 enzyme activity assay and **C)** Western blotting (expression of cleaved caspase 9 and 3 proteins. Representative plots showing DU145 and DUTXR lines are presented. Similar results were obtained for all metastatic PCa cell lines.

Pre- vs. Post FK866 single-agent/combination treatment immunoblotting results of the apoptotic markers cleaved caspase 9 and cleaved caspase 3 corroborated with the finding of our caspase 3/7 assay (**Figure 4C**). Interestingly, among all the treatment regimens, the highest level of apoptosis was observed for combination treatment with FK866+DTX.

Finally, we also showed that FK866 treatment reduced mCRPC cell density and changed nuclear morphology. Cells were exposed to FK866 treatment as a single agent as well as in combination with DTX, and cellular morphology was assessed using phase-contrast microscopy. In agreement with MTT assays, micrographs of PCa cells exposed to both FK866 and FK866+DTX dosing regimens showed decreases in cell density compared to control cells, as shown in **Figure 5A**. Further, assessment of nuclear morphology of attached cells using NucBlue staining, a reagent frequently used to distinguish condensed nuclei in apoptotic cells, suggested FK866-induced morphological changes like nuclear fragmentation and chromatin condensation, which are indicative of apoptosis (**Figure 5B**). In addition, our results showed even higher cell death and more nuclear damages in FK866+DTX combination treatment compared to FK866 single drug treatment.

**Figure 5.**
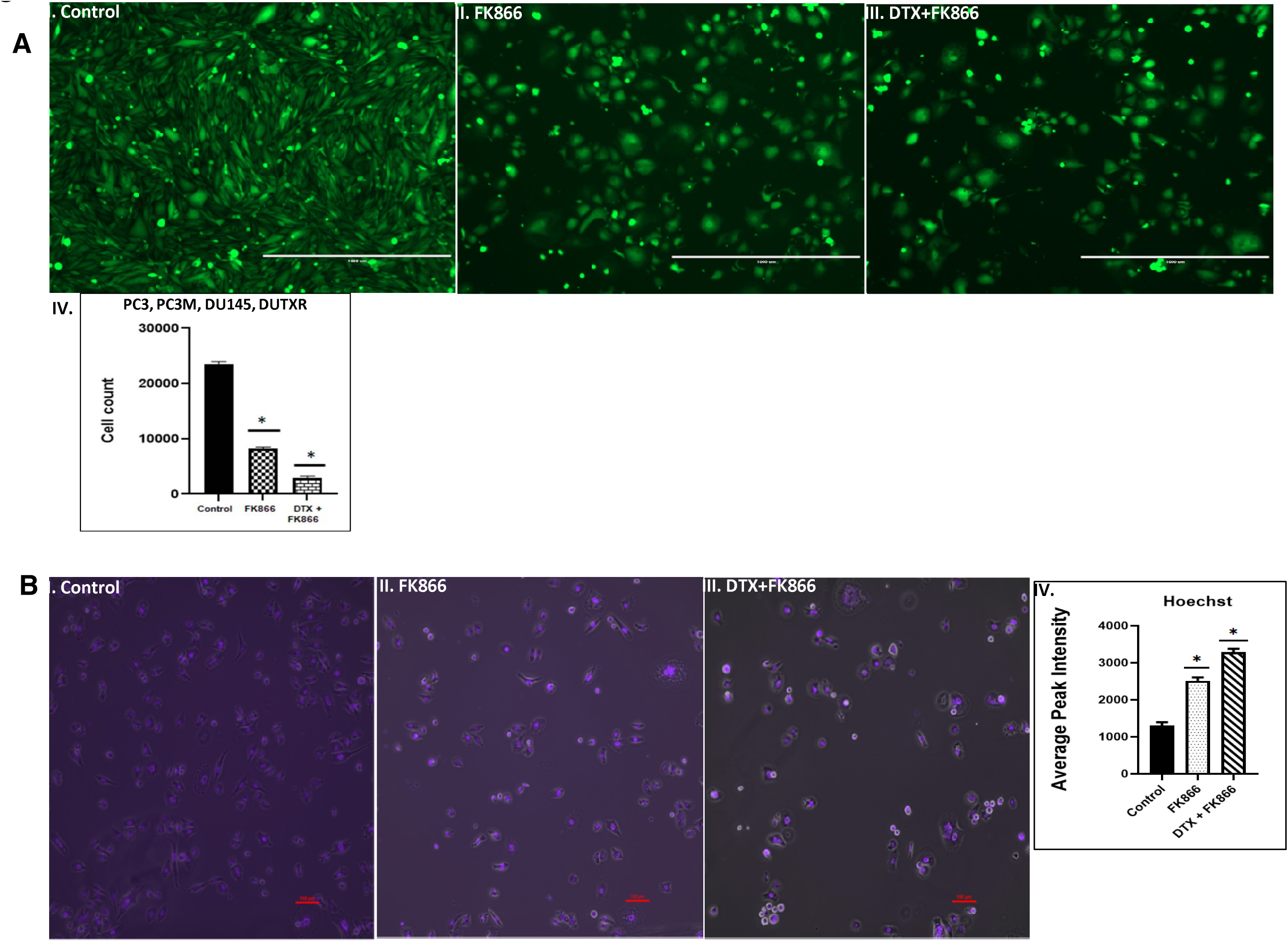
FK866 treatment reduced cell density and changed nuclear morphology. **A)** Assessment of cellular morphology: I-III. Representative figures showing the effect of the primary (DTX) and secondary (FK866) drugs on cell count and cell morphology of mCRPC cells. The images were captured on the PC3-Luc cells before (0h) and after (48h) FK866 treatment either as a single agent or in combination. Microscopy results show significantly higher cell death in combination treatment compared to single-drug treatment for all cell lines; IV. ImageJ data analysis showed a significant difference in cell density for both FK866 single-agent and DTX+FK866 combination treatments. Results show significantly higher cell death in combination treatment at combination dose compared to single-drug treatment for PC3cell lines (Significant value * = p ≤ 0.05). **B)** Assessment of nuclear morphology. I-III. Representative figures showing the FK866- based treatment (single-agent and combination with DTX) on cell nucleus morphology of mCRPC cells. NucBlue Live reagent is frequently used to distinguish condensed nuclei in apoptotic cells. The microscopy images were captured on the PC3-Luc cells before (0h) and after (48h) following treatment. Microscope images showing treatment effect on the cell lines PC3. Similar results were obtained for all mCRPC lines; IV. ImageJ data analysis showed a significant difference in cell density and nucleus damage for FK866 regimens (Significance P-value * = p ≤ 0.05).

### A microfluidic screen showed FK866 is potentially effective against EMT transdifferentiation and metastasis in treatment-refractory aggressive subclones

A Polydimethylsiloxane-PDMS-based μ-channel assay served as a physiologically relevant *in vitro* metastasis model for screening our top secDrugs. This allowed us to study the effect of our drug combination on tumor cell motility through μ-channels of dimensions that mimic the size of channel-like tracks encountered by migrating cells *in vivo*(Paul *et al*., 2017; Weigelin *et al*., 2012). Briefly, we fabricated a PDMS-based μ-channel assay using standard multilayer photolithography and replica molding as previously demonstrated (Mistriotis *et al*., 2019; Wisniewski *et al*., 2020; Wong *et al*., 2019). The device consisted of an array of parallel channels of variable width (3-50 μm) and with fixed length (200 μm) and height (10 μm). Perpendicular to the μ-channels were two larger 2D-like channels that served as cell seeding and chemoattractant inlet lines (**Figure S3**). Prior to cell seeding, the μ-fluidic devices were coated with 20 μg/mL rat tail collagen type I (Corning) for 1 hour at 37°C to facilitate cell adhesion. 1-1.5 x 10^5^ vehicle or FK866-treated mCRPC cells were introduced into the cell seeding inlet line via pressure-driven flow and allowed to adhere for 30 min at 37°C, 5% CO_2_. Next, the cell suspension was removed and substituted with a serum-free medium. Medium supplemented with 10% FBS was added into the chemoattractant inlet line to trigger cell entry into the channels. The devices were placed on an automated Nikon Ti2 Inverted Microscope equipped with a Tokai Stage-Top incubator unit, which maintains cells at 37°C and 5% CO_2_. Cell motility was recorded via time-lapse microscopy. Images were taken every 20 min for 10 hours with a 10x /0.45 NA Ph1 objective. To assess the migration efficiency of drug-treated mCRPC cells compared to control, we calculated the percentage of cell entry into the microfluidic channels defined as the total number of cells entering the channels divided by the total number of cells seeded within 50μm diameter from the μ-channel entrances. Because our prostate cancer cells did not frequently enter narrower microchannels (≤10 μm), we focused our analysis on wider channels (≥20 μm).

Our microfluidic-based cell migration assay revealed that our FK866 single-agent and FK866+DTX combination treatment reduced cell entry into 50 and 20 μm wide channels, suggesting that these interventions may potentially suppress prostate cancer cell invasion and possibly metastasis (**Figure 6A** and **Videos**).

**Figure 6.**
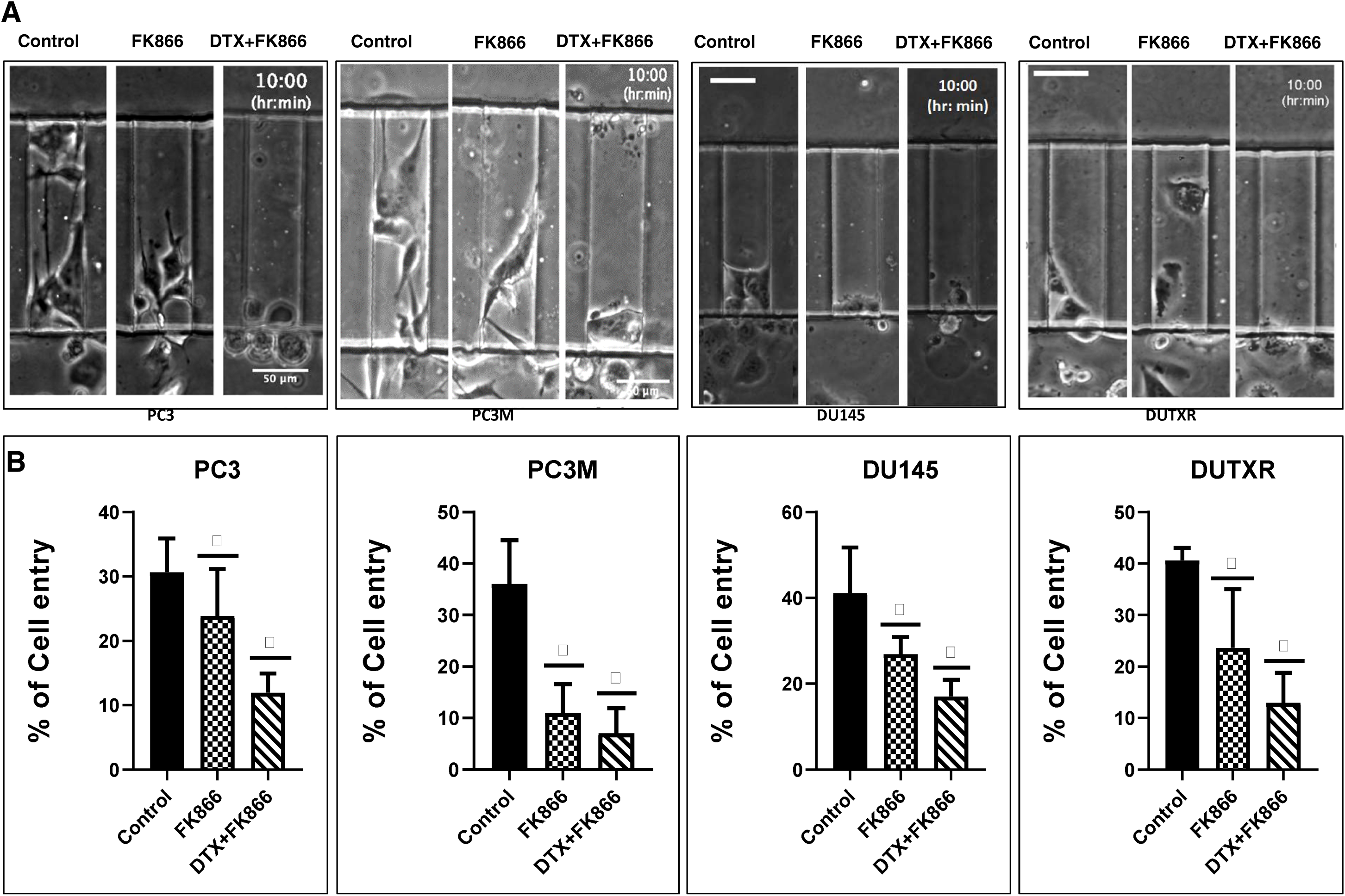
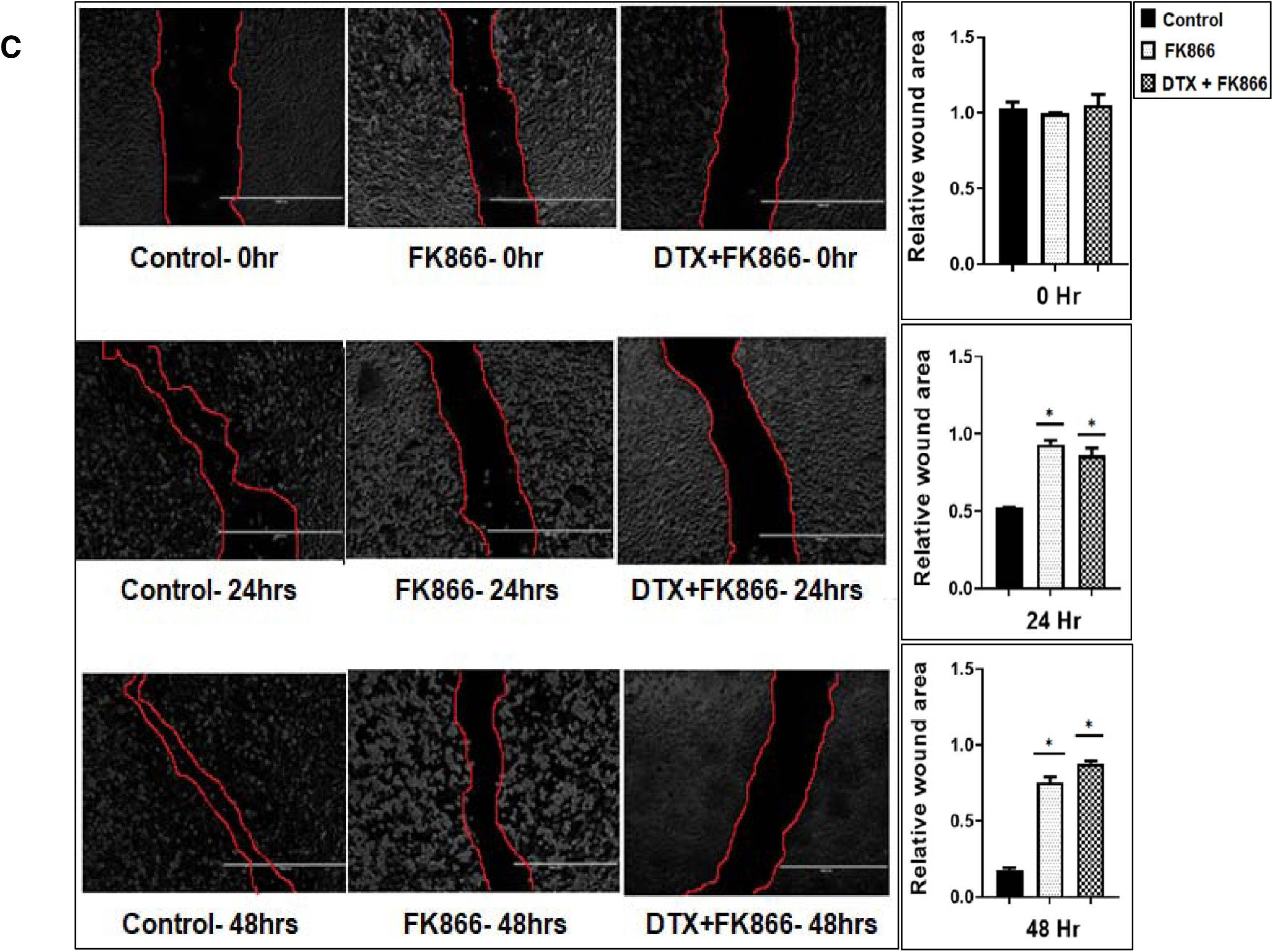
FK866 reduces cell migration and is potentially effective against metastasis and EMT transdifferentiation in lethal Pca. **A) Representative images of control, FK866 and DTX+FK866-treated prostate cancer cells migrating through 50 μm wide μ-channels** The fabrication of a μ-channel assay is described in the *Methods* section. Reduced cell entry into channels indicates a more generic defect in the migratory capacity of tumor cells. B) FK866 single-agent and combination therapy with DTX reduces the entry of lethal PCa (aggressive mCRPC and acquired taxane resistant mCRPC lines) into 50 and 20 μm wide μ-channels, indicating a potential role of FK866 in abrogating the metastatic potential. n≥3 experiments (Significance P-value * = p ≤ 0.05 relative to control). C) **Scratch assay:** Representative plots showing results of wound healing (Scratch) assay. Cell migration after 24 and 48h FK866 single-agent and FK866+DTX combination were assessed by measuring the scratch size. Images were captured before (0h) and after (24h, and 48h) drug treatments (Significance P-value * = p ≤ 0.05). Bar graphs showed a significant reduction in cell migration (wound healing) following FK866-based single- agent and combination treatments.

Further, FK866+DTX combination therapy showed a higher reduction in cell migration in AI-mCRPC/NEPC cells (PC-3, PC-3M, DU145) and acquired taxane resistant mCRPC (DUTXR) cells compared to single-agent FK866 treatment (**Figure 6B**).

To confirm FK866 treatment-induced reduction of mCRPC cell migration, we performed scratch assays by creating a “scratch” in cell monolayer followed by capturing the images at the beginning and regular intervals (0, 24, and 48hr for all treatments - single dose FK866 at the calculated IC_50_ and FK866+DTX combination at estimated IC_50_) during cell migration to compare the images and quantify the migration rate of the cells to close the scratch. Our results showed that cell migration was higher in control cells compared to FK866 and FK866+DTX post-treated cells (**Figure 6C**). Further, combination treatment (FK866+DTX) had a higher effect in reducing cell migration in PCa cell lines compared to treatment with FK866 alone (p<0.05).

### FK866 showed selective on-target inhibition of NAMPT activity and a distinct impact on gene expression signature

The effect of FK866 on its intended target, NAMPT, was assessed using NAD/NADH-Glo activity assay that measures the ratio between total oxidized and reduced nicotinamide adenine dinucleotides (NAD^+^ and NADH, respectively). We found that FK866 selectively inhibited the total cellular NAD+/NADH ratio in all PCa cell lines in a dose-dependent manner. **Figure 7A** demonstrates a significant (p<0.05) decrease of NAD+/NADH ratio following 24h FK866 IC_50_ and IC_50_/2-treatment compared to the control cells. (Range: 2.70 to 12.35 for IC_50_, and 2.15 to 10.01 for IC_50_/2).

**Figure 7.**
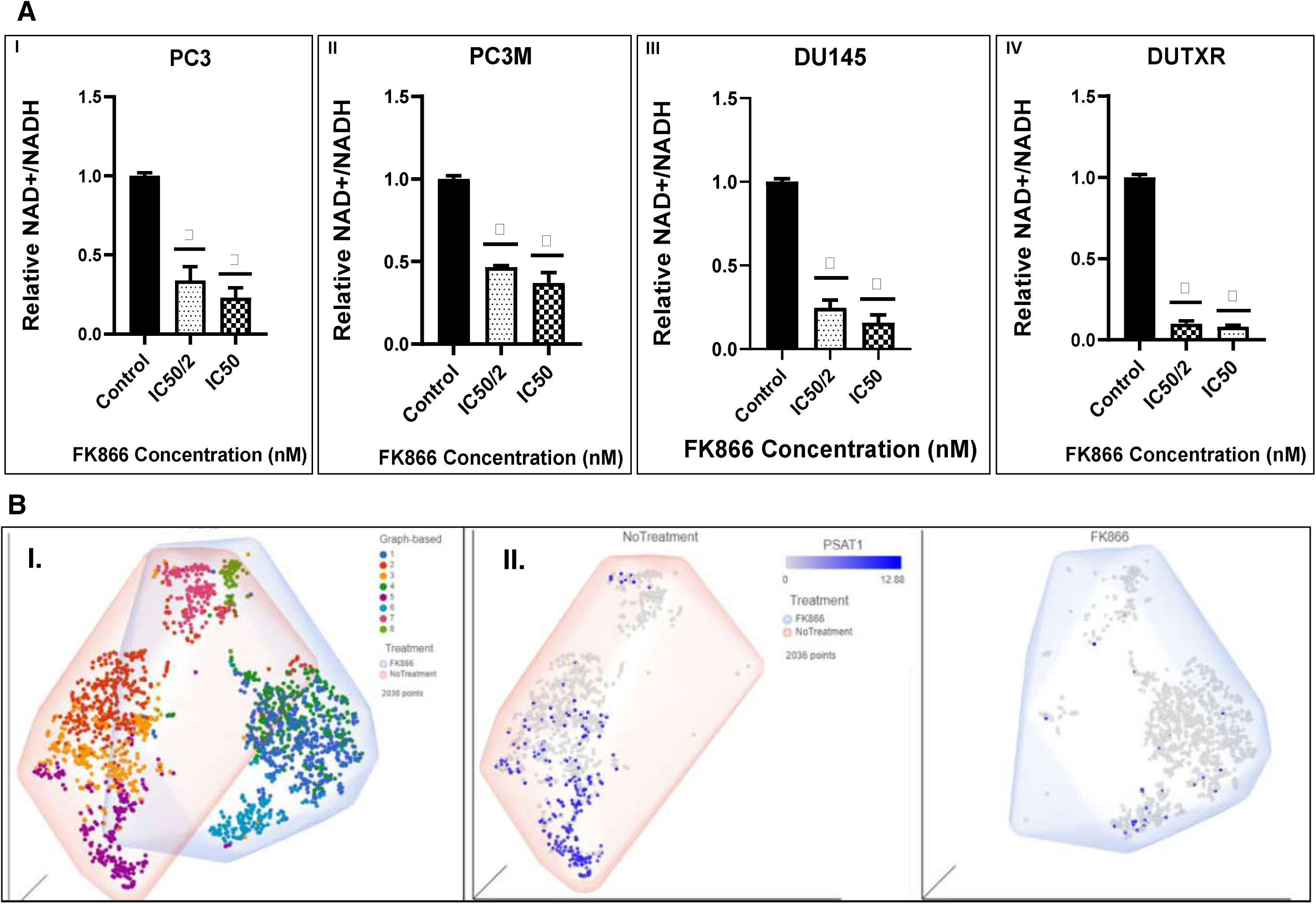
FK866 showed selective on-target inhibition of NAMPT activity. **A) FK866 post-treatment NADH activity:** Effect of FK866 on total cellular NAD+/NADH ratio in PCa cell lines was detected by NADH activity assay in AR^lo^ mCRPC (PC3, PC3M, DU145) and in Acquired taxane resistant mCRPC (DUTXR) cell lines treated with FK866 doses equivalent to IC50 and IC50/2 of each cell lines. A significant decrease in the NAD+/NADH ratio was observed compared to control cells after 24h of treatment. Results are presented as mean ± SD; n = 3 for each experiment (*P < 0.05, compared to untreated controls). **B) Pre- vs. Post-treatment scRNAseq data analysis:** Effect of NAMPT inhibitor (FK866) on the sub-clonal population of human prostate cells in vitro following single-cell transcriptomics analysis. **I.** Pre- vs. Post- treatment t-SNE clusters, showing **II.** Erosion of the PSAT1^high^ cluster (Cluster 5) in the FK866-treated scRNAseq data.

The on-target effect of FK866 was confirmed at the single-cell level using post FK866-treatment scRNAseq datasets compared to baseline/untreated PCa (**Figure 7B**). Our single-cell transcriptomic analysis showed that FK866 treatment resulted in the loss of the tSNE cluster 5 (the PSAT1^high^ cluster) following FK866 treatment represented by high expression of PSAT1 gene, indicating FK866-induced down-regulation of PSAT1 gene, which is a downstream protein in NAMPT-mediated NAD salvage biosynthesis pathway.

Next, we performed global whole-transcriptome profiling by bulk tumor RNAseq to compare changes in gene expression induced by FK866 in AVPC cell lines. GEP data were normalized to baseline (no-treatment). Volcano plots in **Figure S4** show differentially expressed genes (DEGs) following DTX, FK866 single-agent or combination treatment in human mCRPC cell lines heatmaps were generated following differential gene expression analysis (**Figure 8A**). A total of 247 genes were uniquely differentially expressed above the significance threshold (p<0.05) at 48hr post-FK866 single-agent treatment, while 85 and 289 genes were differentially expressed following DTX single-agent and FK866+DTX combination treatments, respectively (**Figure 8B**).

**Figure 8.**
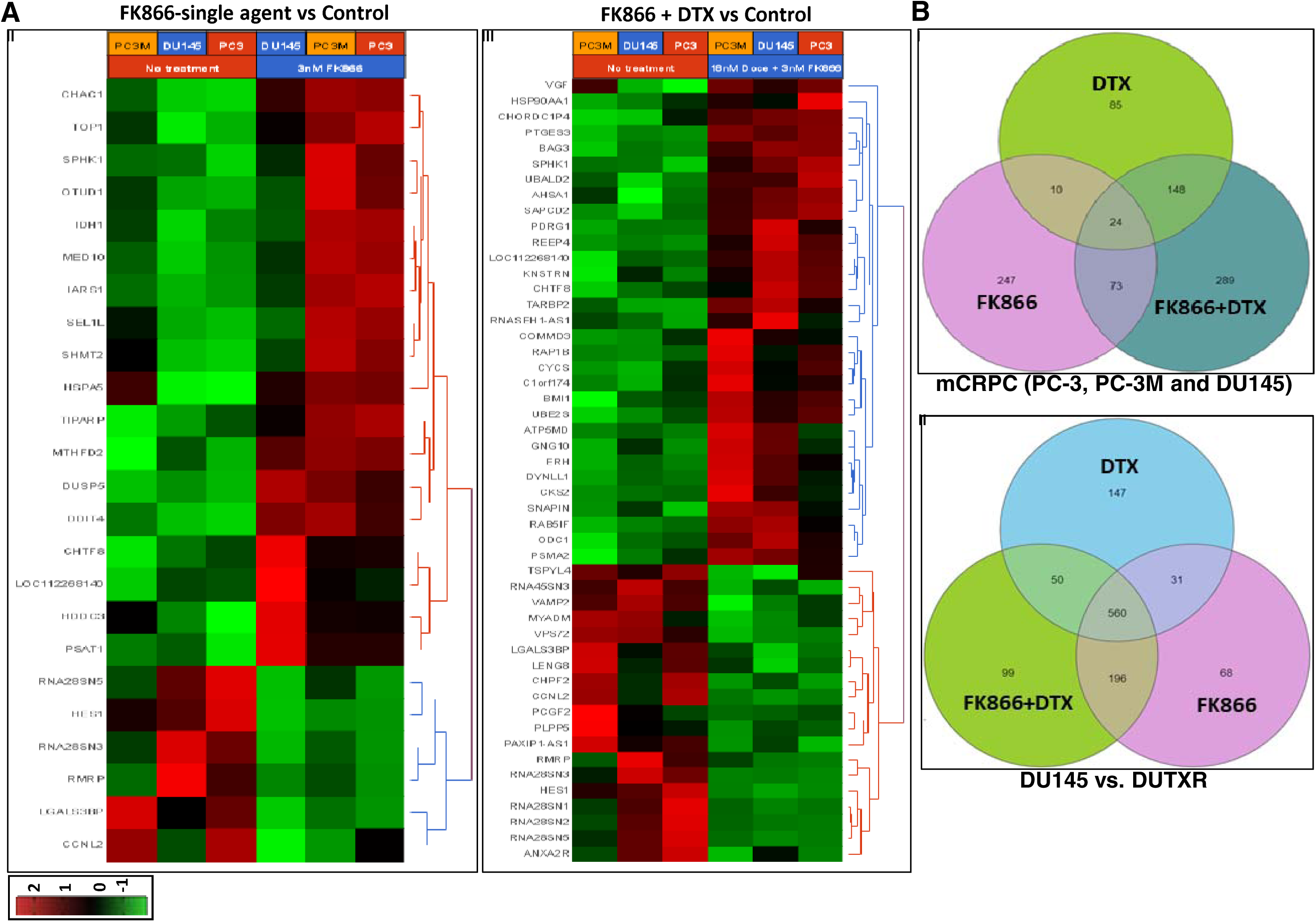
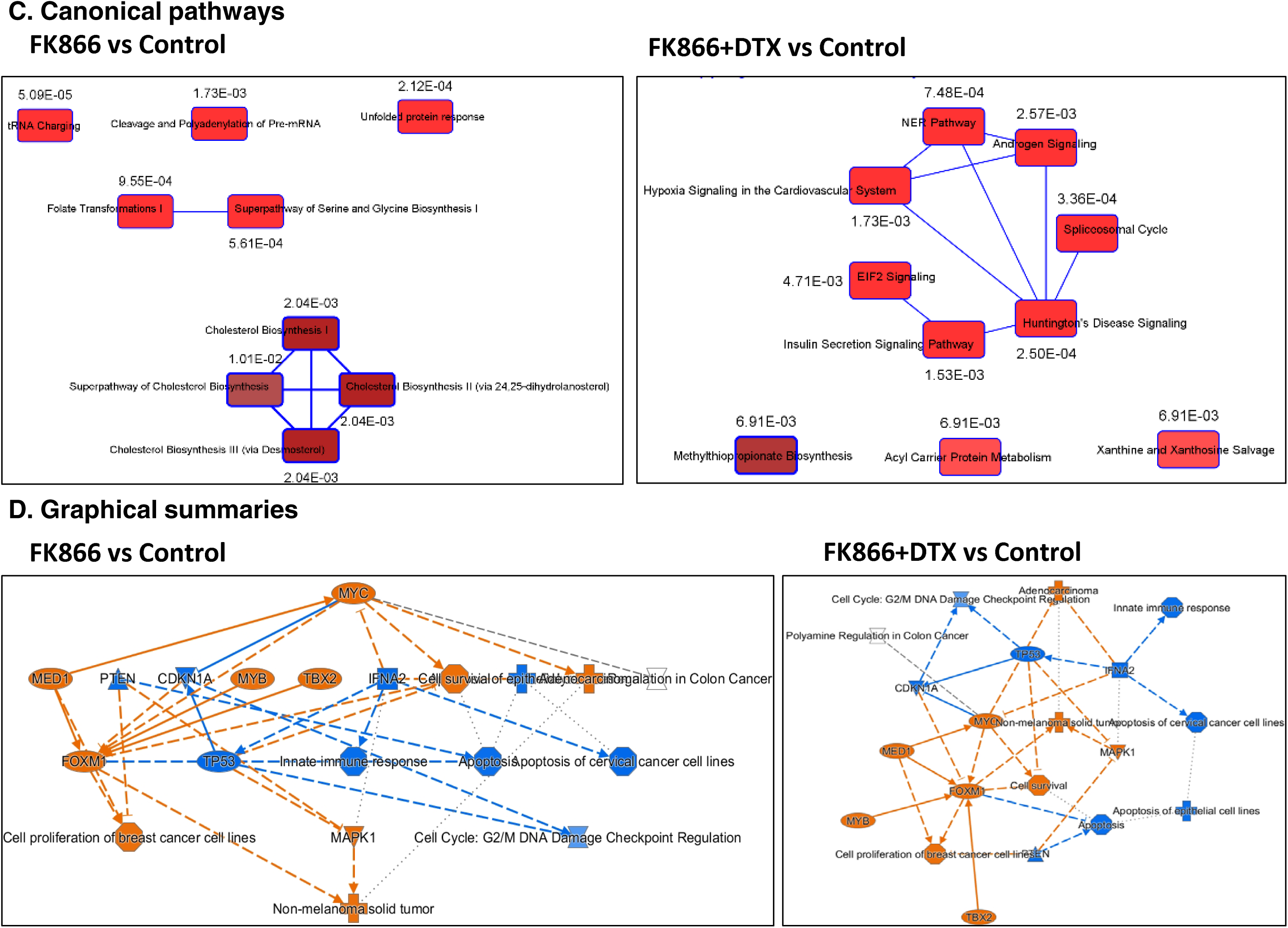
Differential gene expression profiling analysis results. **A)** Heatmaps representing top differentially expressed genes (DEGs) following DTX, FK866 single-agent, or DTX+FK866 combination treatments in human mCRPC cell lines (n = 3), 24h following drug exposure. I) FK866 single-agent treatment. II) DTX+FK866 combination treatment. Log_2_ ratios are depicted in a color scale where red represents upregulation and green represents downregulation. Columns represent cell lines, and rows represent genes. Prior to hierarchical clustering, gene expression values were filtered (samples with max TPM < 1 were removed) and z score normalized. **B)** Venn diagrams representing unique and common DEGs (p<0.05) between DTX, FK866, and DTX+FK866 treatments among: I. All mCRPC lines; II. DU145 vs. DUTXR lines in all cell lines. **Ingenuity pathway analysis:** IPA analysis was performed based on the top DEGs following FK866 single- agent of FK866+DTX combination treatments. **C)** Top canonical pathways. **D)** Graphical Summaries.

**Table S2** lists the top differentially expressed genes (DEGs). The top 10 downregulated DEGs between pre- (untreated; 0 h) vs. 48hr post-treatment, irrespective of treatment type (FK866, FK866+DTX, or DTX), were C1S, IFITM3, FAM229A, LHFPL5, DUS4L-BCAP29, ZACN, LGALS3BP, CXCL2, HES1, and FAS. On the other hand, the top upregulated DEGs were ELMOD1, SESN3, CYP1A1, DDIT4, SLC7A11, FTCDNL1, LAMP3, BHLHA15, PAPPA2, and EML5.

Among these, HES1 was downregulated in all treatment groups, with the highest level of downregulation (∼ 6- fold) following combination treatment (FK866+DTX). FAM229A, LGALS3BP, LY6G5B, MAT2A, LTB4R, NSUN5P2 were downregulated in both FK866 single-agent and combination (FK866+DTX) groups, while PITPNC1 was upregulated in both the groups, albeit at different levels.

IPA analysis performed based on the top DEGs associated with FK866 single-agent treatment revealed Inhibition of Matrix Metalloproteases (p=5.866e-03), oxidative stress, and cell cycle among the top dysregulated pathways (**Figure 8C**). Further, ATF4 (p=3.97E-10) was predicted as the top upstream regulator. Interestingly, G2/M DNA Damage Checkpoint Regulation (p= 8.62E-03) and Kinetochore Metaphase Signaling Pathway (p= 5.37E-03) were among the top canonical pathways inferred from the following genes differentially regulated between FK866+DTX vs. baseline: MED1, FOXM1, MAPK1, MYB, TBX2, CDKN1A, PTEN, TP53, TFNA2. Further, IPA predicted IFNA2 as the top upstream regulator based on the expression of genes (**Figure S5A**). Among microRNAs, IPA predicted mir-1-3p as the top upstream regulator based on the expression of target genes and IFNA2 (**Figure S5B)**.

The graphical summaries in **Figure 8D** show FK866-based treatment regimens induce upregulation of apoptosis in tumor cell lines, as well as the genes like STAT1, IRF1, DDX58, TNF, IFNG, and downregulation of tumorigenesis in tissues, polyamine regulation, and the genes FOXM1, MAPK1, AREG, etc. A detailed heatmap comparing differentially regulated pathways between FK866 single-agent and combination treatments are provided in **Figure S6,** included differential regulation of the following pathways between the treatment regimes Putrescine Biosynthesis, Prostanoid Biosynthesis, Eicosanoid Signaling, Cell cycle-G2/M damage, Kinetochore Metaphase signaling, Polyamine regulation, Phosphatidylethanolamine Complement system, P53 signaling, tRNA charging, Matrix metalloprotein, and Ferroptosis Signaling Pathway.

Since cell cycle and checkpoint regulation were significant among the IPA-predicted signaling pathways, we performed *in vitro* assessment of cell population in each cell cycle checkpoint (G2/M) following FK866 single-agent and DTX combination treatment using Flow cytometry. This effect of FK866 either alone or in combination on the cell cycle distribution was assessed by quantifying DNA content. We observed that treatment of PCa cell lines with FK866 resulted in G2/M checkpoint arrest. Furthermore, a higher number of cells were arrested at G2/M following combination treatments compared to FK866 or DTX alone (**Figure 9A**). Treatment of cells with FK866+DTX increased the highest percentage of cells in the M phase (p<0.05) with a concomitant decrease in S and G0/G1 populations.

**Figure 9.**
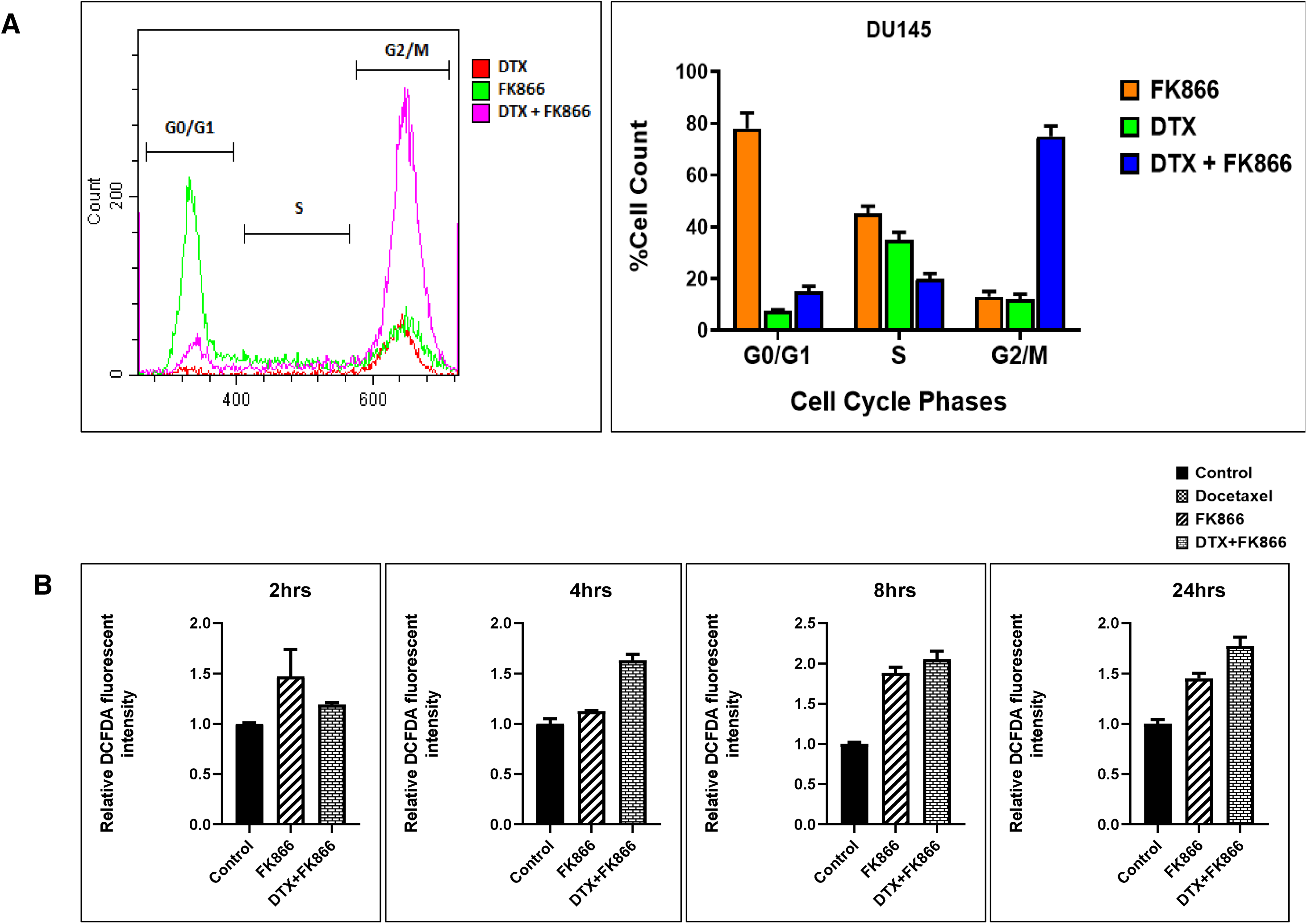
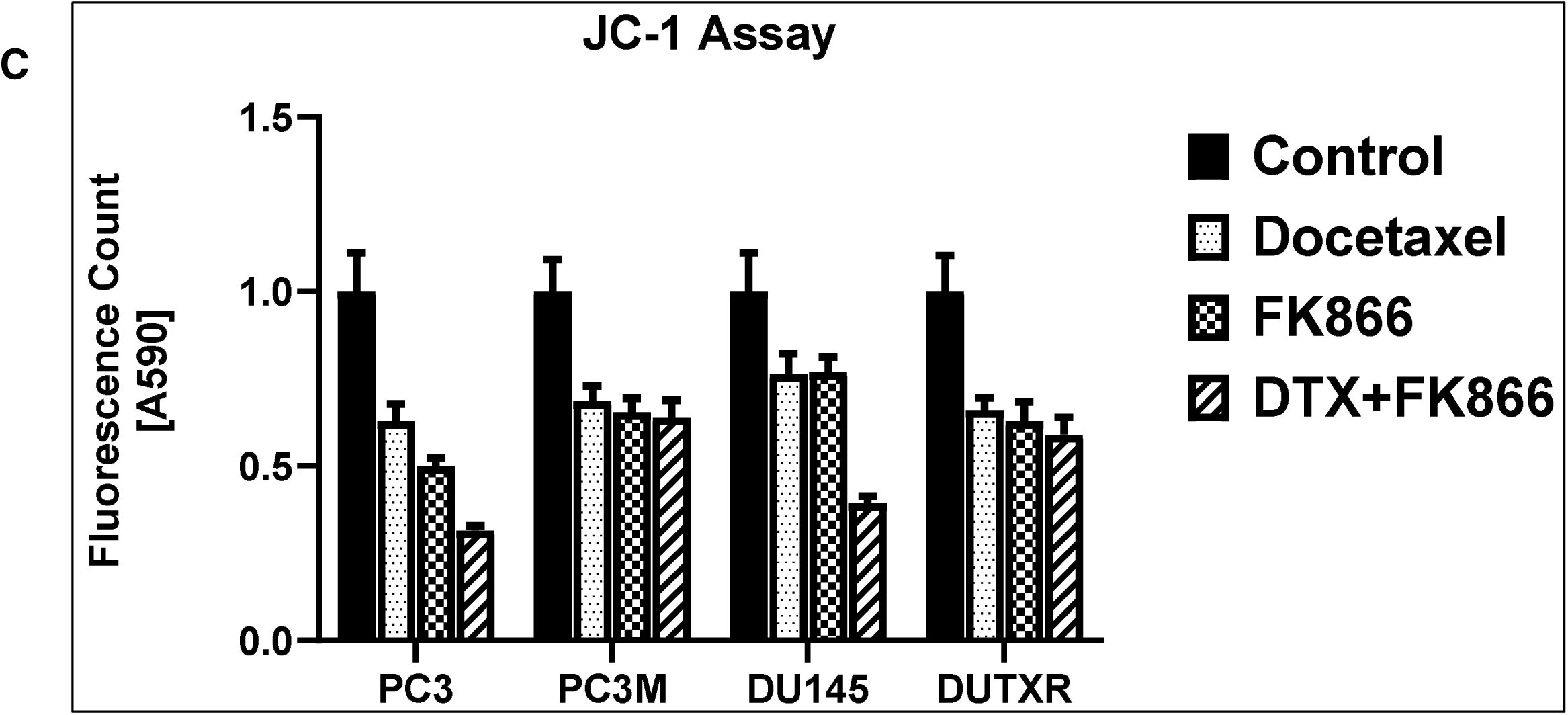
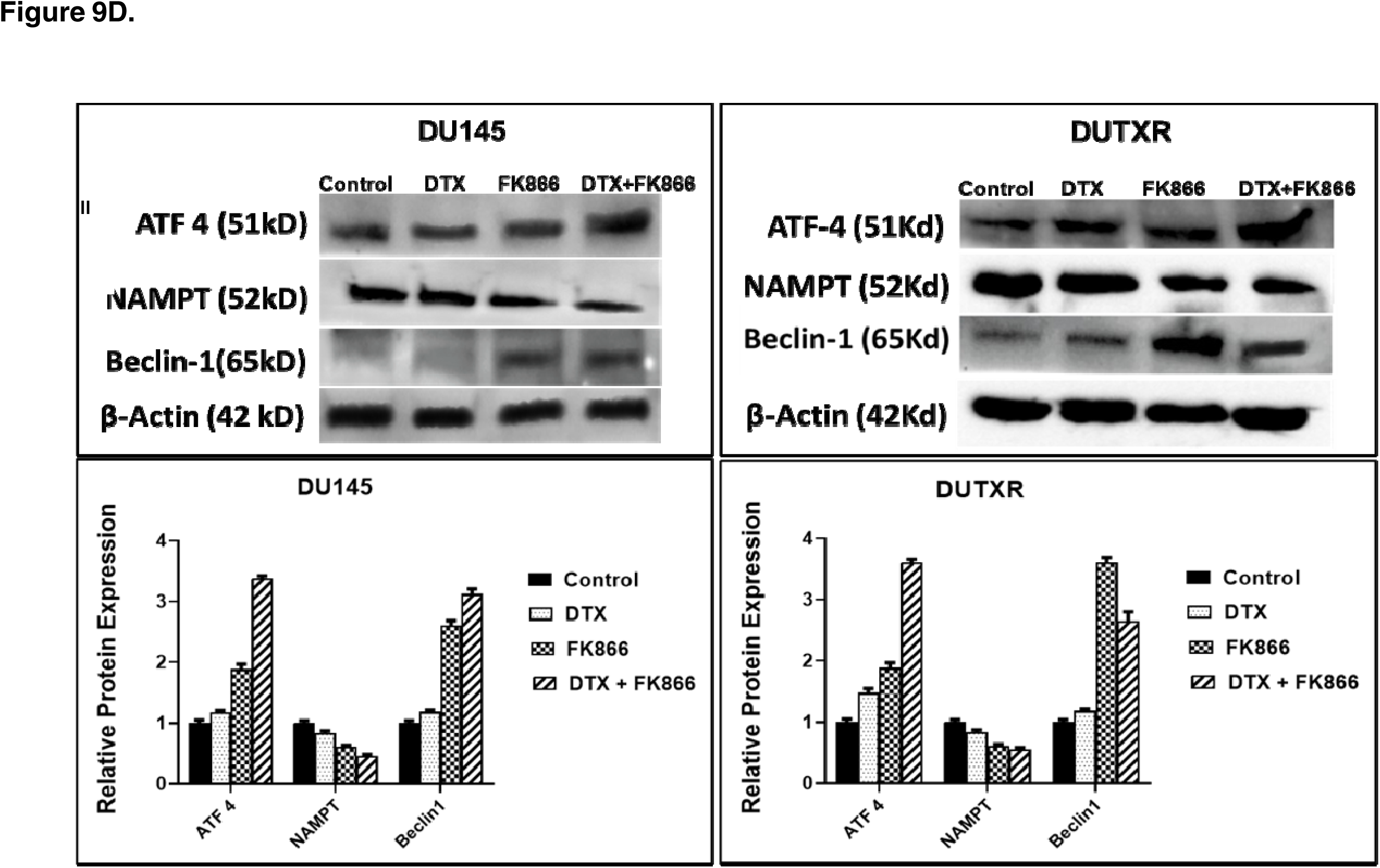
Functional validation of top key pathways. **A) Effect of FK866 treatments on Cell cycle:** Representative figure showing results on the DU145 cell line. Cells were exposed to DTX, FK866, DTX+FK866 treatment for 24h, stained with propidium iodide (PI), and cell cycle phases were assessed by flow cytometry. Cell cycle stages were analyzed by CytExpert (Beckman Coulter). FK866 was shown to arrest cell cycle at the G2/M checkpoint. Cell Cycle in treatment groups was normalized to the corresponding control. (Significance P-value * = p ≤ 0.05). Data are presented as the mean ± SEM of three separate experiments (n = 5/study). **B) FK866 post-treatment ROS generation measured by DCFDA assay:** ROS generation was measured after 2, 4, 8, and 24h DTX, FK866 single-agent, and in DTX+FK866 combination treatments. **C) Mitochondrial Membrane Potential:** FK866 treatment induced mitochondrial dysfunction was measured by JC-1 Assay. **D) Western blotting:** ER stress and Autophagy markers Beclin-1 and ATF-4 protein level was measured after DTX, FK866, and DTX+FK866 treatment. Post-treatment FK866 exhibited upregulation of Beclin-1 and ATF-4, as well as NAMPT compared to control (no drug treatment). **B.** Densitometry analysis confirmed top FK866 treatment-related genes/ pathways.

Among the IPA-predicted pathways following FK866 treatment were oxidative stress and mitochondrial dysfunction. Intracellular ROS levels are the indicator of oxidative stress. Therefore, to validate the mechanism of action of the drug combination, we quantified the intracellular ROS level in the pre-and post-treatment condition of FK866 as a single-agent vs. combination with DTX using the fluorogenic probe 2,7-dichlorofluorescein diacetate (DCFDA), a cell-permeable non-fluorescent probe that shows fluorescence when it is oxidized. Cellular superoxide anions were measured by using the fluorescent dye DHE. **Figure 9B** depicts significant ROS generation following 2, 4, 8, and 24hr DTX or FK866 single agent and DTX+FK866 combination treatments. Further, combination treatment exhibited higher ROS generation than single-agent treatment and control in Acquired taxane resistant AI-mCRPC (DUTXR) cell lines (p ≤ 0.05).

Mitochondrial membrane potential and FK866 treatment-related mitochondrial dysfunction were accessed using JC-1 (Sigma). JC-1 is a cationic carbocyanine dye that accumulates in mitochondria. FK866 treatment enhanced mitochondrial dysfunction in AR^lo^ mCRPC/NEPC (PC-3, PC-3M, DU145) and acquired taxane resistant AI-mCRPC (DUTXR) cell lines (**Figure 9C**).

To confirm the results of our differential gene expression analysis, IPA pathway analysis, and *in silico* patient data validation analyses, we selected ATF4, Beclin-1, and NAMPT for immunoblotting in PCa cell lines. **Figure 9D** shows immunoblotting results confirming upregulation of ATF4 and beclin-1 and downregulation of the FK866 target gene NAMPT following FK866-based treatment.

### Validation of FK866 treatment-induced gene signatures using Patient datasets

RNAseq data on PCa patients were obtained from the Gene expression omnibus database (GSE54460)(Long *et al*., 2014). The dataset includes 100 PCa patients (49 with BCR, 51 with no BCR) from the Atlanta VA Medical Center, Moffitt Cancer Center, and Sunnybrook Health Science Center. First, we performed differential gene expression analysis between patients with or without biochemical recurrence (BCR). **Figure 10A** shows the top pathways that were significantly different between BCR vs. no-BCR based on DEGs with p<0.05. Next, as a reverse-matching approach, we compared the list of shared dysregulated (down or upregulated) genes with our list of top FK866-treatment-induced DEGs. **Table S3** lists the genes that were up-or down-regulated in PCa patients with BCR AND had significant fold changes in the opposite direction following FK866 treatment in our model systems, indicating that FK866 might be capable of reversing the input signature in the patient cohort. The top genes that were significantly upregulated in patients and showed significant downregulation following FK866 treatment in PCa cell lines were LTB4R, IFITM3, and TMEM120B. Finally, **Figure 10B** shows that several FK866 treatment-induced pathways were significantly downregulated in PCa patients with BCR.

**Figure 10.**
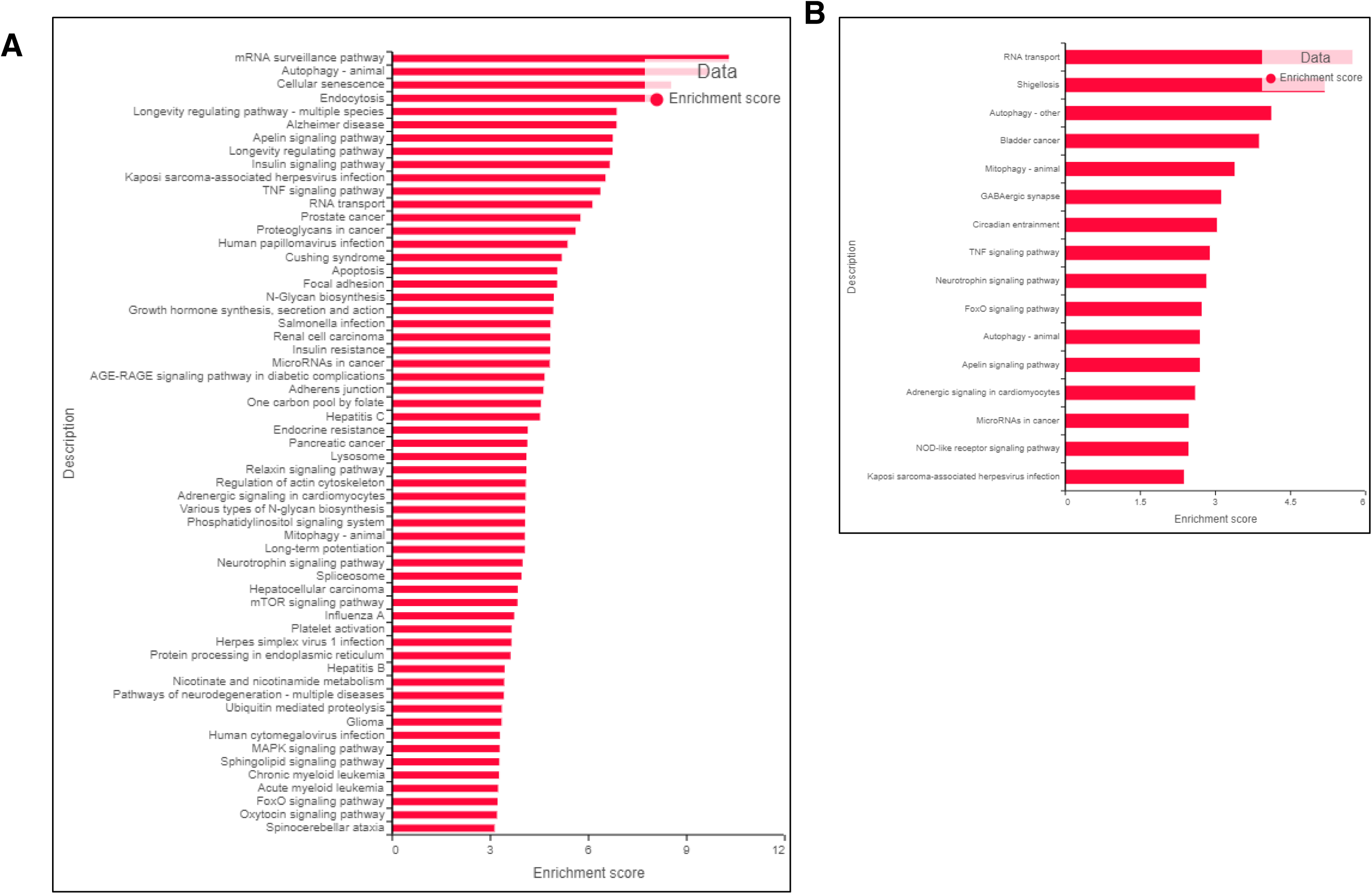
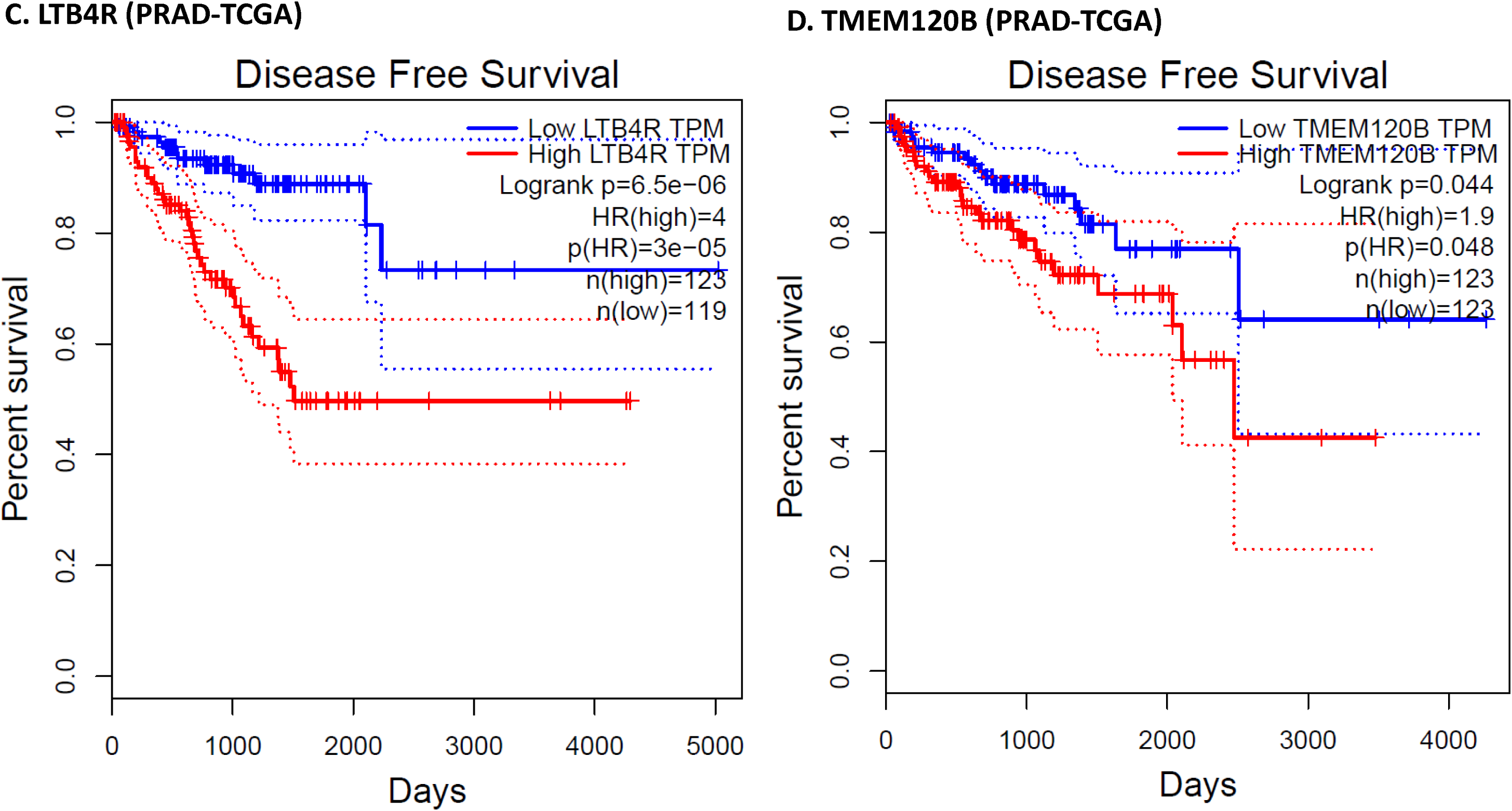
Validation of FK866 treatment-related gene signatures using patient cohort datasets. Reverse-matching using patient cohort datasets show FK866 treatment has the potential to reverse PCa lethality. Pathway analysis was performed based on the top DEGs in **A)** PCa patient cohort; **B)** Top FK866 treatment induced upregulated pathways that were significantly downregulated in PCa patients with BCR. Validation using TCGA’s prostate adenocarcinoma (PRAD) GEP dataset: *In silico* analysis of the top FK866 treatment-induced downregulated pathways (**Table S3**) that were significantly downregulated in PCa patients with BCR using TCGA’s PCa dataset: Kaplan-Meier Curves showed that the genes **C)** LTB4R, and **D)** TMEM120B were significantly associated with disease-free survival.

Interestingly, additional *in silico* analysis using TCGA’s prostate adenocarcinoma (PRAD) GEP dataset showed that the genes LTB4R and TMEM120B were also significantly associated with disease-free survival (Kaplan-Meier curves are presented in **Figure 10C-D**), with Hazards Ratios 4 (p=0.00003) and 1.9 (p=0.048), respectively. Thus, this biomarker-based method served as a novel tool to screen drugs against aggressive PCa.

## 3. DISCUSSION

Drug development for aggressive and/or lethal treatment-resistant PCa poses a significant challenge with very few therapeutic successes(Conteduca *et al*., 2019; Scher *et al*., 2015). In this study, we introduced a pipeline that integrated a pharmacogenomics data-driven approach with scRNAseq-based rapid drug screening method and identified FK866 as a proof-of-concept secondary drug (‘secDrugs’) against lethal PCa including aggressive, acquired taxane resistant and stem-like cell types representing NEPC and stem-like (EMT) phenotypes. Notably, we used scRNAseq as an innovative approach to demonstrate that a subset of AR^low^ PCa cells in metastatic prostate cancer, including castration-sensitive and castration-resistant tumors, harbored signatures of Epithelial-mesenchymal transition (EMT) and cancer ‘stemness’ which we also showed as targets of FK866.

Earlier studies have reported that FK866 is a NAMPT inhibitor and controls cancer progression by down- regulation of TNFα, IL-6 expressions, CXCR4(Galli *et al*., 2020). NAMPT is the rate-limiting enzyme in the NAD+ salvage pathway(Audrito, 2020). Tumor cells have increased requirements for NAD+. Therefore, NAMPT inhibitors play a key role in energy metabolism in cancer. Furthermore, recent insights have revealed several additional targetable cellular processes that are impacted by inhibition of NAMPT, such as - sirtuin function (tumor cell proliferation and progression), DNA repair machinery, redox homeostasis (ROS), cellular stemness, and immune processes(Audrito, 2020; Galli *et al*., 2020; Heske, 2019; Tan *et al*., 2013; B. Wang *et al*., 2011). Concurrently, high NAMPT expression was also associated with the presence of a higher proportion of cancer-initiating/stem cells. The loss of cancer stem cells by inhibition of NAMPT was the result of an excess of autophagy, which disrupted the maintenance of cancer cell stemness(Nacarelli *et al*., 2020; P. Yang *et al*., 2015).

Since FK866 is an inhibitor of NAD biosynthesis, we confirmed that FK866 selectively inhibited NAD+/NADH ratio in PCa cell lines in a dose-dependent manner. Further, we combined a novel micro-fluidic-based cell migration assay, genome-wide bulk inter-tumor (RNAseq), and single-cell transcriptomics (scRNAseq) analysis to elucidate in detail the treatment-induced genes and molecular pathways/networks underlying FK866 mechanism of action and its potential impact on tumor metastasis, migration, invasion, intracellular ROS activity, autophagy and most importantly, ‘cancer stemness,’ in AVPC.

Among the top downregulated genes were the potential proto-oncogene IFITM3 (<-7-fold), C1S (<-8-fold), FAS (<-3.6-fold), as well as LGALS3BP, LTB4R, MAT2A, and CXCL2.

Interferon-inducible Transmembrane Protein 3 (IFITM3) has an oncogenic role and promotes cell proliferation, cell migration through EMT, invasion stemness, bone metastasis, and PCa progression through activation of a novel TGF-β-Smads-MAPK pathway(X. Liu *et al*., 2019; Rajapaksa *et al*., 2020). Therefore, IFITM3 expression is correlated with poor prognosis of PCa, as well as tumorigenesis, progression, differentiation, and tumor relapse(M. Yang *et al*., 2013; Zhao *et al*., 2013). Concurrently, we showed that baseline IFITM3 gene expression was upregulated in PCa patients with biochemical recurrence compared to patients without any BCR event. Furthermore, IFITM3 expression was also found significantly higher in TCGA-PRAD patients (>1000 folds; p= 0.00992) with African ancestry compared to Caucasians. Since African American (AA) men are disproportionally affected, more likely to develop PCa, less likely to respond to conventional therapies, and twice as likely to die from PCa compared to other ethnicities, this indicates a possible role of high IFITM3 expression in these inter-ethnic differences(Cronin *et al*., 2018; Howlader *et al*., 2015). In our study, we observed ∼7-fold downregulation of IFITM3 gene expression following FK866 treatment.

High expression LTB4R has been reported in various cancers, including PCa(Larré *et al*., 2008). Interestingly, we also observed that higher expression of LTB4R is associated with poorer disease-free survival (DFS) in the TCGA-PRAD PCa patient cohort. Our observed downregulation of LT4BR following FK866-based treatment regimens indicates its potential efficacy in improving patient survival. Thus, through reverse-matching, we showed that FK866 has the potential to reverse the effects of genes/pathways that were significantly dysregulated in PCa patients with a lethal/aggressive form of the disease.

Gene expression level of the complement protein C1S is significantly elevated in cancers and promoted cancer cell proliferation along with poor prognosis(Magrini and Garlanda, 2021). FAS protein overexpression indicates poor biochemical recurrence (BCR)-free survival in PCa(Cao *et al*., 2020; Gutierrez *et al*., 2005). Additionally, the androgen receptor directly binds to the Fas/FasL domain and promotes the androgen-independent growth of PCa(Gao *et al*., 2005). LGALS3BP has earlier been shown to be upregulated in human colorectal and prostate cancer that may influence oncogenesis and promote cancer growth and angiogenesis through the PI3K/AKT/VEGFA pathway(Song *et al*., 2021). MAT2A plays a crucial role in various cancer progression, including PCa(Maldonado *et al*., 2018; Munkley *et al*., 2018). Inhibition of MAT2A gene expression repressed the growth of human PCa(C. Ma *et al*., 2008).

Further, our study also showed that FK866 downregulates CXCL2 expression (Fold change <-3.5). The CXCL8 axis is beneficial for the regulation of cancer cell progression(Toraih *et al*., 2016). We also report that FK866 downregulated FOS, ATP1B3, and GOLGA8B, which have known benefits in PCa treatment. C-Fos is upregulated in advance PCa and correlated with Erk MAPK pathway activation with disease recurrence(Ouyang *et al*., 2008; Riedel *et al*., 2021; Shankar *et al*., 2016). ATP1B3 expression was increased in various cancers, including PCa, and increased cell proliferation, migration, apoptosis, and epithelial to mesenchymal transition (EMT) of cell(Chen *et al*., 2006; Yao *et al*., 2010). Recent studies reported that high expression of (GOLGA8B) is associated with PCa progression and poor prognosis(Cheng *et al*., 2020).

Although the number of FK866 treatment-induced upregulated genes were lower, the list of top genes included GABARAPL1, OTUD1, CHAC1, and SESN3. A recent study reported that elevated levels of GABARAPL1 suppress metastasis and cell proliferation through PI3K/Akt pathway in PCa(Su *et al*., 2017; C.-W. Xie *et al*., 2015). High expression levels of OTUD1 were associated with improved prognosis in non-small cell lung cancer and adenocarcinoma(J. Deng *et al*., 2019). While CHAC1 inhibits cell viability and increases the sensitivity to DTX for PCa(He *et al*., 2021), and is associated with autophagy marker ATF4(Crawford *et al*., 2015; Saavedra-García *et al*., 2021). We observed treatment-related downregulation of CHAC1 as well as ATF4. Further, Overexpression of SESN3 was associated with cancer cell proliferation suppression through mTORC1, Oxidative Stress, and Autophagy pathway(Sánchez-Álvarez *et al*., 2019). Additionally, we also observed FK866 treatment upregulated MERTK, APLF which are beneficial for treatment response against mCRPC. A recent study reported that MERTK regulates PCa dormancy, which inhibits PCa growth and increases metastasis-free survival(Cackowski *et al*., 2017). Further, APLF suppression promotes breast cancer and bladder cancer invasiveness through inhibition of proliferative capacity, altered cell cycle behavior, induced apoptosis, and impaired DNA repair ability(Majumder *et al*., 2018; Richter *et al*., 2019).

Traditional gene fusions are involved in the development of various cancer. DUS4L-BCAP29, a chimeric fusion RNA, has been reported to be a cancer-fusion in prostate and gastric cancer, which play a tumorigenic role(Kim *et al*., 2014; Tang *et al*., 2017; Wu *et al*., 2019). We observed >4.5 fold downregulation of DUS4L- BCAP29following FK866 treatment. Finally, our pathway analysis predicted mir-1-3p as the top upstream regulator based on DEGs. Interestingly, several recent studies have reported that miR-1-3p is a tumor suppressor and regulates PCa aggressiveness(Hudson *et al*., 2012; Kojima *et al*., 2012; S.-M. Li *et al*., 2018; Z.-C. Xie *et al*., 2018).

Importantly, we identified HES1 as one of the top genes downregulated following FK866 treatment. HES1 gene encodes Transcription factor HES1 (hairy and enhancer of split-1) protein. HES1 plays a critical role development of cancer stem cells (CSCs), cancer metastasis, and multidrug resistance in many cancers(Z.-H. Liu *et al*., 2015). Overexpression of HES1 PCa has been shown to play a crucial role in PCa progression(Carvalho *et al*., 2015; Soylu *et al*., 2016, 2021).

Cancer Metastasis is characterized by dissociation of cancer cells from the primary tumor site and colonization in a distant organ by traveling through interstitial tissues, intravasation into the blood or lymphatic vessels, and extravasation into a new site followed by proliferation and plasticity, a common feature of <u>EMT</u>(Van Zijl *et al*., 2011; Welch and Hurst, 2019). Thus, cell motility through confining pores plays a pivotal role in the process of metastatic dissemination, during which cells undergo EMT(Z. Liu *et al*., 2020; Y.-H. V. Ma *et al*., 2018; Tsai and Yang, 2013). Our innovative microfluidic-based approach using a physiologically relevant model of *in vitro* metastasis showed a significant decrease in migration efficiency of FK866-treated lethal PCa cells compared to control, confirming a potential role of FK866 in cell migration and tumor metastasis.

Since our approach is primarily focused on using *in vitro* model systems, further preclinical validation and single-cell multi-omics strategies using mouse xenograft models and patient-derived organoids are warranted to build upon our findings. Further, this will also allow the understanding of subclonal molecular pathways underlying differential patterns of PCa aggressiveness and drug response between AA vs. CA men.

Overall, our study creates a pipeline to introduce secDrugs as potent clinical-trial-ready therapeutic options for the management of lethal PCa with stem-like features.

## 4. MATERIALS AND METHODS

### Drugs and Reagents

Drugs, reagents, antibodies, and kits are listed in **Supplementary Table S4**.

### Identification of secondary drugs (secDrugs)

Design and development of the secDrug algorithm have been described earlier(Kumar *et al*., 2022). Briefly, we used a pharmacogenomics data-driven approach to identify potential agents that can be re-purposed as novel secondary drugs to treat cancers resistant to standard-of-care (primary) drugs when used in combination with the primary drug. As data source, we used the GDSC1000 (Genomics of Drug Sensitivity in Cancer) database, a large-scale pharmacogenomics database of dose-response results (IC_50_ or AUC) on 265 compounds in >1000 cell lines representing a wide spectrum of human cancers(W. Yang *et al*., 2013). These 265 drugs cover a wide range of targets and processes involved in cancer biology, which include drugs that are either approved and used in the clinic, or are undergoing clinical development, or in clinical trials, or are tool compounds in early phase development. For the purpose of this study in Prostate Cancer, we used inclusion criteria to filter cell lines with genito-urinal cancer subtypes. A total of 136 cell lines were selected from the GDSC1000 database breast (n=52), cervix (n=14), endometrium (n=11), ovary (n=45), prostate (n=8), testis (n=3), vulva (n=3).

First, we assumed that IC_50_ values of DTX in these lines (including PCa cell lines) were *S_bi_*:ε{1,…,*n*}, where there are *n* cell lines. Also, we assumed that there are *K* other drugs and the IC_50_ values of the n cell lines for the *K* drugs are given by:

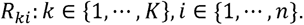

Next, we classified the cell lines as sensitive or resistant to DTX using a quantile of the empirical distribution of *R_ki_*, and a threshold criterion to achieve the classification. Finally, we identified secondary drugs or secDrugs that could kill the maximum number of DTX-resistant cell lines based on individual IC_50_ values. In the case of ties between the top secDrugs, we chose the drug with the lower mean IC_50_ values.

### Human Prostate Cancer Cell Lines

AR^lo^ mCRPC/NEPC (PC3, PC3M, DU145) and AR^high^ mCSPC (LnCAP, 22Rv1) cell lines, and the LnCAP-derived Androgen-independent, osteotropic subline C42b were obtained from the American Type Culture Collection (ATCC) (Manassas, VA, USA). The taxane-resistant cell lines PC3-TXR and DUTXR were generated using dose-escalation of taxanes over time, as described earlier(Takeda *et al*., 2007). The cell lines were authenticated at source and tested randomly at regular intervals for tissue specimen provenance and cell lineage at the AU Center for Pharmacogenomics and Single-Cell Omics (AUPharmGx) using GenePrint 24 System (Promega). All cell lines are mycoplasma negative. PC-3, PC-3M cells were maintained in 10% (v/v) (FBS) supplemented in F-12K, DU145 in Eagle’s Minimum Essential Medium (EMEM). PC3-TXR and DUTXR were maintained in RPMI-1640 media with 1% Penicillin-Streptomycin at 37°C, 21% O2, and 5% CO2 in a humidified cell culture chamber (Heracell™ VIOS 160i CO2; Thermo Scientific™).

### Patient Samples

Next-generation RNAseq data on PCa patient samples at the Atlanta VA Medical Center, Decatur, Georgia, Moffitt Cancer Center, Tampa, Florida) and the Sunnybrook Health Science Center (Toronto, ON) was obtained from the Gene expression omnibus database (GSE54460). Details on patient samples have been provided earlier(Long *et al*., 2014). Briefly, FFPE tissue blocks were obtained, following Institutional Review Board approvals, from patients at the Atlanta VA Medical Center, Decatur, Georgia, Moffitt Cancer Center, Tampa, Florida) and the Sunnybrook Health Science Center (Toronto, ON) next-generation RNAseq was performed on 100 patients. Demographic information (age, race), cancer staging (Gleason score, pSpTage), and biochemical recurrence were available on 100 patients. Among these cases, 49 had biochemical recurrence (BCR), and 51 had no BCR(Long *et al*., 2014).

Gene expression on PCa patients included in The Cancer Genome Atlas (TCGA) database was extracted from the Genomic Data Commons (GDCs) server (cancergenome.nih.gov). The interactive web-portals UALCAN and Gene Expression Profiling Interactive Analysis (GEPIA) were used for in-depth analysis of TCGA gene expression data files and to compare transcriptome data on target candidate pathway genes with tumor metastasis and patient survival from the prostate expression data matrix(Chandrashekar *et al*., 2017; Zheng *et al*., 2020).

### *In vitro* cytotoxicity assays and drug synergy analysis

*In vitro* chemo-sensitivity assays were performed on human PCa cell lines using mitochondrial enzyme activity or MTT (3-(4,5-dimethylthiazol-2-yl)-2,5-diphenyltetrazolium bromide reagent) assay. Briefly, cells were plated in a 96-well culture plate at 2×10^3^ cells/well and incubated for 24hr at 37°C with 5% CO_2_. Cells were treated with increasing concentrations of DTX (0- 2250nM), CBZ (0-2250nM), Enzalutamide (0-5062.5nM), Bicalutamide (0-5062.5nM) and FK866 (0- 625nM) as single agent, or the combination of DTX+FK866, and CBZ+FK866. For mCSPC lines, cells were also treated with Enzalutamide (ENZ) single-agent and ENZ+FK866 combination. Following 48 hour incubation, the tetrazolium dye MTT was added according to the manufacturer’s instructions, and absorbance was measured at 550nm using Synergy Neo2 Microplate Reader (BioTek, USA). Percent change relative to untreated controls was calculated at each drug concentration, and the effect of drug exposure was determined by constructing cytotoxicity (growth) curves. Half-maximal inhibitory drug concentration (IC_50_) values were estimated by nonlinear regression using a sigmoidal dose-response equation (variable slope). Drug synergy was calculated by comparing single-agent and combination drug-response data based on Chou-Talalay’s combination index (CI) method and the isobologram algorithm (CompuSyn software; Biosoft, US)(Chou, 2011). CI values between 0.9-0.3 and 0.3-0.1 signify synergism and strong synergism, respectively, between the drugs treated in combination.

### Caspase-3/7 activity assay

Cell death by apoptosis was measured using Caspase-Glo 3/7 luminescent assay system kit according to the manufacturer’s instructions (Promega Madison, WI). Briefly, 2×10^3^ cells/well were seeded into 96-well plates (triplicates) and treated at the estimated single-agent vs. combination IC_50_ values calculated by MTT assay. Following 48 hours of incubation, Caspase-Glo 3/7 reagent was added, incubated for 2h, and luminescence was measured using a Synergy Neo2 Microplate Reader (BioTek, USA). Apoptosis level in each treatment group was normalized to the control group (no drug treatment with baseline caspase 3/7 assay luminescence) for each cell line.

### Annexin V and propidium iodide (PI) staining

Annexin V and PI staining was used to assess apoptosis and necrosis by flow cytometry. Briefly, cells were seeded in 6 well plates at indicated concentrations and exposed to DTX and FK866 as a single agent and as combinations. After 48h, cells were labeled with binding buffer containing annexin V-FITC (25 µg/ml) and PI (25 µg/ml) as well as 10 mM HEPES, 140 mM NaCl, 5 mM KCl, 1 mM MgCl_2,_ and 1.8 mM CaCl_2_ (pH = 7.4), incubated for 10 min., followed by three washes in binding buffer. Both detached and attached cells were combined, and staining was quantified using a Becton Dickinson FACS Calibur flow cytometer (BD Biosciences, San Jose, CA) at 10,000 events per measurement.

### Assessment of cellular and nuclear morphology

To assess **<u>cellular morphology</u>**, PCa cells were seeded 0.025*10^6^ cells/ml in 6-well plates and exposed to FK866, either as a single agent or in combination with DTX for 48 h. Three areas with approximately equal cell densities were identified in each well, and images were captured with an EVOS FL digital cell imaging system (Thermo Fisher Scientific, Inc.) using a 10XLobjective.

For **<u>nuclear morphology</u>**, PCa cells were plated on top of the glass coverslip (1.5*10^5^ cells/ml), incubated overnight, and treated with either vehicle or FK866 alone or as a combination with DTX. After 48h, the cells were labeled with NucBlue Live reagent and incubated for 20 minutes. Images were captured using Nikon Eclipse Ti2 microscope and recorded in bright field and phase contrast modes at 20X and 40X magnifications. Images were analyzed using Image J software (National Institutes of Health, Bethesda, MD, USA).

### NAD/NADH Activity Assay

PCa cell lines (PC-3, PC-3M, DU145, and DUTXR) were plated in a white-welled luminometer 96-well plate at a seeding density of 2×10^3^ cells/well and treated with indicated concentrations of FK866 as a single agent. Following 48hr incubation, NAD cycling enzyme, NAD Cycling Substrate, and NAD/NADH-Glo Detection Reagent were added to the cells and incubated for 1 h. Luminescence was measured using a Synergy Neo2 plate reader (BioTek, USA).

### Assessment of cell cycle

Control (no drug) and post-treated cells were prepared for cell cycle analysis by staining with PI (50 µg/ml) in sample buffer [PBS + 1% (w/v) glucose], containing RNAse A (100 units/ml) for 30 min at room temperature and analyzed by flow cytometry using a Becton Dickinson FACS Calibur flow cytometer (BD Biosciences, San Jose, CA). Cell cycle data were analyzed using CytExpert (Beckman Coulter Inc, Indianapolis, IN). Data are presented as the mean ± SEM of three separate experiments (n = 3/study).

### Assessment of intracellular ROS levels (DCFDA assay)

Cells were plated at a seeding density of 2000 cells/well and incubated overnight at 37°C. After 24h, 100 ul of 10 uM DCFDA solution was added to each well and incubated in the dark for 45 minutes at 37°C. DCFDA solution was then discarded and treated with either vehicle (0.5% DMSO) or FK866 single agent and in combination with DTX. Samples were collected at different time points (2, 4, 8, and 24h). Fluorescent intensity was measured on Synergy Neo2 Hybrid Multi-Mode Microplate Reader, BioTek (Winooski, VT, USA) at excitation - 485nM and emission - 535 nm in endpoint mode.

### Assessment of Mitochondrial Membrane Potential

Cell lines were treated with DTX, FK866, and DTX+FK866. Further, 100 μL/well of working JC-1 solution was added to the plate and incubated at 37°C for 10 minutes in the dark. Read plate endpoint in the presence of compounds, media on a fluorescent plate reader Synergy Neo2 Hybrid Multi-Mode Microplate Reader, BioTek (Winooski, VT, USA) at 535 nm.

### Assessment of Side Population

A total of 1×10^6^ /ml cells were cultured in 6 well plates and treated with FK866 alone or in combination with DTX. After 24h, cells were stained with 5 μM Vybrant DyeCycle Violet and 1 ug of 7-AAD for 30 min at 37°C. Following dye incubation, cells were immediately analyzed (10,000 events per measurement) using a Becton Dickinson FACS Calibur flow cytometer (BD Biosciences, San Jose, CA).

### Colony formation assay

PCa cells were seeded in a 6-well plate at 0.025*10^6^ cells/ml, incubated overnight, and treated with DTX and FK866 as a single agent or in combination. The cells were then harvested and plated in a 24-well plate at a concentration of 1000 cells/well and incubated for 1- 2 weeks. The colonies were fixed with 100% methanol and stained with Crystal Violet. Images were taken for control, treated cells, and the colonies using an EVOS FL digital cell imaging system (Thermo Fisher Scientific, Inc.). Images were recorded in bright field and phase contrast modes at 20X and 40X magnifications and analyzed using Image J software.

### Cell migration/Scratch Assay

Cells were plated in 6-well plates at 1×10^5^ cells/well and incubated for 48hr to a 95% confluency. The monolayer was scratched with a SPLScar Scratcher 6 well Tip at a width of 0.50 mm at the center of the well. FK866 as single-agent or DTX+FK866 combination doses were applied to the cells in the respective wells. F- 12K culture medium supplemented with 10% FBS containing the vehicle (0.05 % DMSO) was added to the cells in the control wells. Micrographs of the wound areas were obtained at 0, 24, and 48hr using an EVOS FL digital cell imaging system (Thermo Fisher Scientific, Inc.). Images were recorded in brightfield and phase contrast modes at 20X and 40X magnifications. The area of the initial wound (at 0 h) and the “gap area” were measured at 48hr with Image J software.

### Microfluidic (μ)-channel Cell Migration Assay

The fabrication of a Polydimethylsiloxane (PDMS)-based μ-channel assay using standard multilayer photolithography and replica molding have been demonstrated earlier (**Figure S3**)(Wisniewski *et al*., 2020). In this study, PCa cells were seeded in 6 well plates exposed to FK866 as a single agent and FK866+DTX combinations at indicated concentrations. Next, 1-1.5 x 10^5^ cells were introduced into the cell seeding inlet line of the microfluidic channel via pressure-driven flow and were allowed to adhere for 30 min at 37°C, 5% CO_2_. Next, the cell suspension was removed and substituted with a serum-free medium. Medium supplemented with 10% FBS was added into the chemoattractant inlet line to trigger cell entry into the channels. The devices were placed on an automated Nikon Ti2 Inverted Microscope equipped with a Tokai Stage-Top incubator unit, which maintained cells at 37 °C and 5% CO_2_. Cell entry into the channels was recorded via time-lapse microscopy. Images were recorded every 20 min for 10hr with a 10x /0.45 NA Ph1 objective.

### Pre- and post-treatment tumor mRNA sequencing (RNAseq)

The effects of DTX and FK866 as a single agent and in combination exposure on gene expression in PCa cell lines were assessed using next-generation RNAseq of bulk tumor cells. Pre- and post- drug-exposure FK866 single-agent, TX+FK866 combination) tumor cells were harvested, and high-quality RNA will be extracted using QIAshredder and RNeasy kit (Qiagen) according to the manufacturer’s protocol. RNA concentration and integrity were assessed using Nanodrop-8000 spectrophotometer (Thermo Scientific, USA), Qubit 2.0 Fluorometer (Invitrogen, Carlsbad, CA, USA), and Agilent 2100 Bioanalyzer (Applied Biosystems, Carlsbad, CA, USA) and stored at -80°C. An RNA integrity number (RIN) threshold >8 was applied, and RNAseq libraries were constructed using Illumina TruSeq RNA Sample Preparation kit v2. Libraries were then size-selected to generate inserts of ∼200 bp, and RNAseq was performed on Illumina’s NovaSeq platform using a 150bp paired-end protocol with a depth of >20million reads per sample. Average quality scores were thoroughly above Q30 for all libraries in both R1 and R2.

### RNAseq data analysis

RNAseq data from the cell lines and patient RNAseq data (described above) was pre-processed and normalized, and differential expression (DE) analysis was performed using command-line based analysis pipeline (DEseq2 and edgeR) and Partek Flow software (Partek, Inc, USA). Quality control (QC) check on the RNAseq raw reads was performed using the FastQC tool, followed by read-trimming to remove base positions that have a low median (or bottom quartile) score. STAR Aligner tool mapped processed RNAseq reads to the hg38 human genome build. Next, read counts were CPM-normalized, and then we used GSA (Gene-specific analysis) based on limma trend that applies an empirical Bayesian method to perform differential gene expression analysis between groups and detect the DE genes. Genes with mean fold-change>|1| and p<0.05 were considered as the threshold for reporting significant differential gene expression. Heatmaps were generated using unsupervised hierarchical clustering (HC) analysis based on the differentially expressed genes (DEGs).

### Pre- and post-treatment Single-cell RNA sequencing (scRNAseq)

Automated single-cell capture and cDNA synthesis were performed on the untreated and FK866-treated acquired taxane resistant mCRPC DUTXR using the 10X Genomics Chromium platform. Single-cell RNAseq-based gene expression analysis will be performed on Illumina HiSeq 2500 NGS platform (Paired- end. 2*125bp, 100 cycles. v3 chemistry) at ∼10 million reads per sample.

### ScRNAseq data analysis

Single-cell RNAseq datasets were obtained as matrices in the Hierarchical Data Format (HDF5 or H5). We used CellRanger, Seurat, and Partek Flow software packages will be used to pre-process the scRNAseq data and perform single-cell transcriptomics. Highly variable genes were selected for clustering analysis based on a graph-based clustering approach. The visualization of cell populations was performed by T-distributed stochastic neighbor embedding (t-SNE) and UMAP (Uniform Manifold Approximation and Projection) for biomarker-based identification of subclones representing TX-resistant cells, potential FK866 target subclones, and cancer stem-cell signatures, as well as FK866 treatment-induced erosion of these subclones.

### Ingenuity pathway analysis (IPA)

Ingenuity pathway analysis (IPA; Qiagen) analysis was performed using top DEGs to reveal molecular pathways/mechanisms, upstream regulator molecules, downstream effects, biological processes, and predicted causal networks governing FK866 function and successful drug combinations in AVPC(Krämer *et al*., 2014).

### Immunoblotting

PCa cells were seeded and exposed to drugs at the indicated concentrations of each treatment protocol. Post-treatment cells were lysed in cell lysis buffer (Thermo Scientific RIPA Lysis and Extraction Buffer). Quantification of proteins was performed using Bradford assay, and a calibration curve of protein content was created from the BSA protein standard kit (Bio-Rad Laboratories, CA). An equal amount of protein (50 ng) was loaded onto 4-15% Criterion TGX Stain-Free Precast Gels. Proteins were separated under reducing conditions and then transferred to a PVDF membrane using Fisherbrand Semidry Blotting Apparatus. Nonspecific binding was limited by incubating the membrane in blocking buffer (2.5% (w/v) casein, pH 7.6, 150mM NaCl, 10mM TRIS-HCl, and 0.02% sodium azide). Membranes were incubated overnight with primary antibodies for targeted gene/protein (1:1000) and then with the appropriate secondary antibody (1:10,000) for 1.5hr at room temperature. Immunoreactivity was detected using Pierce ECL Western Blotting substrate (Bio-Rad, CA). Images were captured and quantified by Gel Doc EZ Gel Documentation System and ImageLab Software (Hercules CA, USA). Densitometry analysis was performed using the standard image analysis software Image J.

### Statistical analysis

All statistical analysis was performed using R (the project for statistical computing and graphics) version 4.1.0 and GraphPad Prism v9.0. All tests were two-sided and p<0.05 to be considered statistically significant. We used a non-parametric Wilcoxon rank-sum test for differential expression analysis between two groups of cells.

## AUTHORSHIP STATEMENT

All authors have read the journal’s authorship statement.

## DATA AVAILABILITY STATEMENT

The datasets generated during and/or analyzed during the current study are available from the corresponding author on reasonable request.

## AUTHOR CONTRIBUTION

SM, TMG, UKM, CY, and AKM participated in research design; SM, TMG, SC, FH, FA, and ASA conducted experiments: SM, TMG, UKM, PM, IE, and AKM performed data analysis; FL helped with the creation of TX- resistant cell lines. SM, TMG, PM, WDC, CY, RDA, and AKM wrote or contributed to the writing of the manuscript; AKM supervised the overall project.

## REFERENCES

1. Audrito V (2020) The dual face of NAMPT: Intracellular/extracellular protein and diagnostic/therapeutic target in cancer. EBioMedicine 62, 103109.

2. Bowlby SC, Thomas MJ, D’Agostino RB, and Kridel SJ (2012) Nicotinamide phosphoribosyl transferase (Nampt) is required for de novo lipogenesis in tumor cells. PLoS One 7, e40195.

3. Cackowski FC, Eber MR, Rhee J, Decker AM, Yumoto K, Berry JE, … Taichman RS (2017) Mer Tyrosine Kinase Regulates Disseminated Prostate Cancer Cellular Dormancy. J Cell Biochem 118, 891–902.

4. Cao Z, Xu Y, Guo F, Chen X, Ji J, Xu H, … Wang F (2020) FASN Protein Overexpression Indicates Poor Biochemical Recurrence-Free Survival in Prostate Cancer. Dis Markers 2020, 3904947.

5. Carvalho FLF, Marchionni L, Gupta A, Kummangal BA, Schaeffer EM, Ross AE, and Berman DM (2015) HES6 promotes prostate cancer aggressiveness independently of Notch signalling. J Cell Mol Med 19, 1624–36.

6. Chandrashekar DS, Bashel B, Balasubramanya SAH, Creighton CJ, Ponce-Rodriguez I, Chakravarthi BVSK, and Varambally S (2017) UALCAN: A Portal for Facilitating Tumor Subgroup Gene Expression and Survival Analyses. Neoplasia 19, 649–658.

7. Chen Q, Watson JT, Marengo SR, Decker KS, Coleman I, Nelson PS, and Sikes RA (2006) Gene expression in the LNCaP human prostate cancer progression model: progression associated expression in vitro corresponds to expression changes associated with prostate cancer progression in vivo. Cancer Lett 244, 274–88.

8. Cheng Y, Li L, Qin Z, Li X, and Qi F (2020) Identification of castration-resistant prostate cancer-related hub genes using weighted gene co-expression network analysis. J Cell Mol Med 24, 8006–8017.

9. Chou TC (2011) The mass-action law based algorithm for cost-effective approach for cancer drug discovery and development Am J Cancer Res 1, 925–954.

10. Conteduca V, Oromendia C, Eng KW, Bareja R, Sigouros M, Molina A, … Beltran H (2019) Clinical features of neuroendocrine prostate cancer. Eur J Cancer 121, 7–18.

11. Contreras HR, López-Moncada F, and Castellón EA (2020) Cancer stem cell and mesenchymal cell cooperative actions in metastasis progression and hormone resistance in prostate cancer: Potential role of androgen and gonadotropinLreleasing hormone receptors (Review). Int J Oncol 56, 1075–1082.

12. Cornford P, Bellmunt J, Bolla M, Briers E, De Santis M, Gross T, … Mottet N (2017) EAU-ESTRO-SIOG Guidelines on Prostate Cancer. Part II: Treatment of Relapsing, Metastatic, and Castration-Resistant Prostate Cancer. Eur Urol 71, 630–642.

13. Crawford RR, Prescott ET, Sylvester CF, Higdon AN, Shan J, Kilberg MS, and Mungrue IN (2015) Human CHAC1 Protein Degrades Glutathione, and mRNA Induction Is Regulated by the Transcription Factors ATF4 and ATF3 and a Bipartite ATF/CRE Regulatory Element. J Biol Chem 290, 15878–15891.

14. Cronin KA, Lake AJ, Scott S, Sherman RL, Noone A-M, Howlader N, … Jemal A (2018) Annual Report to the Nation on the Status of Cancer, part I: National cancer statistics. Cancer 124, 2785–2800.

15. De Bono JS, Oudard S, Ozguroglu M, Hansen S, MacHiels JP, Kocak I, … Sartor AO (2010) Prednisone plus cabazitaxel or mitoxantrone for metastatic castration-resistant prostate cancer progressing after docetaxel treatment: A randomised open-label trial Lancet 376, 1147–1154.

16. Deng J, Hou G, Fang Z, Liu J, and Lv X-D (2019) Distinct expression and prognostic value of OTU domain-containing proteins in non-small-cell lung cancer. Oncol Lett 18, 5417–5427.

17. Deng L, Gu X, Zeng T, Xu F, Dong Z, Liu C, and Chao H (2019) Identification and characterization of biomarkers and their functions for docetaxelLresistant prostate cancer cells Oncol Lett.

18. Galletti G, Leach BI, Lam L, and Tagawa ST (2017) Mechanisms of resistance to systemic therapy in metastatic castration-resistant prostate cancer. Cancer Treat Rev 57, 16–27.

19. Galli U, Colombo G, Travelli C, Tron GC, Genazzani AA, and Grolla AA (2020) Recent Advances in NAMPT Inhibitors: A Novel Immunotherapic Strategy. Front Pharmacol 11, 656.

20. Gao S, Lee P, Wang H, Gerald W, Adler M, Zhang L, … Wang Z (2005) The androgen receptor directly targets the cellular Fas/FasL-associated death domain protein-like inhibitory protein gene to promote the androgen-independent growth of prostate cancer cells. Mol Endocrinol 19, 1792–802.

21. Gutierrez LS, Noria F, Finol H, Sun L, Castellino F, and Pollard M (2005) Fas ligand expression and its correlation with apoptosis and proliferation in Lobund-Wistar prostate carcinomas. Pathobiology 72, 260– 8.

22. Hamdy FC, Donovan JL, Lane JA, Mason M, Metcalfe C, Holding P, … ProtecT Study Group (2016) 10-Year Outcomes after Monitoring, Surgery, or Radiotherapy for Localized Prostate Cancer. N Engl J Med 375, 1415–1424.

23. He S, Zhang M, Ye Y, Zhuang J, Ma X, Song Y, and Xia W (2021) ChaC glutathione specific γ- glutamylcyclotransferase 1 inhibits cell viability and increases the sensitivity of prostate cancer cells to docetaxel by inducing endoplasmic reticulum stress and ferroptosis. Exp Ther Med 22, 997.

24. Heske CM (2019) Beyond Energy Metabolism: Exploiting the Additional Roles of NAMPT for Cancer Therapy. Front Oncol 9, 1514.

25. Howlader N, Noone AM, Krapcho M, Miller D, Bishop K, Altekruse SF, … Tatalovich Z (2015) SEER cancer statistics review, 1975–2013, National Cancer Institute Bethesda, MD Http//SeerCancerGov/Csr/1975_2013/Based Novemb.

26. Hudson RS, Yi M, Esposito D, Watkins SK, Hurwitz AA, Yfantis HG, … Ambs S (2012) MicroRNA-1 is a candidate tumor suppressor and prognostic marker in human prostate cancer. Nucleic Acids Res 40, 3689–703.

27. Kapoor A, Wu C, Shayegan B, and Rybak AP (2016) Contemporary agents in the management of metastatic castration-resistant prostate cancer *J Can Urol Assoc* **10**, E414–E423.

28. Kim H-P, Cho G-A, Han S-W, Shin J-Y, Jeong E-G, Song S-H, … Kim T-Y (2014) Novel fusion transcripts in human gastric cancer revealed by transcriptome analysis. Oncogene 33, 5434–41.

29. Kojima S, Chiyomaru T, Kawakami K, Yoshino H, Enokida H, Nohata N, … Seki N (2012) Tumour suppressors miR-1 and miR-133a target the oncogenic function of purine nucleoside phosphorylase (PNP) in prostate cancer. Br J Cancer 106, 405–13.

30. Krämer A, Green J, Pollard J, and Tugendreich S (2014) Causal analysis approaches in Ingenuity Pathway Analysis. Bioinformatics 30, 523–30.

31. Kumar H, Mazumder S, Chakravarti S, Sharma N, Mukherjee UK, Kumar S, … Mitra AK (2022) secDrug: a pipeline to discover novel drug combinations to kill drug-resistant multiple myeloma cells using a greedy set cover algorithm and single-cell multi-omics Blood Cancer J 12, 39.

32. Larré S, Tran N, Fan C, Hamadeh H, Champigneulles J, Azzouzi R, … Olivier JL (2008) PGE2 and LTB4 tissue levels in benign and cancerous prostates. Prostaglandins Other Lipid Mediat 87, 14–9.

33. Li P, Yang R, and Gao W-Q (2014) Contributions of epithelial-mesenchymal transition and cancer stem cells to the development of castration resistance of prostate cancer. Mol Cancer 13, 55.

34. Li S-M, Wu H-L, Yu X, Tang K, Wang S-G, Ye Z-Q, and Hu J (2018) The putative tumour suppressor miR-1-3p modulates prostate cancer cell aggressiveness by repressing E2F5 and PFTK1. J Exp Clin Cancer Res 37, 219.

35. Liu X, Chen L, Fan Y, Hong Y, Yang X, Li Y, … Xu D (2019) IFITM3 promotes bone metastasis of prostate cancer cells by mediating activation of the TGF-β signaling pathway. Cell Death Dis 10, 517.

36. Liu Z-H, Dai X-M, and Du B (2015) Hes1: a key role in stemness, metastasis and multidrug resistance. Cancer Biol Ther 16, 353–9.

37. Liu Z, Lee SJ, Park S, Konstantopoulos K, Glunde K, Chen Y, and Barman I (2020) Cancer cells display increased migration and deformability in pace with metastatic progression FASEB J 34, 9307–9315.

38. Long Q, Xu J, Osunkoya AO, Sannigrahi S, Johnson BA, Zhou W, … Moreno CS (2014) Global transcriptome analysis of formalin-fixed prostate cancer specimens identifies biomarkers of disease recurrence. Cancer Res 74, 3228–37.

39. Ma C, Ci M, Yoshioka M, Mayumi Y, Boivin A, André B, … Jonny S-A (2008) Prostate-specific genes and their regulation by dihydrotestosterone. Prostate 68, 241–54.

40. Ma Y-HV, Middleton K, You L, and Sun Y (2018) A review of microfluidic approaches for investigating cancer extravasation during metastasis Microsystems Nanoeng 4, 1–13.

41. Magrini E, and Garlanda C (2021) Noncanonical Functions of C1s Complement Its Canonical Functions in Renal Cancer. Cancer Immunol Res 9, 855.

42. Majumder A, Syed KM, Mukherjee A, Lankadasari MB, Azeez JM, Sreeja S, … Dutta D (2018) Enhanced expression of histone chaperone APLF associate with breast cancer. Mol Cancer 17, 76.

43. Maldonado LY, Arsene D, Mato JM, and Lu SC (2018) Methionine adenosyltransferases in cancers: Mechanisms of dysregulation and implications for therapy. Exp Biol Med (Maywood) 243, 107–117.

44. Mistriotis P, Wisniewski EO, Bera K, Keys J, Li Y, Tuntithavornwat S, … Konstantopoulos K (2019) Confinement hinders motility by inducing RhoA-mediated nuclear influx, volume expansion, and blebbing. J Cell Biol 218, 4093–4111.

45. Munkley J, Maia TM, Ibarluzea N, Livermore KE, Vodak D, Ehrmann I, … Elliott DJ (2018) Androgen-dependent alternative mRNA isoform expression in prostate cancer cells. F1000Research 7, 1189.

46. Nacarelli T, Fukumoto T, Zundell JA, Fatkhutdinov N, Jean S, Cadungog MG, … Zhang R (2020) NAMPT Inhibition Suppresses Cancer Stem-like Cells Associated with Therapy-Induced Senescence in Ovarian Cancer. Cancer Res 80, 890–900.

47. Ouyang X, Jessen WJ, Al-Ahmadie H, Serio AM, Lin Y, Shih W-J, … Abate-Shen C (2008) Activator protein-1 transcription factors are associated with progression and recurrence of prostate cancer. Cancer Res 68, 2132–44.

48. Paul CD, Mistriotis P, and Konstantopoulos K (2017) Cancer cell motility: lessons from migration in confined spaces. Nat Rev Cancer 17, 131–140.

49. Petrylak DP, Tangen CM, Hussain MHA, Lara PN, Jones JA, Taplin ME, … Crawford ED (2004) Docetaxel and estramustine compared with mitoxantrone and prednisone for advanced refractory prostate cancer N Engl J Med 351, 1513–1520.

50. Rajapaksa US, Jin C, and Dong T (2020) Malignancy and IFITM3: Friend or Foe? Front Oncol 10, 593245.

51. Richter C, Marquardt S, Li F, Spitschak A, Murr N, Edelhäuser BAH, … Logotheti S (2019) Rewiring E2F1 with classical NHEJ via APLF suppression promotes bladder cancer invasiveness. J Exp Clin Cancer Res 38, 292.

52. Riedel M, Cai H, Stoltze IC, Vendelbo MH, Wagner EF, Bakiri L, and Thomsen MK (2021) Targeting AP-1 transcription factors by CRISPR in the prostate. Oncotarget 12, 1956–1961.

53. Saad F, and Miller K (2014) Treatment options in castration-resistant prostate cancer: Current therapies and emerging docetaxel-based regimens Urol Oncol Semin Orig Investig 32, 70–79.

54. Saavedra-García P, Roman-Trufero M, Al-Sadah HA, Blighe K, López-Jiménez E, Christoforou M, … Auner HW (2021) Systems level profiling of chemotherapy-induced stress resolution in cancer cells reveals druggable trade-offs. Proc Natl Acad Sci U S A 118.

55. Sánchez-Álvarez M, Strippoli R, Donadelli M, Bazhin A V, and Cordani M (2019) Sestrins as a Therapeutic Bridge between ROS and Autophagy in Cancer. Cancers (Basel) 11.

56. Sartor AO (2011) Progression of metastatic castrate-resistant prostate cancer: Impact of therapeutic intervention in the post-docetaxel space J Hematol Oncol 4, 1–7.

57. Sauer H, Kampmann H, Khosravi F, Sharifpanah F, and Wartenberg M (2021) The nicotinamide phosphoribosyltransferase antagonist FK866 inhibits growth of prostate tumour spheroids and increases doxorubicin retention without changes in drug transporter and cancer stem cell protein expression. Clin Exp Pharmacol Physiol 48, 422–434.

58. Scher HI, Fizazi K, Saad F, Taplin M-E, Sternberg CN, Miller K, … AFFIRM Investigators (2012) Increased survival with enzalutamide in prostate cancer after chemotherapy. N Engl J Med 367, 1187–97.

59. Scher HI, Halabi S, Tannock I, Morris M, Sternberg CN, Michael a, … Eisenberger a (2014) NIH Public Access Progressive Prostate Cancer and Castrate Levels of Trials Working Group 26, 1148–1159.

60. Scher HI, Solo K, Valant J, Todd MB, and Mehra M (2015) Prevalence of Prostate Cancer Clinical States and Mortality in the United States: Estimates Using a Dynamic Progression Model. PLoS One 10, e0139440.

61. Shafi AA, Yen AE, and Weigel NL (2013) Androgen receptors in hormone-dependent and castration-resistant prostate cancer. Pharmacol Ther 140, 223–38.

62. Shankar E, Song K, Corum SL, Bane KL, Wang H, Kao H-Y, and Danielpour D (2016) A Signaling Network Controlling Androgenic Repression of c-Fos Protein in Prostate Adenocarcinoma Cells. J Biol Chem 291, 5512–5526.

63. Song Y, Wang M, Tong H, Tan Y, Hu X, Wang K, and Wan X (2021) Plasma exosomes from endometrial cancer patients contain LGALS3BP to promote endometrial cancer progression. Oncogene 40, 633–646.

64. Soylu H, Acar N, Ozbey O, Unal B, Koksal IT, Bassorgun I, … Ustunel I (2016) Characterization of Notch Signalling Pathway Members in Normal Prostate, Prostatic Intraepithelial Neoplasia (PIN) and Prostatic Adenocarcinoma. Pathol Oncol Res 22, 87–94.

65. Soylu H, Kırca M, Avcı S, Ozpolat B, and Ustunel I (2021) Antiandrogen abiraterone and docetaxel treatments affect Notch1, Jagged1 and Hes1 expressions in metastatic prostate cancer cells. Exp Mol Pathol 119, 104607.

66. Su W, Li S, Chen X, Yin L, Ma P, Ma Y, and Su B (2017) GABARAPL1 suppresses metastasis by counteracting PI3K/Akt pathway in prostate cancer. Oncotarget 8, 4449–4459.

67. Takeda M, Mizokami A, Mamiya K, Li YQ, Zhang J, Keller ET, and Namiki M (2007) The establishment of two paclitaxel-resistant prostate cancer cell lines and the mechanisms of paclitaxel resistance with two cell lines. Prostate 67, 955–67.

68. Tan B, Young DA, Lu Z-H, Wang T, Meier TI, Shepard RL, … Zhao G (2013) Pharmacological inhibition of nicotinamide phosphoribosyltransferase (NAMPT), an enzyme essential for NAD+ biosynthesis, in human cancer cells: metabolic basis and potential clinical implications. J Biol Chem 288, 3500–11.

69. Tang Y, Qin F, Liu A, and Li H (2017) Recurrent fusion RNA DUS4L-BCAP29 in non-cancer human tissues and cells. Oncotarget 8, 31415–31423.

70. Tannock IF, de Wit R, Berry WR, Horti J, Pluzanska A, Chi KN, … TAX 327 Investigators (2004) Docetaxel plus prednisone or mitoxantrone plus prednisone for advanced prostate cancer. N Engl J Med 351, 1502– 12.

71. Theodoros Karantanos, Paul G. Corn and TCT (2013) Prostate cancer progression after androgen deprivation therapy: mechanisms of castrate-resistance and novel therapeutic approaches Theodoros Curr Opin Support Palliat Care 7, 258–264.

72. Toraih EA, Fawzy MS, El-Falouji AI, Hamed EO, Nemr NA, Hussein MH, and Abd El Fadeal NM (2016) Stemness-related transcriptional factors and homing gene expression profiles in hepatic differentiation and cancer. Mol Med 22, 653–663.

73. Tsai JH, and Yang J (2013) Epithelial-mesenchymal plasticity in carcinoma metastasis Genes Dev 27, 2192–2206.

74. Van Zijl F, Krupitza G, and Mikulits W (2011) Initial steps of metastasis: Cell invasion and endothelial transmigration Mutat Res - Rev Mutat Res 728, 23–34.

75. Vashchenko N, and Abrahamsson P-A (2005) Neuroendocrine differentiation in prostate cancer: implications for new treatment modalities. Eur Urol 47, 147–55.

76. Wadosky KM, and Koochekpour S (2016) Molecular mechanisms underlying resistance to androgen deprivation therapy in prostate cancer Oncotarget 7, 64447–64470.

77. Wang B, Hasan MK, Alvarado E, Yuan H, Wu H, and Chen WY (2011) NAMPT overexpression in prostate cancer and its contribution to tumor cell survival and stress response. Oncogene 30, 907–21.

78. Wang Y, Chen J, Wu Z, Ding W, Gao S, Gao Y, and Xu C (2021) Mechanisms of enzalutamide resistance in castration-resistant prostate cancer and therapeutic strategies to overcome it. Br J Pharmacol 178, 239– 261.

79. Weigelin B, Bakker G-J, and Friedl P (2012) Intravital third harmonic generation microscopy of collective melanoma cell invasion: Principles of interface guidance and microvesicle dynamics. Intravital 1, 32–43.

80. Welch DR, and Hurst DR (2019) Defining the Hallmarks of Metastasis Cancer Res 79, 3011–3027.

81. Wisniewski EO, Mistriotis P, Bera K, Law RA, Zhang J, Nikolic M, … Konstantopoulos K (2020) Dorsoventral polarity directs cell responses to migration track geometries. Sci Adv 6, eaba6505.

82. Wong BS, Shea DJ, Mistriotis P, Tuntithavornwat S, Law RA, Bieber JM, … Konstantopoulos K (2019) A Direct Podocalyxin-Dynamin-2 Interaction Regulates Cytoskeletal Dynamics to Promote Migration and Metastasis in Pancreatic Cancer Cells. Cancer Res 79, 2878–2891.

83. Wu H, Li X, and Li H (2019) Gene fusions and chimeric RNAs, and their implications in cancer. Genes Dis 6, 385–390.

84. Xie C-W, Zhou Y, Liu S-L, Fang Z-Y, Su B, and Zhang W (2015) Gabarapl1 mediates androgen-regulated autophagy in prostate cancer. Tumour Biol 36, 8727–33.

85. Xie Z-C, Huang J-C, Zhang L-J, Gan B-L, Wen D-Y, Chen G, … Yan H-B (2018) Exploration of the diagnostic value and molecular mechanism of miR-1 in prostate cancer: A study based on meta-analyses and bioinformatics. Mol Med Rep 18, 5630–5646.

86. Yang M, Gao H, Chen P, Jia J, and Wu S (2013) Knockdown of interferon-induced transmembrane protein 3 expression suppresses breast cancer cell growth and colony formation and affects the cell cycle. Oncol Rep 30, 171–8.

87. Yang P, Zhang L, Shi Q-J, Lu Y-B, Wu M, Wei E-Q, and Zhang W-P (2015) Nicotinamide phosphoribosyltransferase inhibitor APO866 induces C6 glioblastoma cell death via autophagy. Pharmazie 70, 650–5.

88. Yang W, Soares J, Greninger P, Edelman EJ, Lightfoot H, Forbes S, … Garnett MJ (2013) Genomics of Drug Sensitivity in Cancer (GDSC): a resource for therapeutic biomarker discovery in cancer cells. Nucleic Acids Res 41, D955–61.

89. Yao S, Bee A, Brewer D, Dodson A, Beesley C, Ke Y, … Foster CS (2010) PRKC-ζ Expression Promotes the Aggressive Phenotype of Human Prostate Cancer Cells and Is a Novel Target for Therapeutic Intervention. Genes Cancer 1, 444–64.

90. Zhao B, Wang H, Zong G, and Li P (2013) The role of IFITM3 in the growth and migration of human glioma cells. BMC Neurol 13, 210.

91. Zheng H, Zhang G, Zhang L, Wang Q, Li H, Han Y, … Guo X (2020) Comprehensive Review of Web Servers and Bioinformatics Tools for Cancer Prognosis Analysis. Front Oncol 10, 68.

92. Zhou J, Wang H, Cannon V, Wolcott KM, Song H, and Yates C (2011) Side population rather than CD133(+) cells distinguishes enriched tumorigenicity in hTERT-immortalized primary prostate cancer cells. Mol Cancer 10, 112.

