## Supplementary Figures for "Integrating pharmacogenomics data-driven prediction with bulk and single-cell RNAseq to demonstrate the efficacy of an NAMPT inhibitor against aggressive, taxane-resistant, and stem-like cells in lethal prostate cancer"

### Slide 1
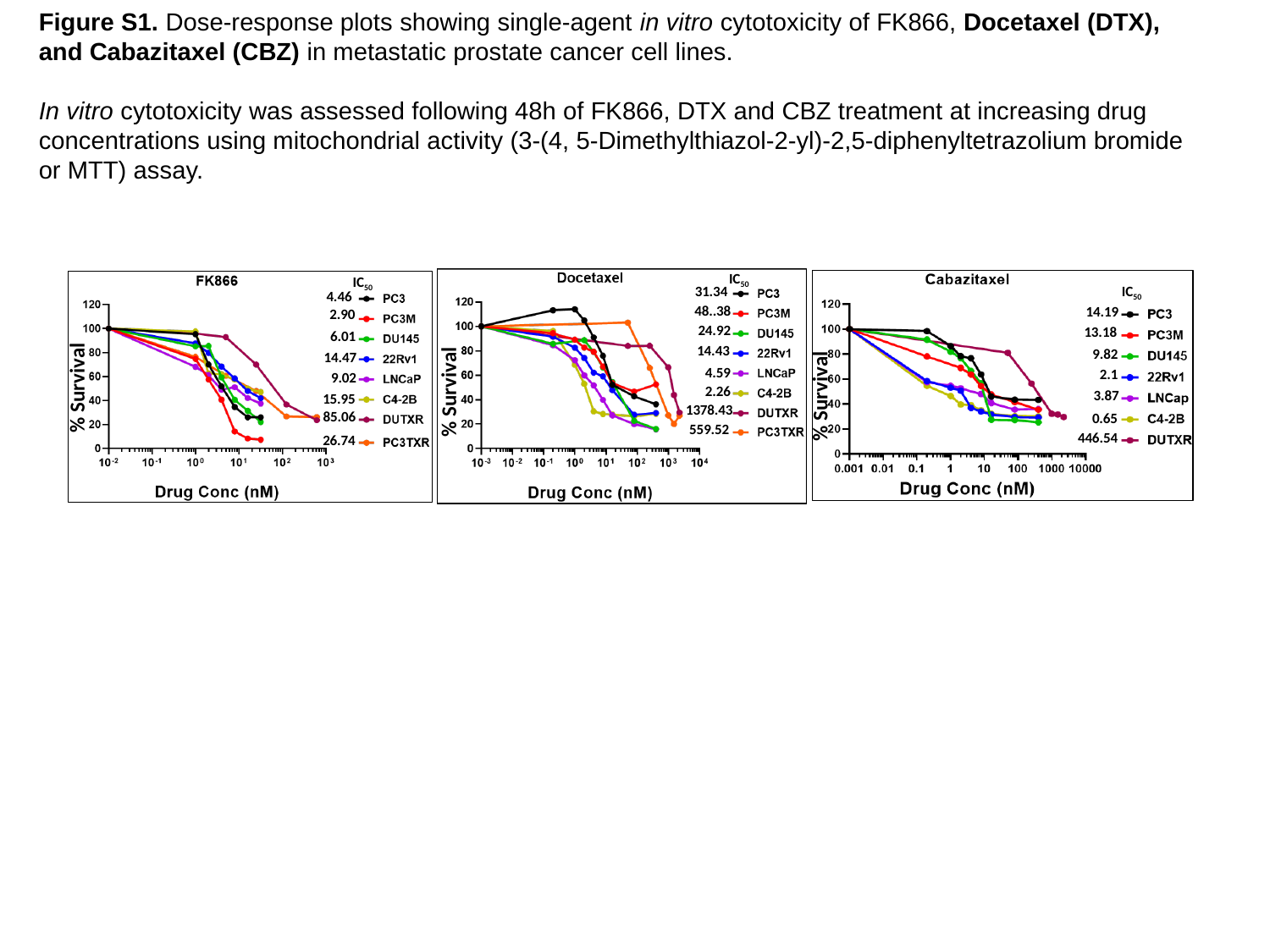

Figure S1. Dose-response plots showing single-agent in vitro cytotoxicity of FK866, Docetaxel (DTX), and Cabazitaxel (CBZ) in metastatic prostate cancer cell lines.
In vitro cytotoxicity was assessed following 48h of FK866, DTX and CBZ treatment at increasing drug concentrations using mitochondrial activity (3-(4, 5-Dimethylthiazol-2-yl)-2,5-diphenyltetrazolium bromide or MTT) assay.
IC_50_
31.34
48..38
24.92
14.43
4.59
% Survival
2.26
1378.43
559.52
IC_50_
4.46
2.90
6.01
14.47
9.02
% Survival
15.95
85.06
26.74
IC_50_
14.19
13.18
9.82
2.1
% Survival
3.87
0.65
446.54

### Slide 2
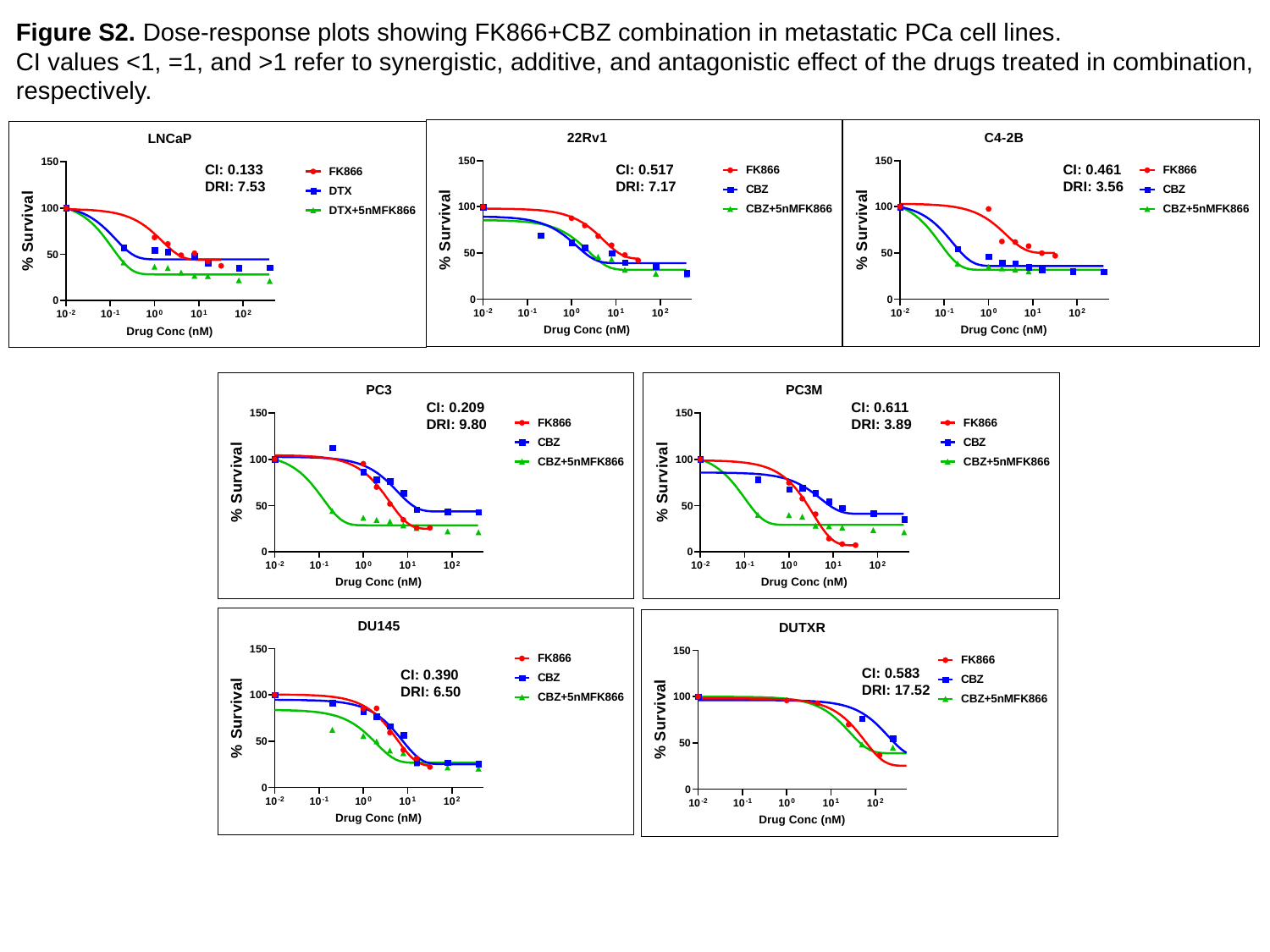

Figure S2. Dose-response plots showing FK866+CBZ combination in metastatic PCa cell lines.
CI values <1, =1, and >1 refer to synergistic, additive, and antagonistic effect of the drugs treated in combination, respectively.
CI: 0.133
DRI: 7.53
CI: 0.461
DRI: 3.56
CI: 0.517
DRI: 7.17
CI: 0.209
DRI: 9.80
CI: 0.611
DRI: 3.89
CI: 0.583
DRI: 17.52
CI: 0.390
DRI: 6.50

### Slide 3
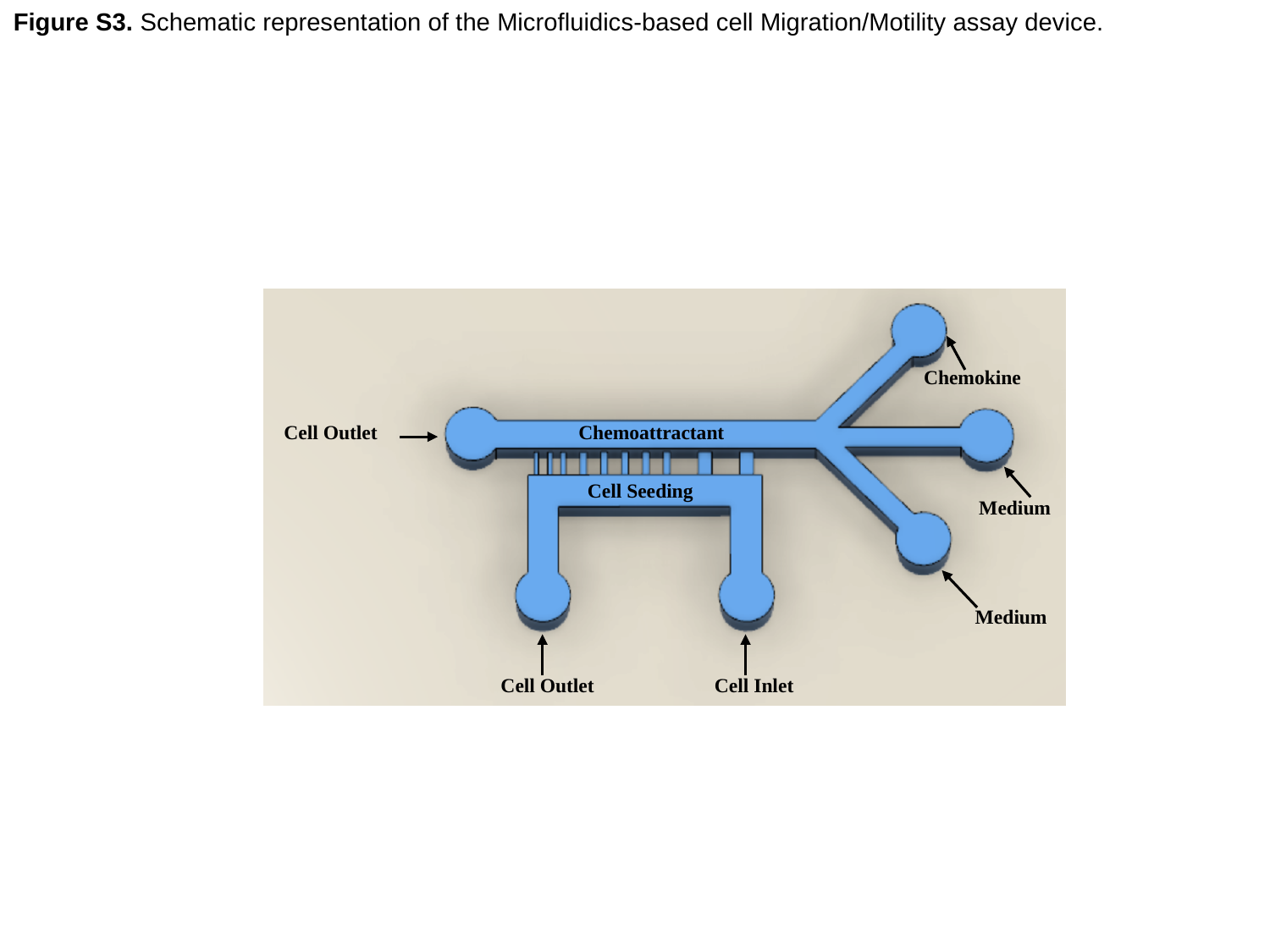

Figure S3. Schematic representation of the Microfluidics-based cell Migration/Motility assay device.
Chemokine
Cell Outlet
Chemoattractant
Cell Seeding
Medium
Medium
Cell Outlet
Cell Inlet

### Slide 4
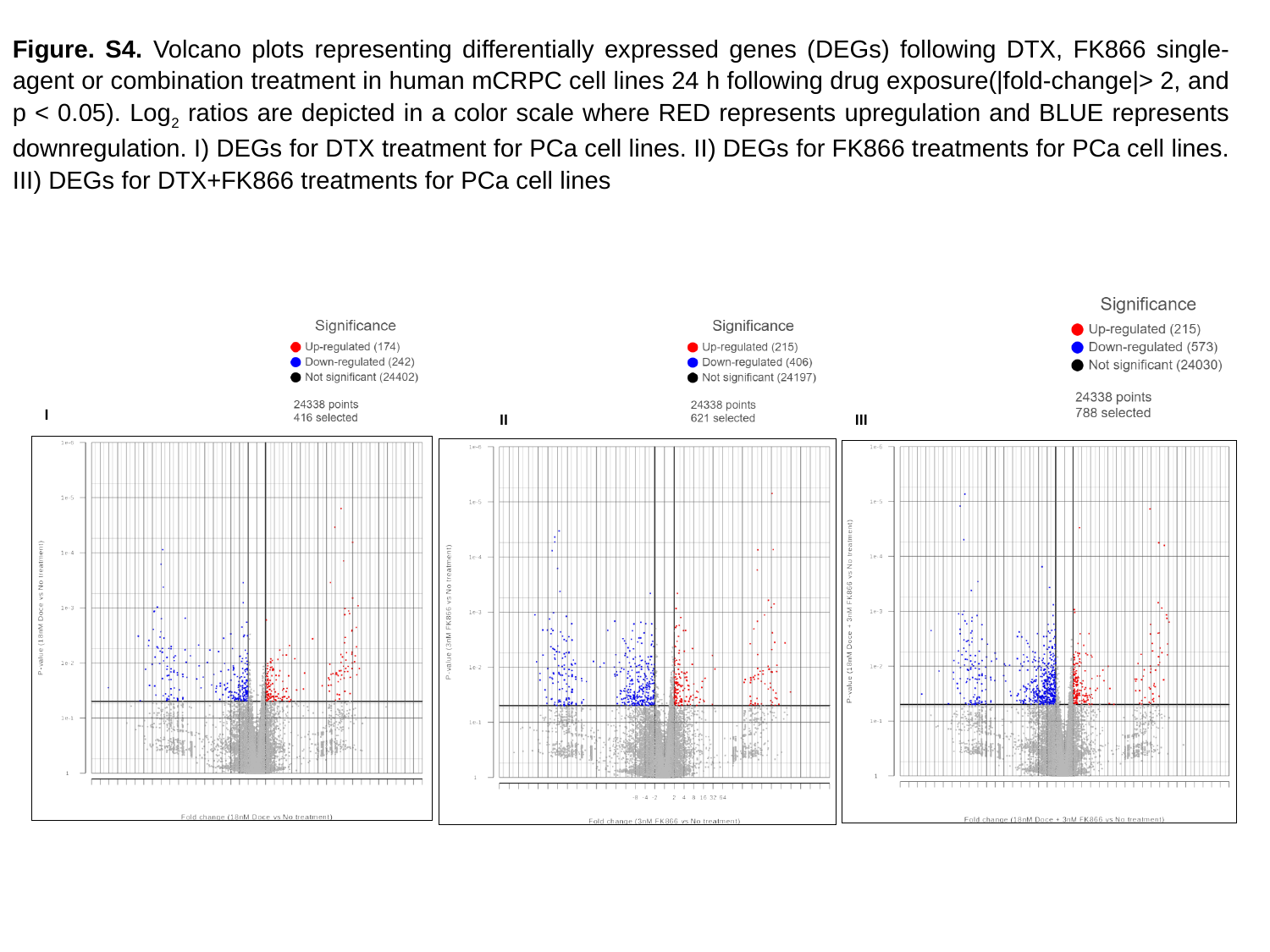

Figure. S4. Volcano plots representing differentially expressed genes (DEGs) following DTX, FK866 single-agent or combination treatment in human mCRPC cell lines 24 h following drug exposure(|fold-change|> 2, and p < 0.05). Log_2_ ratios are depicted in a color scale where RED represents upregulation and BLUE represents downregulation. I) DEGs for DTX treatment for PCa cell lines. II) DEGs for FK866 treatments for PCa cell lines. III) DEGs for DTX+FK866 treatments for PCa cell lines
Volcano Plots
I
II
III

### Slide 5
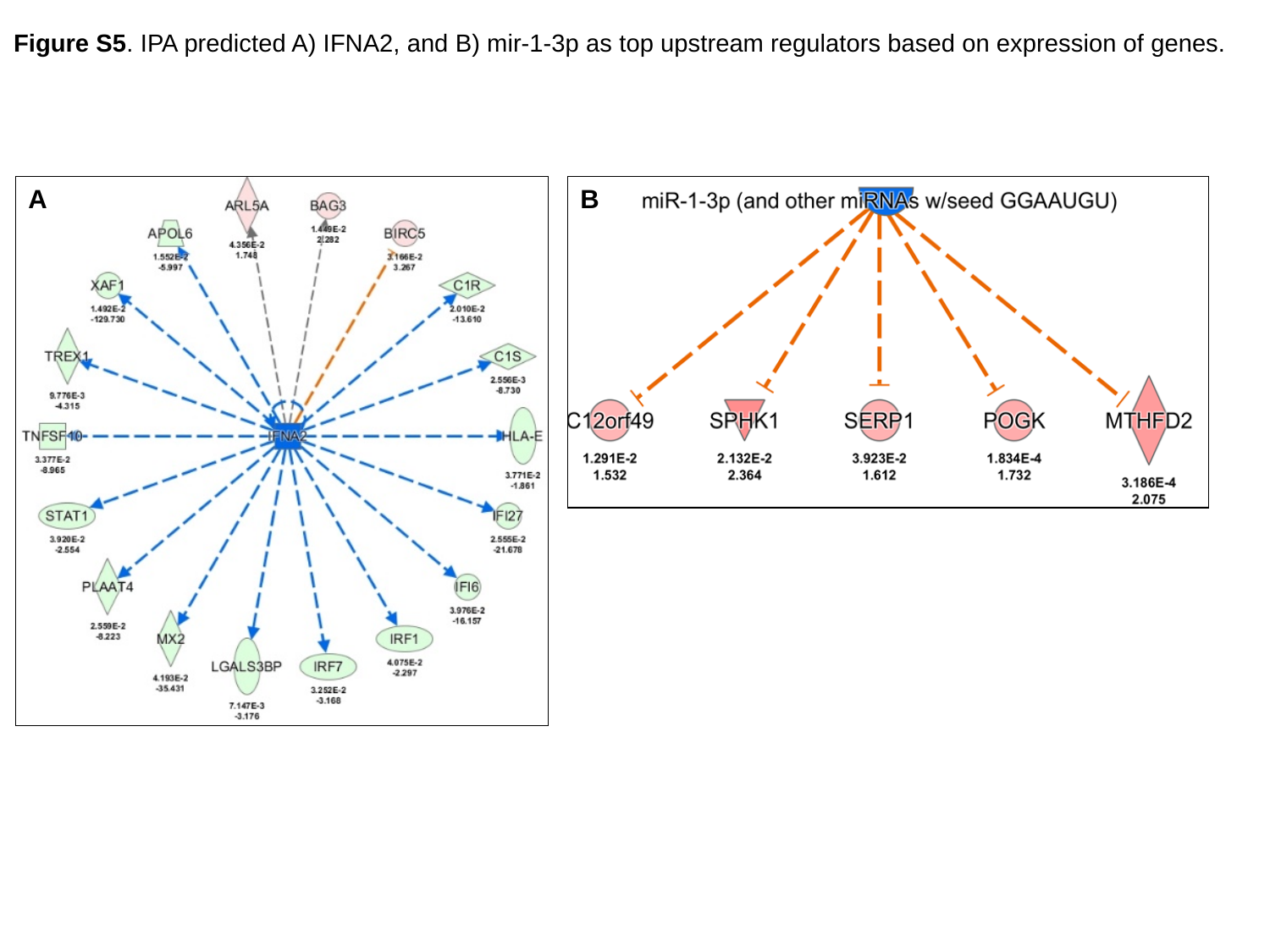

Figure S5. IPA predicted A) IFNA2, and B) mir-1-3p as top upstream regulators based on expression of genes.
A
B

### Slide 6
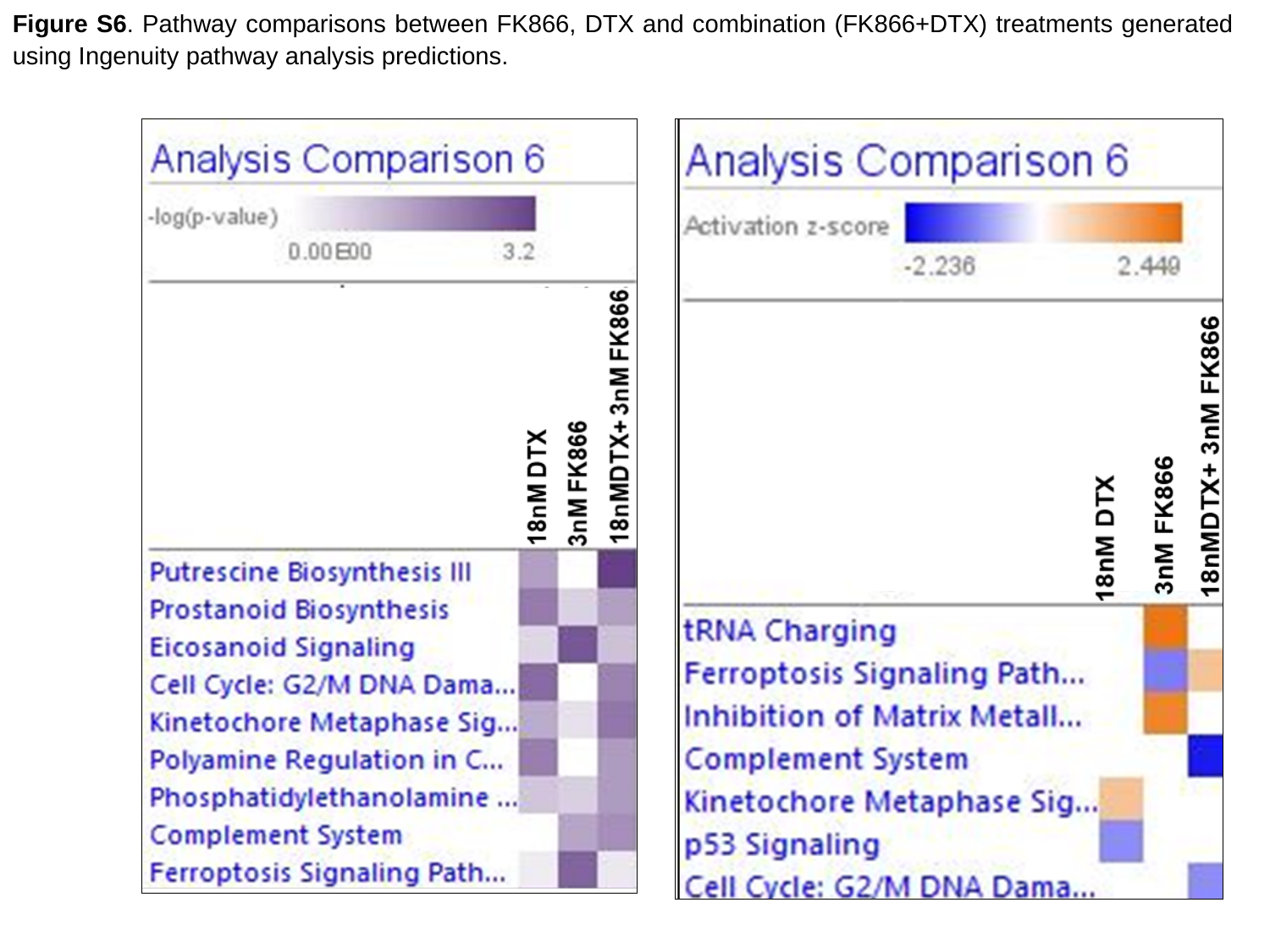

Figure S6. Pathway comparisons between FK866, DTX and combination (FK866+DTX) treatments generated using Ingenuity pathway analysis predictions.
