## Supplementary material for "Integrating pharmacogenomics data-driven prediction with bulk and single-cell RNAseq to demonstrate the efficacy of an NAMPT inhibitor against aggressive, taxane-resistant, and stem-like cells in lethal prostate cancer": Video

### Slide 1
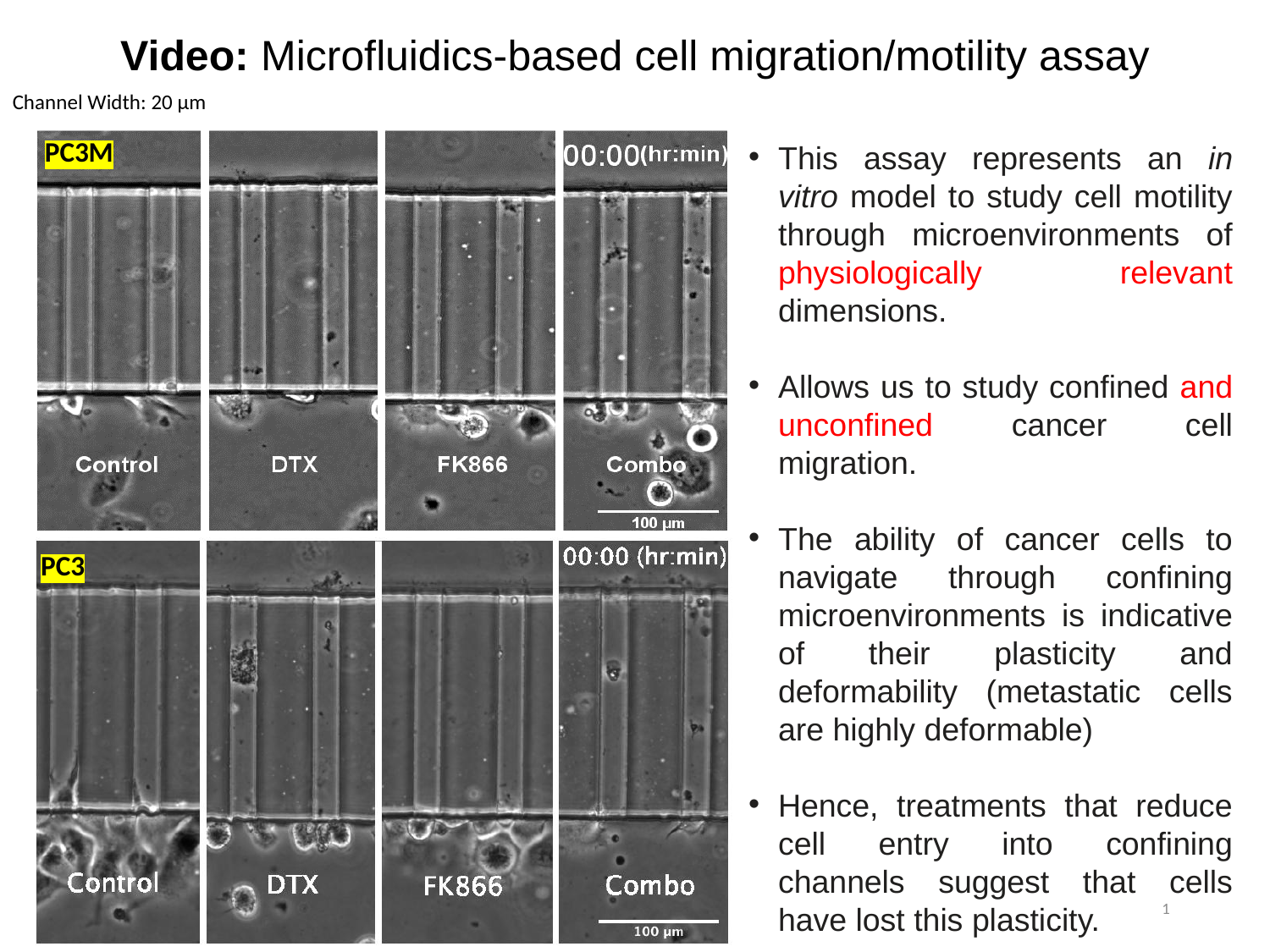

Video: Microfluidics-based cell migration/motility assay
Channel Width: 20 µm
PC3M
This assay represents an in vitro model to study cell motility through microenvironments of physiologically relevant dimensions.
Allows us to study confined and unconfined cancer cell migration.
The ability of cancer cells to navigate through confining microenvironments is indicative of their plasticity and deformability (metastatic cells are highly deformable)
Hence, treatments that reduce cell entry into confining channels suggest that cells have lost this plasticity.
PC3
1
