## Supplementary Tables for "Integrating pharmacogenomics data-driven prediction with bulk and single-cell RNAseq to demonstrate the efficacy of an NAMPT inhibitor against aggressive, taxane-resistant, and stem-like cells in lethal prostate cancer"

|  | **Drug Name** | **Target** | **Target Pathway** |
| --- | --- | --- | --- |
| 1 | **Afatinib** | ERBB2, EGFR | EGFR signaling |
| 2 | **AKT inhibitor VIII** | AKT1, AKT2, AKT3 | PI3K/AKT pathway |
| 3 | **AMG-706 (Motesanib)** | VEGFR, RET, KIT, PDGFR | RTK signaling |
| 4 | **AZD6482** | PI3Kβ | PI3K/MTOR signaling |
| 5 | **Cetuximab** | EGFR | EGFR signaling |
| 6 | **CP724714** | ERBB2 | RTK signaling |
| 7 | **FH535** | PPARγ, PPARδ | Wnt/β-catenin signaling |
| 8 | **FK866** | NAMPT | NAD+ salvage pathway |
| 9 | **GSK2126458 (Omipalisib)** | PI3K (class 1), MTORC1, MTORC2 | PI3K/MTOR signaling |
| 10 | **GW441756** | NTRK1 | RTK signaling |
| 11 | **KIN001-260** | IKKB | NF-κB pathway |
| 12 | **LY317615** | PKCB | Other, kinases |
| 13 | **MK-2206** | AKT1, AKT2 | PI3K/MTOR signaling |
| 14 | **Navitoclax** | BCL2, BCL-XL, BCL-W | Apoptosis regulation |
| 15 | **NSC-87877** | SHP-1 (PTPN6), SHP-2 (PTPN11) | Other |
| 16 | **PD-0325901** | MEK1, MEK2 | ERK MAPK signaling |
| 17 | **PD-173074** | FGFR1, FGFR2, FGFR3 | RTK signaling |
| 18 | **PI-103** | PI3Kα, DAPK3, CLK4, PIM3, HIPK2 | Other, kinases |
| 19 | **RDEA119** | MEK1, MEK2 | ERK MAPK signaling |
| 20 | **SNX-2112** | HSP90 | Protein stability and degradation |
| 21 | **TAK-715** | p38α, p38β | JNK and p38 signaling |
| 22 | **TL-2-105** | CRAF | ERK MAPK signaling |
| 23 | **WZ3105** | SRC, ROCK2, NTRK2, FLT3, IRAK1 | Other |
| 24 | **XAV939** | TNKS1, TNKS2 | WNT signaling |
| 25 | **YM155** | BIRC5 | Apoptosis regulation |

**Table S1.** Top drugs (‘secDrugs’) derived from our pharmacogenomics data-driven analysis

|  | **3nM FK866** | | **18nM DTX + 3nM FK866** | | **18nM DTX** | |
| --- | --- | --- | --- | --- | --- | --- |
| **Gene** | **P value** | **Fold change** | **P value** | **Fold change** | **P value** | **Fold change** |
| **C1S** | 6.59E-03 | -7.95 | NA | NA | NA | NA |
| **IFITM3** | 3.17E-02 | -6.90 | NA | NA | NA | NA |
| **FAM229A** | 7.60E-03 | -5.24 | 4.11E-02 | -3.22 | NA | NA |
| **LHFPL5** | 9.81E-03 | -5.17 | NA | NA | NA | NA |
| **DUS4L-BCAP29** | 2.80E-02 | -4.60 | NA | NA | NA | NA |
| **ZACN** | 2.35E-02 | -4.53 | NA | NA | NA | NA |
| **LGALS3BP** | 2.39E-03 | -4.28 | 7.15E-03 | -3.18 | NA | NA |
| **CXCL2** | 4.71E-02 | -3.93 | NA | NA | NA | NA |
| **HES1** | 1.62E-03 | -3.84 | 1.54E-04 | -5.96 | 2.24E-03 | -3.29 |
| **FAS** | 2.09E-02 | -3.60 | NA | NA | NA | NA |
| **LY6G5B** | 4.27E-02 | -3.58 | 3.17E-02 | -4.50 | NA | NA |
| **CDRT4** | 2.14E-02 | -3.53 | NA | NA | NA | NA |
| **MAT2A** | 3.58E-02 | -3.51 | 4.56E-02 | -3.23 | NA | NA |
| **ZC3H12A** | 6.56E-03 | -3.23 | NA | NA | NA | NA |
| **IL18BP** | 8.62E-03 | -3.16 | NA | NA | NA | NA |
| **PRSS53** | 1.11E-02 | -3.10 | NA | NA | NA | NA |
| **PILRB** | 2.08E-02 | -3.06 | NA | NA | NA | NA |
| **IGFLR1** | 3.73E-02 | -3.04 | NA | NA | NA | NA |
| **ABCA1** | 2.30E-02 | -2.87 | NA | NA | 2.44E-02 | -2.49 |
| **LTB4R** | 7.37E-03 | -2.79 | 5.60E-03 | -3.19 | NA | NA |
| **UBXN11** | 4.59E-04 | -2.75 | NA | NA | NA | NA |
| **NSUN5P2** | 6.42E-03 | -2.70 | 3.54E-03 | -3.22 | NA | NA |
| **RASL11A** | 4.66E-02 | -2.69 | NA | NA | NA | NA |
| **RNF207** | 2.01E-02 | -2.64 | NA | NA | NA | NA |
| **ZNF789** | 1.62E-02 | -2.61 | NA | NA | NA | NA |
| **TERT** | 3.50E-02 | -2.57 | NA | NA | NA | NA |
| **NAIP** | 6.45E-03 | -2.56 | NA | NA | NA | NA |
| **MOK** | 3.97E-03 | -2.54 | NA | NA | NA | NA |
| **BSCL2** | 9.00E-03 | -2.51 | NA | NA | NA | NA |
| **VAMP1** | 3.80E-03 | -2.49 | NA | NA | NA | NA |
| **PPIH** | 2.61E-03 | -2.48 | NA | NA | NA | NA |
| **STK26** | 1.62E-02 | 2.50 | NA | NA | NA | NA |
| **GABARAPL1** | 2.17E-02 | 2.50 | NA | NA | NA | NA |
| **NCR3LG1** | 1.92E-02 | 2.50 | NA | NA | NA | NA |
| **RB1CC1** | 4.59E-04 | 2.51 | NA | NA | NA | NA |
| **CPLANE2** | 1.91E-02 | 2.58 | NA | NA | NA | NA |
| **TRAPPC6B** | 4.88E-02 | 2.63 | NA | NA | NA | NA |
| **WDR31** | 8.77E-03 | 2.66 | NA | NA | NA | NA |
| **TBCK** | 2.76E-03 | 2.69 | NA | NA | NA | NA |
| **ASNS** | 5.57E-03 | 2.80 | NA | NA | NA | NA |
| **OTUB2** | 4.99E-03 | 2.87 | NA | NA | NA | NA |
| **OTUD1** | 2.10E-02 | 2.88 | NA | NA | NA | NA |
| **TVP23C-CDRT4** | 7.40E-03 | 2.90 | NA | NA | NA | NA |
| **SYNE3** | 1.50E-02 | 2.93 | NA | NA | NA | NA |
| **PITPNC1** | 8.92E-03 | 2.97 | 5.92E-03 | 3.29 | NA | NA |
| **ADM2** | 4.08E-02 | 2.98 | NA | NA | NA | NA |
| **MOCOS** | 3.40E-02 | 3.02 | NA | NA | NA | NA |
| **CHAC1** | 1.25E-03 | 3.18 | NA | NA | NA | NA |
| **JPH1** | 2.26E-02 | 3.26 | NA | NA | NA | NA |
| **ELMOD1** | 4.25E-02 | 3.35 | NA | NA | NA | NA |
| **SESN3** | 3.85E-02 | 3.40 | NA | NA | NA | NA |
| **CYP1A1** | 2.64E-02 | 3.72 | NA | NA | NA | NA |
| **DDIT4** | 2.17E-03 | 3.87 | NA | NA | NA | NA |
| **SLC7A11** | 5.16E-03 | 4.08 | NA | NA | NA | NA |
| **FTCDNL1** | 1.28E-02 | 4.12 | NA | NA | NA | NA |
| **LAMP3** | 4.16E-02 | 4.47 | NA | NA | NA | NA |
| **BHLHA15** | 2.14E-02 | 5.30 | NA | NA | NA | NA |
| **PAPPA2** | 4.78E-02 | 6.27 | NA | NA | NA | NA |
| **EML5** | 8.91E-03 | 7.23 | NA | NA | NA | NA |

**Table S2.** Top differentially expressed genes (DEGs) (fold change compared to untreated) following 48 h of Fk866, FK866+DTX and DTX treatment in PC-3, PC-3M and DU145 (AI-mCRPC/NEPC) cell lines (Fold change >2; p<0.05).

|  | **Patient Cohort (BCR=1 vs BCR=0)** | | | | **PCa Cells lines (FK866 treatment vs baseline)** | | | |
| --- | --- | --- | --- | --- | --- | --- | --- | --- |
| **Gene symbol** | **Fold change (BCR=1 vs. BCR=0)** | **LSMean (BCR=1)** | **LSMean (BCR=0)** | **P-value (BCR=1 vs. BCR=0)** | **Fold change (3nM FK866 vs. No treatment)** | **LSMean (3nM FK866)** | **LSMean (No treatment)** | **P-value (3nM FK866 vs. No treatment)** |
| **SLX1B-SULT1A4** | 1.74 | 1.52 | 0.88 | 5.72E-03 | -2.20 | 3.51 | 7.73 | 1.07E-02 |
| **IFITM3** | 1.73 | 13.19 | 7.64 | 1.45E-03 | -6.90 | 5.53 | 38.15 | 3.17E-02 |
| **TMEM80** | 1.37 | 3.64 | 2.66 | 6.27E-03 | -1.96 | 7.12 | 13.98 | 3.09E-02 |
| **NPIPB12** | 1.30 | 6.12 | 4.71 | 3.02E-02 | -1.58 | 9.77 | 15.48 | 4.19E-02 |
| **LTB4R** | 1.28 | 4.41 | 3.43 | 2.28E-03 | -2.79 | 5.90 | 16.48 | 7.37E-03 |
| **ZC3H12A** | 1.24 | 11.91 | 9.59 | 4.22E-02 | -3.23 | 3.34 | 10.78 | 6.56E-03 |
| **TMEM120B** | 1.23 | 12.73 | 10.35 | 3.95E-02 | -1.64 | 36.80 | 60.23 | 2.32E-02 |
| **POGK** | -1.11 | 11.05 | 12.26 | 3.01E-02 | 1.44 | 138.76 | 96.57 | 3.42E-02 |
| **RPS6KA5** | -1.14 | 9.74 | 11.10 | 4.43E-02 | 2.25 | 34.36 | 15.30 | 3.51E-02 |
| **ZNF770** | -1.20 | 14.74 | 17.66 | 4.43E-02 | 1.75 | 172.40 | 98.49 | 1.57E-02 |
| **AKAP11** | -1.21 | 30.41 | 36.90 | 3.32E-04 | 1.77 | 75.18 | 42.56 | 3.86E-02 |
| **PROSER1** | -1.22 | 10.82 | 13.22 | 3.90E-03 | 1.55 | 65.05 | 41.85 | 4.81E-02 |
| **FERMT2** | -1.23 | 11.70 | 14.39 | 4.20E-02 | 1.97 | 190.70 | 97.00 | 1.34E-02 |
| **SPRED1** | -1.23 | 9.71 | 11.97 | 4.59E-02 | 1.83 | 85.88 | 46.93 | 2.26E-02 |
| **EIF5** | -1.25 | 29.42 | 36.77 | 8.90E-04 | 1.72 | 829.98 | 482.60 | 2.91E-02 |
| **TRNT1** | -1.26 | 17.60 | 22.14 | 3.28E-02 | 1.74 | 27.99 | 16.10 | 1.87E-02 |
| **ARL5B** | -1.33 | 22.89 | 30.36 | 3.41E-03 | 1.77 | 46.36 | 26.20 | 3.67E-02 |
| **GABARAPL1** | -1.33 | 14.59 | 19.37 | 8.31E-05 | 2.50 | 27.86 | 11.15 | 2.17E-02 |
| **EML5** | -1.34 | 7.66 | 10.26 | 1.38E-02 | 7.23 | 41.73 | 5.77 | 8.91E-03 |
| **FBXL4** | -1.37 | 12.12 | 16.55 | 1.57E-02 | 1.61 | 43.43 | 27.03 | 3.74E-02 |

**Table S3.** List the genes that were significantly dysregulated in PCa patients with BCR AND showed significant differential expression in the opposite direction following FK866 treatment in our model systems, indicating that FK866 might be was capable of reversing the input signature in the patient cohort.

| **Reagents** | **Manufacturer** | **Location** |
| --- | --- | --- |
| Fetal bovine serum (FBS) | Hyclone (Thermo Fisher Scientific Inc.) | Rockford, IL, USA |
| Trypsin (0.25% w/v) | Hyclone (Thermo Fisher Scientific Inc.) | Rockford, IL, USA |
| Penicillin-Streptomycin (10,000 U/mL) | Gibco^TM^ (Thermo Fisher Scientific Inc.) | Waltham, MA, USA |
| Docetaxel | Selleck Chemicals LLC | Houston, TX, USA |
| Cabazitaxel | Selleck Chemicals LLC | Houston, TX, USA |
| FK866 | Selleck Chemicals LLC | Houston, TX, USA |
| FITC Annexin V Apoptosis Detection Kit | BD Biosciences | San Jose, CA, USA |
| FxCycle™ PI/RNase Staining Solution kit | Thermo Fisher Scientific Inc. | Waltham, MA, USA |
| RIPA lysis buffer | Thermo Fisher Scientific Inc. | Waltham, MA, USA |
| Protease Inhibitor Cocktail | Thermo Fisher Scientific Inc. | Waltham, MA, USA |
| Phosphatase Inhibitor | Thermo Fisher Scientific Inc. | Waltham, MA, USA |
| Pierce™ ECL Western Blotting Substrate | Thermo Fisher Scientific Inc. | Waltham, MA, USA |
| HES-1 specific primer Hs00172878_m1 | Thermo Fisher Scientific Inc. | Waltham, MA, USA |
| TaqMan Gene Expression Assays | Thermo Fisher Scientific Inc. | Waltham, MA, USA |
| Vybrant™ DyeCycle™ Violet Stain | Thermo Fisher Scientific Inc. | Waltham, MA, USA |
| CM-H2DCFDA (General Oxidative Stress Indicator) | Thermo Fisher Scientific Inc. | Waltham, MA, USA |
| NucBlue™ Live ReadyProbes™ Reagent (Hoechst 33342) | Thermo Fisher Scientific Inc. | Waltham, MA, USA |
| RNeasy Plus Mini Kit | QIAGEN | Hilden, Germany |
| QuantiTect Reverse Transcription Kit | QIAGEN | Hilden, Germany |
| Quick Start Bovine Serum Albumin Standard | Bio-Rad | Hercules, CA, USA |
| Tris Buffer Saline (TBS) | Bio-Rad | Hercules, CA, USA |
| 10% Tween 20 | Bio-Rad | Hercules, CA, USA |
| Polyvinylidene fluoride membrane (PVDF) | EMD Millipore | Billerica, MA, USA |
| Bovine Serum Albumin (BSA) | VWR | Radnor, PA, USA |
| 3-(4,5-dimethylthiazol-2-yl)-2,5-diphenyltetrazolium bromide (MTT) | Sigma-Aldrich Inc | St. Louis, MO, USA |
| Dimethyl sulfoxide (DMSO) | Sigma-Aldrich Inc | St. Louis, MO, USA |
| 2′,7′-Dichlorofluorescin diacetate (DCFDA) | Sigma-Aldrich Inc | St. Louis, MO, USA |
| Bradford Reagent | Sigma-Aldrich Inc | St. Louis, MO, USA |
| NAD/NADH-Glo™ Assay Kit (NAMPT activity) | Promega Corp. | Madison, WI, USA |
| JC-1 - Mitochondrial Membrane Potential Assay Kit | Abcam | Waltham, MA, USA |
| NAMPT (86634S) | Cell Signaling Technology | Danvers, MA, USA |
| Becline-1 (3495S) | Cell Signaling Technology | Danvers, MA, USA |
| HES1 (11988S) | Cell Signaling Technology | Danvers, MA, USA |
| CD44 (3570T) | Cell Signaling Technology | Danvers, MA, USA |
| Cleave-caspase 9 (9505T) | Cell Signaling Technology | Danvers, MA, USA |
| Cleave-caspase 3 (9661T) | Cell Signaling Technology | Danvers, MA, USA |
| ATF-4 (11815S) | Cell Signaling Technology | Danvers, MA, USA |
| Anti-rabbit IgG, HRP-linked Antibody (7074S) | Cell Signaling Technology | Danvers, MA, USA |
| β-actin (A3854) | Sigma-Aldrich Inc | St. Louis, MO, USA |

**Supplementary Table S4.** List of Drugs, reagents, antibodies, and kits.
